# Improved short nascent strand sequencing (iSNS-seq) enhances DNA replication origin detection and reduces non-origin biases

**DOI:** 10.64898/2025.12.22.696044

**Authors:** Miiko Sokka, John M. Urban, Nicola Neretti, Susan A. Gerbi

## Abstract

Identifying DNA replication origins in human and other metazoan genomes has been challenging, as highlighted by the fact that various methods for mapping them have produced conflicting results. A popular method, short nascent strand sequencing (SNS-seq), enriches newly replicated short single-stranded DNA by size selection and *λ*-exonuclease (*λ*-exo) digestion of parental DNA. Surprisingly, SNS-seq has never been validated in *Saccharomyces cerevisiae* where origins have been well characterized genome-wide. Here we improved the SNS-seq protocol through biochemical optimization and benchmarked its origin-mapping sensitivity and precision against traditional SNS-seq in asynchronous populations of *S. cerevisiae*, a genetically tractable system that allows direct comparison against a well-defined set of confirmed origins. Relative to traditional SNS-seq, the improved protocol substantially increased enrichment of origin-derived DNA. Strikingly, traditional SNS-seq failed to detect known origins and instead enriched non-origin DNA, likely arising from RNA:DNA hybrids. These findings have important implications for the interpretation of previously published datasets that rely on *λ*-exo for origin mapping, and provide a proof-of-concept benchmark for extending this improved protocol to metazoan systems. Furthermore, our biochemical and genomic analyses help unravel the mystery of the inconsistencies between SNS-seq and other techniques used to map DNA replication origins genome-wide.

## Introduction

Initiation of DNA replication requires stepwise DNA binding and the accumulation of several factors that ultimately lead to the firing of replication forks and to bidirectional DNA synthesis (1). Origins of replication initiation (hereafter referred to simply as *origins*, used to denote replication fork initiation sites) are genomic sites where this process occurs; eukaryotic chromosomes contain numerous origins distributed across their length (2).

In metazoan cells, accumulating evidence indicates plasticity in origin specification, and they might be better categorized as initiation zones with preferred sites of initiation (3; 4; 5; 6). However, whether discrete origins exist in metazoans or origin selection occurs stochastically within broad initiation zones remains a matter of debate. Consequently, mapping origins within metazoan genomes remains an area of active research, even after several decades where multiple approaches have been developed to map replication origin locations genome-wide (7; 8; 9).

The challenge of reliably mapping metazoan replication origins stems from the infrequency of site-specific initiation events, resulting in a weak signal to noise ratio in origin mapping datasets. Moreover, any off-target bias present in a technique may be far more common than site-specific initiation events and result in systematic false positives. Therefore, it is crucially important to understand the technical principles underlying each origin mapping method to critically evaluate biological conclusions of the resulting origin maps.

One of the most widely applied methods for identifying preferred initiation sites is short nascent strand sequencing (SNS-seq). Among the many approaches to map origins, SNS-seq stands out due to its relative simplicity and practicality: it can be applied to any cell type without experimental manipulations such as synchronization, metabolic labeling, or genetic engineering. However, the poor concordance between SNS-seq and other origin mapping methods has contributed significantly to the ongoing debate over whether metazoan origins are discrete or dispersed (7; 8; 9; 10). On one side are techniques that have suggested discrete initiation sites, such as SNS-seq (11; 12; 13; 14) and INI-seq (15; 16), the latter of which does not rely on *λ*-exo digestion. On the opposing side are methods such as Bubble-seq (17), OK-seq (4; 18), nanopore-based mapping using incorporated BrdU signal (19; 20) and optical replication mapping (5) that provide evidence of broad initiation zones with more dispersed initiation sites. It has recently been argued that these apparently contradictory results may be reconciled through models incorporating fork stalling, unidirectional initiation, and the aggregation of inefficient sites into larger zones (10). Such reconciliation frameworks, however, presuppose that the discrete signals detected by initiation-mapping techniques like SNS-seq represent bona fide replication intermediates rather than systematic technical artifacts. This assumption has rarely been directly tested in a system where origin locations are known with confidence. Indeed, it has been previously noted that SNS-seq has not been applied to budding yeast, where its sensitivity and precision could be benchmarked against well-characterized origins (9).

SNS-seq enriches 500–2,000 nucleotide nascent single-stranded DNA in two major steps (21). Heat-denatured genomic DNA is size-fractionated on a sucrose gradient, yielding a mixture of nascent strands and excess parental DNA. The size-selected sample is then treated with *λ*-exonuclease (*λ*-exo), which digests DNA but not RNA from the 5’ end (22). Because nascent strands carry a 5’ RNA primer derived from DNA primase/polymerase *α*, they are thought to be protected from digestion while parental DNA is degraded. Multiple rounds of *λ*-exo treatment are typically performed to ensure efficient background removal (23; 24).

Using SNS-seq to map replication origins therefore also crucially depends on the assumption that RNA primers serve as the most substantial or the sole factor that inhibits DNA digestion by *λ*-exo, resulting in enrichment of short nascent strands. However, it has been shown that *λ*-exo non-uniformly enriches DNA sequences independent of nascent DNA, and that digestion is impeded in GC-rich DNA and by specific GC-rich motifs including G4 quadruplexes (25; 26; 27; 28). Thus, it is possible that the discrete sites mapped with SNS-seq reflect other phenomena or arise from a mixture of sources that inhibit enzymatic digestion, some of which remain unknown, in addition to nascent strands. An open question is if discrete origins existed and their locations were known, how well would SNS-seq perform in enriching them? One system where this question can be addressed is budding yeast (*Saccharomyces cerevisiae*), which has discrete origins and a rich origin-mapping history (29). Any new origin mapping method should minimally be characterized by its sensitivity in finding the known origins in budding yeast, and by its precision against enriching non-origin sites. Intriguingly, SNS-seq has never been tested for its ability to map origins within the budding yeast genome. In this work, we set out to determine the sensitivity and precision of SNS-seq in budding yeast, and to test improved conditions suggested by our *in vitro* experiments. Our results reveal a novel bias and show that traditional SNS-seq enriches non-origin DNA that appears to originate from RNA:DNA hybrids due to residual cellular RNA, while failing to enrich known replication origins above random expectation. Our data provide a concrete explanation for the longstanding discrepancies between traditional SNS-seq results and results from orthogonal origin-mapping methods. Our improved conditions address the major known biases of *λ*-exo and enable detection of known origins in budding yeast.

## Materials and Methods

### Oligo assays

All single-stranded oligomeric DNAs were ordered from Integrated DNA Technologies (Coralville, Iowa) with sequences shown in Table 1. For each experiment, the oligos were heated for 1 min at 95°C to make sure they are single-stranded, then treated with 10 U of T4 polynucleotide kinase (PNK) (New England Biolabs, Ipswich, MA, catalog Nr. M0201), purified with Qiaquick spin nucleotide removal kit (Qiagen, Germantown, MD, catalog Nr. 28306). The purified oligos were once again heated for 1 min at 95°C to make sure they are single-stranded, and then subjected to 10 U of *λ*-exo (New England Biolabs, Ipswich, MA, catalog Nr. M0262) digestion for the indicated times in either glycine-KOH buffer (67 mM glycine-KOH pH 9.4, 2.5 mM MgCl_2_, 0.01% (v/v) Triton X-100) or Tris-HCl (20 mM Tris-HCl pH 9.0, 1 mM MgCl_2_, 0.01% v/v Triton X-100) buffer, before ending the reaction by denaturing the *λ*-exo protein for 15 min at 75 °C. An aliquot of the DNA sample was run in 15% PAGE (polyacrylamide gel electrophoresis) and stained with SYBR Gold (Thermo Fisher Scientific, Waltham, MA, catalog nr. S11494). Equal molar amounts of oligos were used in experiments where multiple oligos were mixed in the same reaction. The lower detection limit was about 5 ng of chimeric RNA–DNA oligo using 15% PAGE and SYBR Gold.

**Table 1.**
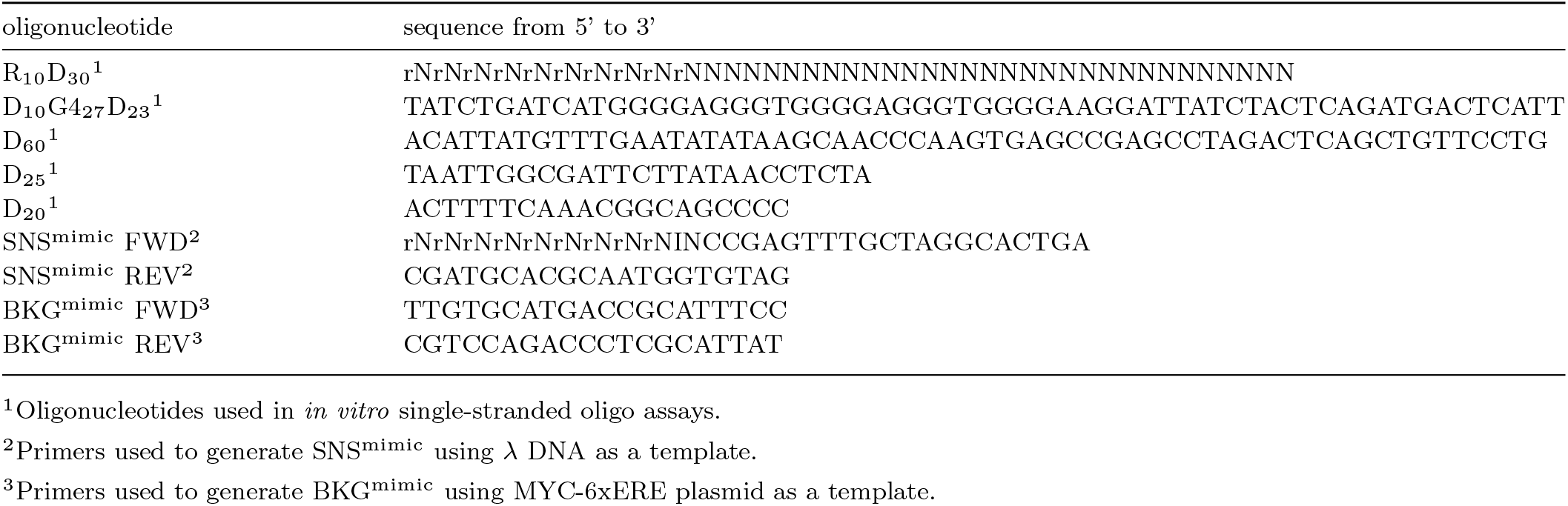
The oligonucleotide sequences used in this study.

A note on reproducibility of gel-based digestion assays: Buffer conditions and additives for the *λ*-exo and RecJf digestion assays (Figures 1B, C, E, and 2D) were established through an extensive experimental search aimed at identifying conditions that increase digestion through G4 motifs while preserving the RNA-DNA chimera. Across this large number of experiments the same core observations consistently emerged, including that *λ*-exo digests through the RNA-DNA chimera, that *λ*-exo digestion of G4 motifs is impeded in Glycine-KOH buffer but not in Tris-HCl, that a 5’-OH end protects DNA from *λ*-exo digestion more effectively than an RNA primer, and that RecJf does not digest through a 5’ RNA end but reduces background DNA levels. The gel panels shown are representative of these consistently observed outcomes.

**Fig. 1:**
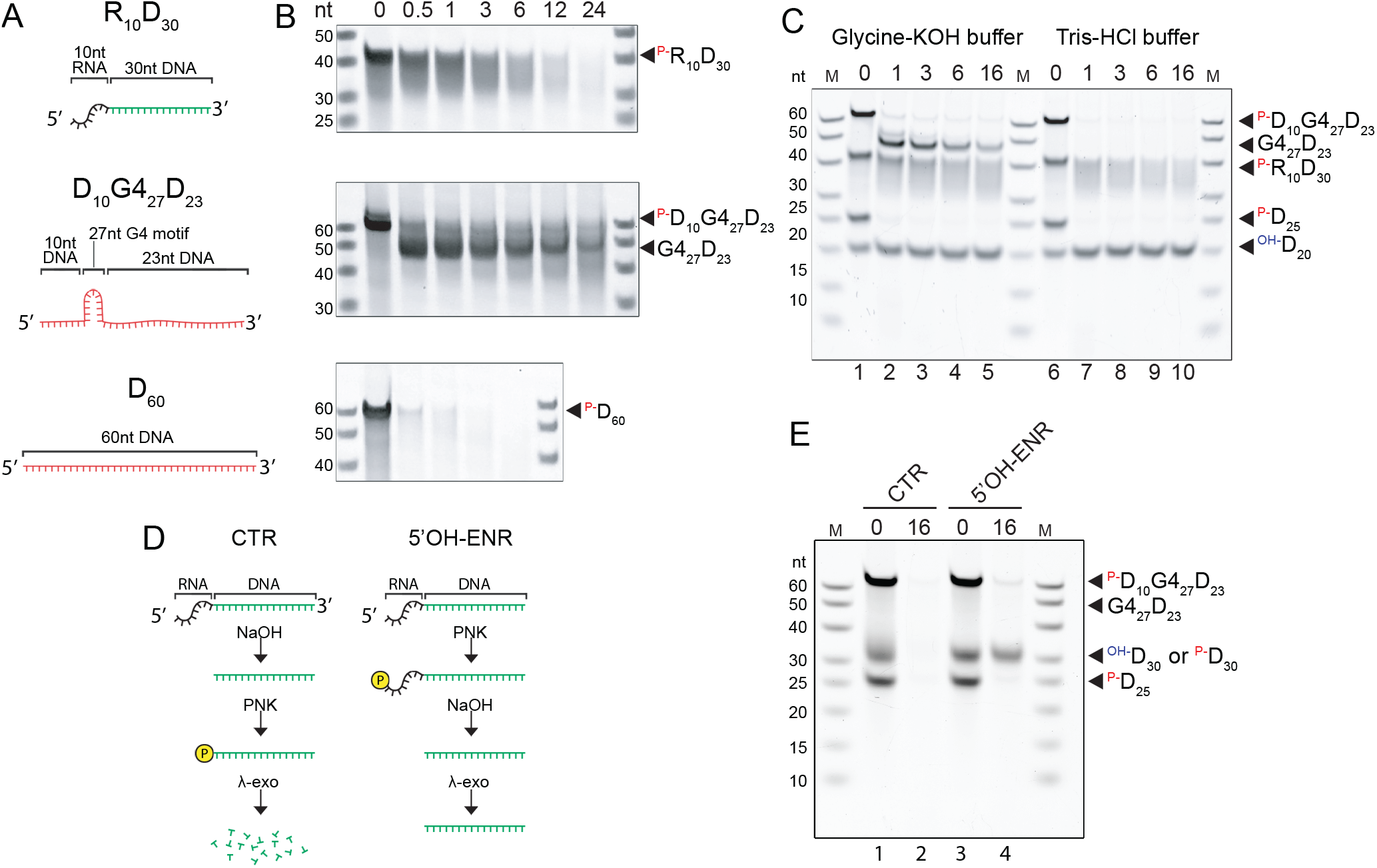
*λ*-exo digests through 5’ RNA ends but is blocked by a G4 motif in glycine-KOH buffer, an effect not observed with 5’ OH ends in Tris-HCl buffer. (A) Schematic of the oligomeric nucleic acid constructs. (B) Urea-PAGE gels showing a time course of *λ*-exo digestion for indicated times in hours in glycine-KOH buffer (67 mM glycine-KOH pH 9.4, 2.5 mM MgCl_2_, 0.01% (v/v) Triton X-100). (C) A time course of all oligos used in B combined in the same reaction and treated with either glycine-KOH buffer as above or Tris-HCl buffer (20 mM Tris-HCl pH 9.0, 1 mM MgCl_2_, 0.01% v/v Triton X-100) for the indicated times in hours. (D) Schematic of the experimental steps to prepare samples for panel E. The RNA-primed chimera R_10_D_30_ is depleted in the control sample (CTR) because NaOH is used to hydrolyze the RNA before adding the phosphate necessary for *λ*-exo. In contrast, the RNA-primed DNA chimera is preserved in the enrichment sample (ENR), because hydrolysis of the RNA also removes the phosphate added before the NaOH step. (E) Using the experimental setup shown in D the same oligos as above (except that the oligo labeled in panel D as OH-D_30_ is the product remaining after NaOH hydrolysis of ^P-^R_10_D_30_) were subjected to *λ*-exo for 0 or 16 hours as indicated. In all experiments, each oligo was used in equal molarity of 30 pmol. Each gel panel (B, C, E) is representative of results consistently observed across repeated independent experiments (see Methods).

### Artificial short nascent strand DNA experiments

To prepare 1.2 kb RNA-primed DNA (SNS^mimic^), we used Q5 High Fidelity PCR (New England Biolabs, Ipswich, MA, catalog Nr. E0555) with chimeric RNA-DNA primers (for the primer sequences, see Table 1) and Lambda DNA (New England Biolabs, Ipswich, MA, catalog Nr. N3011) as a template (Supplementary Figure S1). We first optimized the PCR cycling conditions and verified that the RNA primers are preserved during the high temperatures required for PCR cycling (Supplementary Figure S2). For the 1.2 kb control (BKG^mimic^) we used primers listed in Table 1 and a pFRT.myc6xERE plasmid (28) as a template to amplify a region with multiple G4 motifs. All amplified PCR products were purified using 1x AMPure XP beads (Beckman Coulter Life Sciences, Indianapolis, IN, catalog Nr. A63881).

To prepare samples for the *λ*-exo reaction, 15 ng (20 fmol) of each 1.2 kb SNS^mimic^ and BKG^mimic^ were mixed with a polyacryl carrier (Molecular Research Center, Cincinnati, OH, catalog Nr. PC-152) (to help subsequent recovery of the DNA) and denatured for 1 min at 95 °C. The DNAs were directly subjected to T4 PNK (New England Biolabs, Ipswich, MA, catalog Nr. M0201) in 700 mM Tris-HCl pH 7.6, 100 mM MgCl_2_, 50 mM DTT followed by PNK inactivation by adding 1 *µ*l of 0.5 M EDTA and incubating at 65 °C and then purified with 1x AMPure XP beads according to the manufacturer’s instructions. For samples that were subjected to RNA hydrolysis, we treated the sample with 280 mM NaOH for 15 min at 50 °C, neutralized by adding 280 mM acetic acid and purified with 1x AMPure XP beads. For *λ*-exo digestion, the DNA sample was denatured for 1 min at 95 °C and then subjected to 1,000 U of *λ*-exo (New England Biolabs, Ipswich, MA, catalog Nr. M0262) per 1 *µ*g of DNA either in glycine-KOH (67 mM glycine-KOH pH 9.4, 2.5 mM MgCl_2_, 0.01% (v/v) Triton X-100) or Tris-HCl (20 mM Tris-HCl pH 9.0, 1 mM MgCl_2_, 0.01% v/v Triton X-100) reaction buffer. The input (0 h) sample was taken before adding *λ*-exo, and each timepoint sample was immediately subjected to a 15 min treatment at 75 °C to inactivate *λ*-exo enzymatic activity.

For the samples treated with RecJf, the DNA was subjected to 16 h treatment in 20 U of RecJf (New England Biolabs, Ipswich, MA, catalog Nr. M0264) in the presence of 20 U of SUPERase.In RNase inhibitor (Thermo Fisher, Waltham, MA, catalog Nr. AM2694) after denaturation as above and before PNK and NaOH treatments. The DNA that remained after RecJf digestion was purified using AMPure XP beads as described above.

### Genomic DNA extraction

For each replicate, we collected 1.5 liters of logarithmic phase culture of W303 strain *Saccharomyces cerevisiae* (a gift from Joachim Li) grown in standard YPAD media (1% Yeast Extract, 2% Peptone, 40 *µ*g/ml Adenine Sulfate, 2% Dextrose) by mixing cells first in water with 0.1% Na-azide and then quickly cooling the solution by mixing the cell suspension in a centrifuge bottle with a frozen (−70 °C) pellet of 133 mM EDTA, 33% glycerol (approximately 3:20 ratio of EDTA-glycerol and cell suspension). To extract genomic DNA, yeast cells were first “softened” by incubating the cell suspension in 100 mM *β*-mercaptoethanol in 10 mM Tris-HCl pH 8.0 on ice for 10 min, then washed once with cold water. Each approximately 0.5 ml cell pellet was then suspended in 0.9 ml of SCE buffer (800 mM sorbitol, 100 mM *β*-mercaptoethanol, 80 mM Na-citrate, 8 mM EDTA supplemented with SUPERase.In RNase inhibitor) and 0.1 ml of Zymolyase 100T (US Biologicals, Salem, MA, catalog Nr. Z1005) and incubated with rotation for 30 min at 37 °C to break the cell wall. The resulting spheroplasts were washed three times in ice cold SCE buffer and then lysed in DNAzol (Thermo Fisher Scientific, Waltham, MA, catalog Nr. 10503-027) supplemented with 0.2 mg/ml proteinase K (Qiagen, Germantown, MD, catalog Nr. 19131) and incubated at room temperature overnight. After pelleting the debris by centrifugation for 10 min at 10,000 x g, the DNA was precipitated from the supernatant using ethanol and redissolved in TEN20S (10 mM Tris-HCl pH 8.0, 2 mM EDTA, 20 mM NaCl, 0.1% SDS). Standard phenol-chloroform and chloroform extractions were done for further purification. Finally, the genomic DNA was precipitated with standard isopropanol precipitation using 0.2 M NaCl, washed twice with 80% ethanol and redissolved in 10 mM Tris, 1 mM EDTA, pH 8.0.

### Genomic DNA size selection

Before DNA size selection in a sucrose gradient, the sample was spiked-in with 0.5 *µ*l of an equal molar mixture of 1.2 kb SNS^mimic^ and BKG^mimic^ DNA (see their sequences in the Supplementary Table S1). The sample was then denatured by heating at 95 °C for 10 min, quickly cooled on ice and then gently applied on top of a 5–30% linear sucrose gradient prepared in 50 mM Tris-HCl pH 8.0, 50 mM EDTA, 500 mM NaCl. The DNA fragments were separated by centrifuging 11 h at 4°C at 35,000 rpm in a SW-41 swinging bucket rotor. Then 0.5 ml fractions were collected from the top of the tube using a wide-bore P1000 pipette tip. The fractions containing 500 nt–1,000 nt (determined by gel electrophoresis) DNA were pooled and concentrated by Amicon Ultra 10K (MilliporeSigma, catalog Nr. UFC901024, Burlington, MA) filtration, then precipitated with 2 volumes ethanol and eluted in ultra pure water. At this point, the samples were equally split into two aliquots to be processed using either the traditional (tSNS-seq) or the improved (iSNS-seq) protocol.

### Enrichment of short nascent strands using *λ*-exo

The traditional SNS-seq (tSNS) protocol involved two rounds of *λ*-exo digestion. For each round, the sample was heated for 3 min at 95 °C, cooled on ice and treated with 10 U of T4 PNK for 30 min at 37 °C, purified with 1x AMPure XP beads and treated in 200 U of *λ*-exo (50 U/*µ*l custom highly concentrated preparation, New England Biolabs, Ipswich, MA) in the presence of 20 U of SUPERase.In in glycine-KOH (67 mM glycine-KOH pH 9.4, 2.5 mM MgCl_2_, 0.01% (v/v) Triton X-100) for 16 h at 37 °C followed by inactivation of the enzyme for 15 min at 75 °C, and purification with 1x AMPure XP beads. Another round of T4 PNK and *λ*-exo treatment was performed (same conditions as the first round) before the RNA was hydrolyzed in a final solution of 280 mM NaOH (pH 10) for 15 min at 50 °C and then neutralized with an equal molar amount of acetic acid.

The improved SNS-seq (iSNS) protocol was performed as follows. The sample was heated for 3 min at 95 °C, cooled on ice and subjected to 300 U of RecJf (New England Biolabs, Ipswich, MA, catalog Nr. M0264) in the presence of 20 U of SUPERaseIn in 50 mM NaCl, 10 mM Tris-HCl (pH 7.9), 10 mM MgCl_2_, 1 mM DTT for 16 h at 37 °C, purified using 1x AMPure XP beads, treated with 10 U of T4 PNK for 30 min at 37°C, and purified with 1x AMPure XP beads. The RNA was hydrolyzed in a final 280 mM NaOH solution (pH 10) for 15 min at 50 °C and then neutralized with equal molarity acetic acid, followed by a single round of treatment in 200 U of *λ*-exo (50 U/*µ*l custom highly concentrated preparation, New England Biolabs, Ipswich, MA) in Tris-HCl buffer (20 mM Tris-HCl pH 9.0, 1 mM MgCl_2_, 0.01% v/v Triton X-100) for 16 h at 37 °C.

Both tSNS-seq and iSNS-seq samples were made double-stranded by first denaturing the DNA for 3 min at 95 °C and then incubating in the presence of 200 nM random hexamers (Integrated DNA Technologies, Coralville, Iowa) and 5 U of DNA Polymerase I Large (Klenow) fragment (New England Biolabs, Ipswich, MA, catalog Nr. M0212S) in 50 mM NaCl, 10 mM Tris-HCl (pH 7.9), 10 mM MgCl_2_, 1 mM DTT, 100 *µ*M dNTP. The polymerase was inactivated by adding EDTA to a final 10 mM and heating for 20 min at 75 °C. The quality of SNS enrichment was determined using qPCR for the spiked-in SNS^mimic^ and BKG^mimic^.

### Sequencing

The samples were prepared for sequencing using a NEBNext Ultra II library preparation kit (New England Biolabs, Ipswich, MA, catalog Nr. E7645). Briefly, the DNA was fragmented to an average length of 300 bp using a Covaris S220, end-repaired and dA-tailed and ligated with NEBNext adaptors (New England Biolabs, Ipswich, MA, catalog Nrs. E7335, E7500, E7710, E7654). The appropriate number of amplification cycles was determined for each sample by qPCR to reduce over-amplification. For this purpose, an aliquot of the ligated sample was mixed with NEBNext Universal PCR primer and the appropriate NEBNext Index Primer and SYBR Green Master Mix and run in QuantStudio 6 Pro Real-Time PCR (Applied Biosystems). The cycle number that corresponded to 50% of the maximum amplification minus three cycles (to accommodate 10% of material used in the test) was determined as the best library amplification cycle number. The library fragment size was determined with an Agilent Fragment Analyzer and each library was quantified by qPCR before pooling equal molar amounts of each library for indexed sequencing. Sequencing was performed using the NovaSeq 6000 (Novogene) with 150 bp paired-end sequencing methodology.

### Sequence data analysis

The fastq files were aligned to a sacCer3 reference genome where the spike-ins SNS^mimic^ and BKG^mimic^ sequences were added as extra “chromosomes” using Bowtie 2.5.3 (30; 31) (with the following parameters: –maxins 1000, –local, –dovetail, –soft-clipped-unmapped-tlen, -D 500, -R 3, -N 0, -L 19, -i C,1,0, –score-min C,40,0). Samtools 1.22.1 (32) was used to mark duplicate reads and then to remove unmapped, duplicate, low quality and orphan reads; and ultimately to keep only concordantly mapped paired reads (with parameters -F1028 -f3 -q2).

Peaks were called from the BAM file for each replicate sample using MACS 3.0.3 (33) callpeak function with a q-value cutoff of 0.05 without using a control file (-f BAMPE –mfold 2 100, -q 0.05). The fragment pileup bedgraph obtained with the MACS option –bdg, was used as the coverage track throughout the analyses. The bedgraphs were converted to bigwigs using UCSC bedGraphToBigWig 2.10 tool and visualized in Integrative Genomics Viewer (IGV) (34).

For comparison of different origin-mapping methods, the following published datasets were used: FORK-seq (20), OK-seq (35) and ORC ChIP-seq (36) (Table 2). ORC ChIP-seq reads (NCBI SRA Accession SRR1261246, SRR1261247) were mapped and processed as above, and peaks were called using MACS 3.0.2 (with parameters -mfold 2 100, –keep-dup 1, -q 0.001).

**Table 2.**
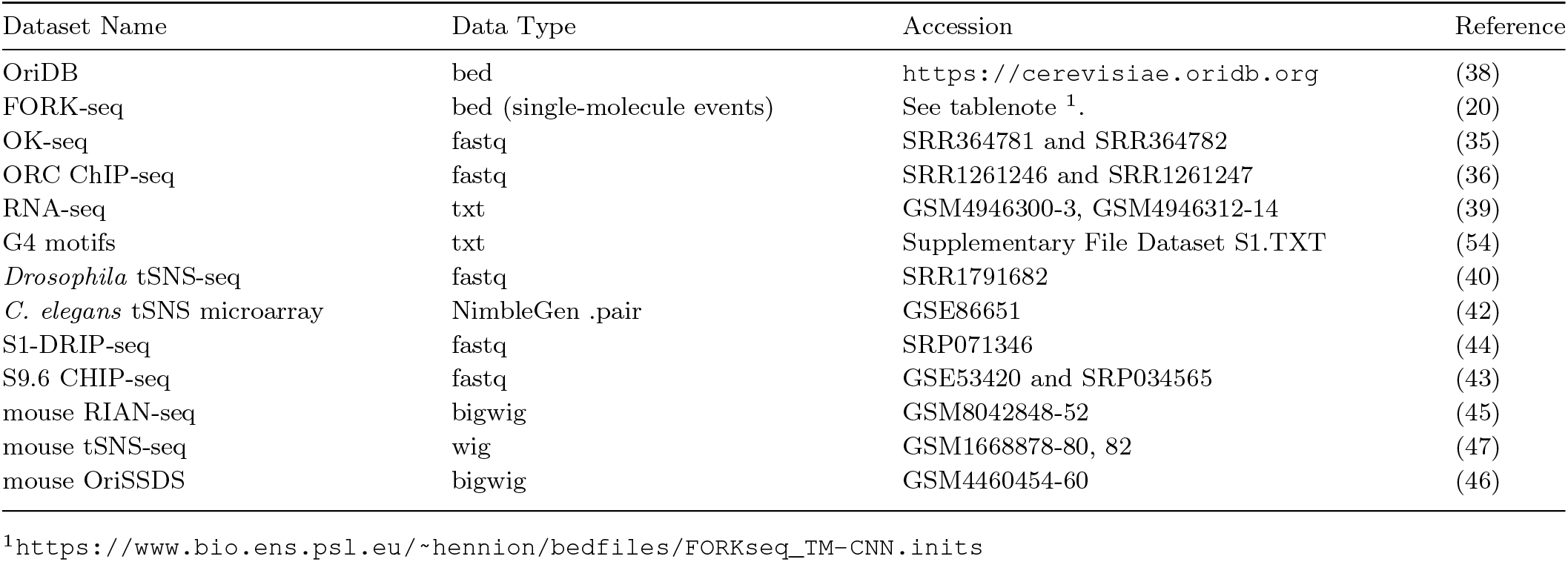
Datasets used in this study.

To define DNA replication origins in the OK-seq datasets (NCBI SRA Accession SRR364781, SRR364782) (35), we used Origin Efficiency Metric (OEM) peaks as described previously (35; 4) with the following modifications. Strand-specific paired-end reads were mapped with bowtie2 (–very-sensitive-local – soft-clipped-unmapped-tlen), and then sorted and filtered with SAMtools to retain only concordantly-mapped read pairs with MAPQ of at least 2 (view -q 2 -F 4 -f 3). BEDtools (37) was used to compute strand-specific concordantly-mapped fragment depth across the genome for both the positive (genomecov -bga -strand + -pc) and negative (genomecov -bga -strand - -pc) strands. The resulting variable-step bedGraphs were converted to single-bp fixed step bedGraphs with “varStepBdgToSingleStepBdg.py” (https://github.com/JohnUrban/sciaratools2), and piped into AWK to create simpler 1-based depth files that had 4 columns (chr, pos, strand, depth), which were then combined and sorted (by chr, pos, strand). The resulting file was the input to “strandSwitchMetrics.py” from the switchblade suite (https://github.com/JohnUrban/switchblade), which was used to compute Replication Fork Directionality (RFD) and origin efficiency metrics (OEM) profiles using 10 kb windows as adapted from McGuffee *et al*. (35). Replication origins were identified as local maxima in the OEM signal. Specifically, genomic regions closest to origins or initiation zones have positive OEM whereas genomic regions closer to termination zones have negative OEM. We first filtered out all genomic positions with OEM *<* 0, which left genomic intervals, or ‘positive zones’, that contain contiguously positive positions. Origin positions were identified inside each positive zone as the position with the maximum OEM value, or in other words, the OEM peak summit positions. The set of origin positions defined this way were then ranked by their summit-position OEM from highest to lowest OEM. Code to reproduce this can be found at https://github.com/JohnUrban/iSNS-seq. For FORK-seq, we used the file defining initiation events previously detected on single-molecule nanopore reads (20) (Table 2). To define origins in a comparable way in the FORK-seq dataset as in others, we started with 3-column BED files defining single bp mid-points of the single-molecule initiation events and aggregated them into initiation zones by clustering mid-points that were within 5 kb of each other (clusterBed -d 5000, which assigns a cluster ID in column 4), which was piped into BEDtools groupby (-grp 4 -opCols 1,2,3,4 -ops distinct,min,max,count) to learn the entire span of the initiation zone and number of initiation events associated with each. Then we defined the single bp preferred origin site within each initiation zone three different ways: the mid-point between the start and end of the zone with AWK, or by using BEDtools “groupby” as above but to find the mean (-grp 4 -opCols 1,2,3,4 -ops distinct,mean,mean,count) or the median (-grp 4 -opCols 1,2,3,4 -ops distinct,median,median,count) of the single-bp positions for initiation events within the initiation zone. We compared the performance of each of these three approaches by plotting sensitivity (SN) and positive predictive value (PPV) curves with respect to known origin locations (Supplementary Figure S3). The median origin site within each initiation zone optimized the SN and PPV, performing almost as well as using the entire length of the initiation zones, and was used as the single bp origin location. Larger origin regions were defined by adding flanks, such as 500 bp. Finally, FORK-seq origin sites were rank sorted from most to fewest number of single-molecule initiation sites within each zone. Code to reproduce this can be found at https://github.com/JohnUrban/iSNS-seq.

### Benchmarking against known replication origins

To benchmark tSNS-seq, iSNS-seq, and other origin mapping techniques (ORC, OK-seq, FORK-seq) on the yeast genome, we compared putative origin calls in each dataset to known replication origins from the OriDB database, which catalogs the origins in three classes: “confirmed”, “likely”, and “dubious” (38). In OriDB, origins were classified as “confirmed” if their replicator activity was confirmed by the autonomously replicating sequence (ARS) assay and/or with 2D gel analysis. Otherwise, they were classified as “likely” if they were identified by two or more genome-wide studies or as “dubious” if identified by only one genome-wide study. To determine the concordance between the origin-mapping methods and the set of known origin locations, we looked at (i) their sensitivity (SN; also known as *recall*), the percentage of known origin locations overlapped by peaks from a given method, and (ii) their positive predictive values (PPV; also known as *precision*), the percentage of putative origins (e.g. peaks) that overlap known confirmed origin locations. Putative origins in the datasets are henceforth referred to as “peaks”.

Factors such as different numbers of peaks or the lengths of peak intervals in different datasets can confound the interpretation of the SN and PPV comparisons between them. Therefore, to make better comparisons of the concordance between the various origin-mapping methods and the set of known origin locations, we controlled for peak interval lengths and peak confidence (e.g. origin efficiency) and plotted the SN or PPV as a function of peak confidence. Moreover, to estimate the expected SN and PPV that could be achieved at random, we generated a shuffled dataset in which peak intervals were randomly reassigned to new positions on the same chromosome. To control for peak interval lengths, we set all peak intervals to the same size by defining single bp origin locations and adding 500 bp to each side. For iSNS-seq, tSNS-seq, and ORC ChIP-seq, we used the single bp narrowPeak summit position defined by MACS peak calling. We used the single bp summit position defined by maximum OEM for OK-seq, the single bp median mid-point of initiation events inside initiation zones for FORK-seq, and the mid-point of the randomly-shuffled intervals. To control for peak confidence, each dataset was sorted by origin efficiency or origin strength such that the most efficient or strongest peaks (putative origins) were first and the weakest were last. In other words, we rank-sorted the peaks in each dataset from most to least confident, which was defined by -log_10_(q-value) for iSNS-seq, tSNS-seq, and ORC ChIP-seq, by Origin Efficiency Metric (OEM) for OK-seq, by the number of initiation events per initiation zone for FORK-seq, or a random order for randomly-shuffled intervals.

To summarize the performance of each origin-mapping dataset, we used the same number (390) of the best ranking peaks for each dataset controlling for peak interval length and peak confidence as for the above analysis (390 was chosen as it was the limiting number for OK-seq). For each comparison, all methods had the same number of their most confident peaks, all with the same peak widths. We used a simple peak overlap method to count all intervals in one set that overlap any interval in the other by at least 1 bp to get the observed overlap rate, then used the binomial approach (28) to compute the expected rate, the p-value of the observed rate, and the ratio of observed to expected. All statistics including SN, PPV, F-scores, and binomial analyses were automated by a custom python script (“originsense.py”) that can be found online (https://github.com/JohnUrban/iSNS-seq).

### Enriched feature analyses

The UMAP plots were created using python Umap-learn 0.5.7 package, using a feature matrix that was generated as follows: The GC contents were calculated for each peak region from the narrowPeak file using BEDtools nuc. The distances to the nearest feature were calculated from the peak summit using BEDtools closest with “-t all” parameter that reports all features in case there are ties. Genomic features were extracted from ensGene.v101.gtf file, which contains gene and transcript annotations based on Ensembl gene models and was downloaded from the UCSC (https://genome.ucsc.edu).

The data from the two replicates were combined, and the UMAP analysis was performed blind with regard to which replicate the data came from. The numerical data in the feature matrix was normalized with Z scoring and the features were one-hot encoded. The umap reducer parameters were: n neighbors=30, min dist=0.15, metric=‘manhattan’, random state=42.

### Transcriptome analysis

The poly(A)-derived transcriptome data (NCBI Gene Expression Omnibus GSE162197) was from (39) (Table 2). We converted the average gene-specific signal listed in the text file into bedgraph file and then averaged all six replicates and used that file in our analyses. Statistical analyses for transcription level correlation analysis were performed in R. Prior to analysis, transcription levels and enrichment signals of the tSNS-seq and iSNS-seq samples were log_2_-transformed and then normalized using z-scoring. Data points with a transcription level and tSNS-seq and iSNS-seq enrichment values of zero were removed. Spearman’s rank correlation was used to assess monotonic relationships between transcription level and enrichment of tSNS-seq and iSNS-seq. Statistical significance for Spearman correlation was assessed using a two-tailed test, with exact p-values disabled to account for tied ranks. Regression models and scatterplots were generated using ggplot2, with each dataset analyzed separately. Replicate pairs (iSNS-seq rep 1, 2 and tSNS-seq rep 1, 2) were treated as independent datasets for statistical comparisons.

### Analysis of *Drosophila melanogaster* and *Caenorhabditis elegans* genome-wide tSNS studies

To explore the association of traditional SNS-seq peaks with enriched features learned from yeast in other systems, *D. melanogaster* tSNS-seq data from S2 cells was downloaded (NCBI SRA Accession SRR1791682), mapped with Bowtie2 to the *D. melanogaster* genome-build (dm3) and parameters (-N 1 -L 22) specified in the original study (40), and sorted with SAMtools. Read depth was obtained using BEDtools (bedtools genomecov -bga -split) and converted to bigWig for viewing in IGV along with the SNS peak call intervals for S2 cells from the original study (GEO GSE65692 “S2 origins”). The latest gene annotation set from *D. melanogaster* genome build r6.64 was mapped to the older genome build (dm3) with Liftoff (41).

For *C. elegans* data, all tSNS NimbleGen Whole-Genome Tiling Array datasets from a previous study (42) were downloaded from NCBI (GEO Study GSE86651), which included three replicates each of pre-Gastrula embryos and mixed embryos (containing both pre- and post-gastrula stages), an arrested L1 larvae non-replicating control, and a mixed embryos RNase control. The average signal across replicates was used. The whole-genome tiling arrays (NCBI GEO Platform GPL16504) corresponded to *C. elegans* genome build WS180, which we validated by using BLAST to confirm probe coordinates. Therefore, to compare the SNS array data to tRNA genes, *C. elegans* genome build WS180 and its annotations were downloaded from WormBase. To plot heatmaps and profiles of SNS signal around tRNA genes, computeMatrix from DeepTools was used (–regionBodyLength 150 –beforeRegionStartLength 3000 –afterRegionStartLength 3000) for both *Drosophila* (–binSize 1) and *C. elegans* (–binSize 10 for pre-computed ratio profiles; –binSize 50 for raw log_2_ ratio heatmaps and profiles).

### Analysis of R-loop datasets in yeast

Sequence reads from two published R-loop mapping studies (43; 44) were downloaded from NCBI (see accessions in Table 2). All R-loop libraries were single-end, 50 bp reads. Reads were aligned to the sacCer3 reference genome using Bowtie 2.5.4 (with the following parameters: –local, –soft-clipped-unmapped-tlen, -D 500, -R 3, -N 0, -L 19, -i C,1,0, –score-min C,40,0).

Peaks were called from the resulting BAM files for each R-loop dataset using MACS 3.0.3 callpeak function with a q-value cutoff of 0.05 (-f BAM -g 1.2e7 –mfold 2 100 –fix-bimodal –extsize 200 –keep-dup all -q 0.05 –bdg), using the matched genomic or chromatin input sample as a control (-c) where available. Peaks for the input control datasets were called without a control.

Input-normalized R-loop coverage was computed from the BAM files as log_2_(IP/input) using deepTools 3.5.6 bamCompare (–binSize 25 –normalizeUsing CPM –scaleFactorsMethod None – extendReads 200 –pseudocount 1), and each resulting track was median-centered genome-wide (interval-length-weighted median subtracted to set the background to zero). Signal was summarized around gene bodies (scale-regions –regionBodyLength 150 -b 2000 -a 2000 –binSize 10) and around peak summits (reference-point –referencePoint center -b 2000 -a 2000 –binSize 20) using the deepTools computeMatrix, plotProfile, and plotHeatmap functions. Bins were averaged using the mean for heatmaps and the median for summary profile plots.

### Analysis of RIAN-seq and SNS-seq in mouse

Replicate-averaged signal tracks were generated for IGV visualization at near-native resolution (50 bp bins), using deepTools bigwigAverage. RIAN-seq (45) signal was averaged across the two wild-type replicates (GSM8042848, GSM8042849). OriSSDS (46) signal was averaged separately for the plus and minus strands across all five biological replicates (GSM4460454– GSM4460458); the minus-strand track was additionally multiplied by 1 during averaging (bigwigAverage –scaleFactors) so that it renders below the zero baseline in the genome browser. tSNS-seq signal (47) was averaged across all three biological replicates following liftover from mm9 to mm10.

For the correlation analysis, total RIAN-seq signal was defined as the mean of the two wild-type biological replicates. Eight individual OriSSDS and tSNS-seq (following liftover to mm10) replicates were compared against this signal. Genome-wide signal was quantified at four bin resolutions: 1 kb, 10 kb, 100 kb, and 1 Mb with deepTools (v3.5.4) multiBigwigSummary (bins mode). Wild-type RIAN-seq signal was quantified in the same bins at each resolution and matched to the SNS-seq replicate values by genomic coordinate. At each resolution, Spearman’s rank correlation was computed between each SNS-seq replicate and total wild-type RIAN-seq signal, restricted to bins with signal defined in both.

A permutation-based null distribution was generated for each replicate at each resolution by randomly reassigning that replicate’s bin values across its own set of covered bins (breaking any spatial correspondence with RIAN-seq signal while exactly preserving the replicate’s own marginal distribution of values) and recomputing the correlation. This was repeated for 100 independent permutations per replicate per resolution (1 Mb, approx. 2,700 bins). The null is reported as the mean *±* SD of Spearman’s *ρ* across the 100 permutations. Analyses were performed in Python (numpy, pandas, scipy) and results plotted with matplotlib.

Unless otherwise specified, all code used to perform the analyses described in this study is available at https://github.com/MiikoSokka/iSNS-seq.

## Results

### *λ*-exo slowly digests a 5’ RNA-DNA chimera, but is blocked by a G4 motif

Two assumptions have been made that are vital for a successful enrichment of short nascent strands (SNSs) using *λ*-exonuclease (*λ*-exo): (i) that *λ*-exo efficiently and uniformly depletes the background DNA, and (ii) that the 5’ RNA protects the SNS from digestion by *λ*-exo. The first assumption implies that no other factors protect DNA from digestion. Previously, we raised concerns that the first assumption is not fully supported by the data. Specifically, we found that *λ*-exo digestion is not uniform; it has a strong bias dependent on GC-content (28). Moreover, we found that G-quadruplex motifs (G4s) are enriched in *λ*-exo-digested preparations from non-replicating genomic DNA, suggesting that factors other than an RNA primer may protect downstream DNA. The second assumption that 5’ RNA protects the SNS DNA from *λ*-exo digestion is at the core of the SNS-seq approach, but remains to be rigorously tested. This presupposition is important because the typical SNS-seq experiment involves several overnight incubations with high enzyme-to-DNA ratios in order to remove and minimize the amount of background parental DNA. Not only must the 5’ RNA primers be stable to protect DNA from *λ*-exo digestion, but they must also offer protection that exceeds the spurious enriching effects from other factors.

We designed three oligomers to study the kinetics of *λ*-exo digestion on RNA-primed DNA and G4-containing single-stranded DNA (Figure 1A): (i) a chimeric oligomer with a 10 nt RNA at the 5’ end followed by 30 nt DNA (R_10_D_30_); (ii) an all-DNA oligo species with a well-studied human *Myc* locus G4 motif 10 nt downstream from the 5’ end (D_10_G4_27_D_23_), followed by 23 nt DNA; and (iii) a 60 nt single-stranded DNA oligonucleotide lacking any RNA or G4 sequence (D_60_) served as a control. Note that the 10 nt length for the 5’ RNA of R_10_D_30_ was chosen to simulate the average length of RNA primers synthesized by primase in yeast (48; 49).

We first performed a *λ*-exo digestion time series on each oligo individually to assess the kinetics of their degradation. The oligos were phosphorylated with T4 PNK and then 100 pmol of each oligo species was subjected to 10 U of *λ*-exo over the course of 30 min to 24 h (Figure 1B). The all-DNA control oligo was digested rapidly by *λ*-exo, with most of it gone within 30 min. The chimeric oligo (R_10_D_30_) was gradually digested by *λ*-exo and was mostly digested by 12 h, and almost completely digested by 24 h. Finally, while the first 10 nt of the 60 nt G4 oligo (D_10_G4_27_D_23_) was digested within 30 minutes by *λ*-exo, the resulting 50 nt oligo with a 5’ G4 motif (G4_27_D_23_) was very slowly digested with a significant amount remaining at 24 h. This is consistent with a G4 structure forming *in vitro* to block *λ*-exo and slowing further digestion. Compared with the control, the 5’ RNA offered some protection against *λ*-exo digestion as expected. Importantly, however, the digestion of the RNA-DNA chimeric oligo was faster than the digestion of the G4 oligo. This suggests that although 5’ RNA offers protection, it is not absolute and is less than the protection offered by the G-rich DNA that may form G4 secondary structures.

The *λ*-exo activity in digesting the chimeric RNA-DNA oligo was independent of the vendor, since the RNA-DNA chimera was depleted by *λ*-exo from all three vendors that were tested: Fermentas (which is now discontinued), ThermoFisher, and NEB (Supplementary Figure S4A). Digestion through the 5’ RNA by *λ*-exo is unlikely to result from hydrolysis, as hydrolytic cleavage would generate a 5’ hydroxyl group, which is a poor substrate for *λ*-exo (50). Moreover, no degradation of the RNA-DNA chimera was observed over the 16 h time course when using heat-inactivated *λ*-exo, indicating that the observed digestion was not due to RNase contamination (Supplementary Figure S4B).

To sum up, our results with short single-stranded oligomeric DNA show that while 5’ RNA gives some protection from *λ*-exo digestion, extended incubation will gradually digest the single-stranded RNA-DNA chimera. Importantly, the G4 motif appears to confer more protection from *λ*-exo digestion than RNA primers. This raises concerns that the second assumption of SNS-seq, described above, is also not supported. We conclude that proper controls, as well as improved conditions, are necessary to implement SNS-seq.

### Tris-HCl buffer increases *λ*-exo digestion through G4, while 5’-OH provides better protection than 5’ RNA

We investigated possibilities to improve the *λ*-exo step of SNS-seq. Our aim was to improve digestion through G4 DNA. The formation of G4 structures requires closely spaced guanine bases running on one DNA strand, which aligns in a tetrad lattice coordinated by a cation. The stability of the G4 structure is dependent on the type of cation around which it forms. For example, K^+^ cations help G4 DNA form one of the most stable structures (51). The glycine-KOH buffer that is traditionally used in *λ*-exo reactions contains a significant amount of K^+^ cations through the addition of KOH to adjust the pH to 9.5, which is the optimal pH for enzymatic activity of *λ*-exo (52). To eliminate the cations from the reaction buffer altogether to prevent G4 stabilization, we switched to using Tris-HCl at pH 9.0, which retains the buffering capacity of Tris-HCl (that becomes weaker at yet higher pH) and retains 90% of *λ*-exo activity (52).

We tested the Tris-HCl buffer in a *λ*-exo time course experiment, but this time added equal molar amounts of phosphorylated RNA-DNA chimera (R_10_D_30_), G4 oligo (D_10_G4_27_D_23_) and an all-DNA oligo (D_25_) in the same reaction. We also added a non-phosphorylated D_20_ oligo as a loading control since *λ*-exo has been reported to be inactive on 5’-OH DNA (50). Overall, this experimental design allowed us to quantify the degradation kinetics of different oligos, differentiated by their lengths, at the same time in the same reaction. As expected, *λ*-exo was again impeded in its digestion through the G4 motif of D_10_G4_27_D_23_ oligo in the traditional K^+^-rich glycine-KOH buffer (Figure 1C), with semi-quantification showing 28% of the G4 oligo remaining after 16 h, compared to 2% for the D_25_ all-DNA control (for semi-quantification, see Supplementary Figure S4C). However, the entire oligo (D_10_G4_27_D_23_) was digested in Tris-HCl buffer very efficiently, with barely visible bands after as early as 1 h of *λ*-exo digestion (1% remaining at 16 h, comparable to the D_25_ control). The RNA-DNA chimera (R_10_D_30_) was digested slowly in both buffers over the 16 h time course, though to a somewhat greater extent in Tris-HCl (35% remaining) than in Glycine-KOH (65% remaining). This shows that RNA-primers offer partial, but not full, protection from *λ*-exo digestion in either buffer, distinct from the buffer-dependent protection conferred by G-rich DNA in glycine-KOH buffer. Thus, conditions for better digestion through G4-containing DNA in Tris-HCl buffer may be more suitable for mapping origins with SNS-seq.

Although switching the *λ*-exo buffer from glycine-KOH to Tris-HCl improved digestion through G4 DNA, the issue of *λ*-exo digesting through the RNA primer remained unresolved, prompting us to explore an alternative strategy to enrich RNA-primed SNS.

The dependence of *λ*-exo on 5’ phosphorylation seemed to provide a solution, since only an oligo with a 5’-P can be digested (see Figure 1C, 5’-P D_25_ oligo) whereas an oligo with a 5’-OH will resist digestion by *λ*-exo (see Figure 1C, 5’-OH D_20_ oligo). In both glycine-KOH and Tris-HCl buffer the phosphorylated all-DNA oligo (5’-P D_20_) was gone similarly at 1 h. As a control for equal loading, the band intensity of the non-phosphorylated all-DNA oligo (5’-OH D_20_) remained the same throughout the 1–16 h time course.

When characterizing *λ*-exo properties *in vitro*, we found that the RNA-DNA chimera was not digested when 5’ phosphorylation by PNK was performed prior to RNA degradation, rather than after the removal of the RNA primer by NaOH (Supplementary Figure S4D); also confirming that a 5’ phosphate is required for *λ*-exo activity. This result prompted us to further assess how hydrolyzing RNA primers to obtain 5’-OH DNA protects short nascent strand (SNS) DNA from *λ*-exo digestion compared to 5’-phosphorylated DNA, and to devise an experiment that varied the order of RNA hydrolysis (NaOH) and phosphorylation (T4 polynucleotide kinase, PNK) (Figure 1D). In the enrichment (5’OH-ENR) sample, PNK treatment preceded RNA hydrolysis to leave SNS DNA with a 5’-OH and background DNA with a 5’-phosphate. Consequently, only background DNA is expected to be digested by *λ*-exo, leading to enrichment of SNS DNA. In contrast, in the no-protection control (CTR) sample, RNA hydrolysis was performed before phosphorylation to result in all DNA species with a 5’-phosphate equally susceptible to *λ*-exo digestion. In the latter case, both background DNA and SNS are expected to be digested.

We prepared CTR and 5’OH-ENR samples each with equal molar amounts of the RNA-DNA (R_10_D_30_), G4-DNA (D_10_G4_27_D_23_), and all-DNA (D_25_) oligo species. Following the PNK and NaOH steps, the samples were subjected to 16 h *λ*-exo digestion in the new Tris-HCl buffer conditions. As expected given our previous results, both the G4-DNA (D_10_G4_27_D_23_) and control (D_25_) oligonucleotides were completely degraded after 16 h of *λ*-exo digestion in both 5’OH-ENR and CTR samples, that is, they were degraded regardless of the order of PNK and NaOH treatment (Figure 1E). Since neither oligo contained an RNA primer, and the G4 structure was not stabilized in Tris-HCl, both were susceptible to digestion. In the 5’OH-ENR sample, only the 5’-OH D_30_ oligo (the resulting oligo species after removing the RNA primer from R_10_D_30_ with PNK →NaOH treatments in the 5’OH-ENR protocol) was protected against *λ*-exo digestion. In contrast, all oligos were completely digested in the CTR sample, as expected. We also ran the same experiment, but with *λ*-exo digestion in the traditional glycine-KOH buffer rather than Tris-HCl. The results were identical with respect to all oligos except the G4_27_D_23_ oligo, which remained undigested in both the 16 h CTR and 16 h 5’RNA-ENR samples (Supplementary Figure S6).

Overall, we conclude that (i) the 5’-OH left by the 5’OH-ENR protocol (PNK →NaOH) provides stronger protection of downstream DNA than the RNA primer, potentially enabling improved enrichment of short nascent strands in SNS-seq, and (ii) Tris-HCl buffer enhances *λ*-exo digestion through G4 motifs, thereby reducing non-origin enrichment.

### *λ*-exo enrichment of long SNS

The oligo experiments are useful for investigating how *λ*-exo operates on single-stranded DNA (and RNA) ends, but the ultimate goal is to enrich much longer, 0.5–2.0 kb, single-stranded RNA-primed short nascent strands (SNS) against a background of single-stranded parental DNA of equivalent length. These longer nucleic acids might behave in a different manner from short oligos, which prompted us to perform experiments with DNA of longer length. Using PCR amplification with primers containing 10 nt RNA at the 5’ end, we prepared a 1.2 kb chimeric RNA-DNA (SNS^mimic^) that mimics *in vivo* SNS to study how *λ*-exo digests longer DNA with a 5’ RNA primer. The 10 nt length for the RNA primer was again chosen based on reports that the RNA primer synthesized by yeast primase is, on average, 10 nt in length (48; 49). In addition, we prepared a 1.2 kb DNA (without an RNA primer) that contained several G4 motifs along its sequence that mimics G4-containing parental background DNA (BKG^mimic^) to investigate how well it is depleted by *λ*-exo. In all subsequent experiments, samples that test the selective enrichment (ENR) of SNS DNA using either the 5’-OH or 5’-RNA are referred to as 5’OH-ENR or 5’RNA-ENR, respectively, whereas the control sample that provides a baseline that reflects the total 5’-phosphorylated input DNA remaining after *λ*-exo digestion is referred to as CTR. As above, ENR and CTR refer to the order of addition of NaOH and PNK (Figure 1D).

We compared *λ*-exo digestion kinetics of mixtures of SNS^mimic^ and BKG^mimic^ using two SNS enrichment protocols: the 5’RNA-ENR protocol tested the protection by 5’-RNA primers in glycine–KOH buffer (PNK →*λ*-exo), and the 5’OH-ENR protocol tested protection by 5’-OH after RNA hydrolysis in Tris–HCl buffer (PNK →NaOH →*λ*-exo) (Figure 2A). For each, the DNA mixture was split in the beginning in equal aliquots, with one to be processed as the enrichment (ENR) sample and the other as a no-protection control (CTR) sample. The CTR samples were prepared by first hydrolyzing all RNA and then generating 5’-phosphorylated DNA (NaOH →PNK). However, the *λ*-exo buffer conditions differed based on whether the CTR sample was the counterpart to 5’RNA-ENR (in which case the *λ*-exo buffer was glycine–KOH) sample or 5’OH-ENR (in which case the buffer was Tris–HCl). This allowed us to monitor the level of each long DNA species (SNS^mimic^ and BKG^mimic^) using qPCR and compute the fold enrichment using the ENR/CTR ratio for each protocol.

**Fig. 2:**
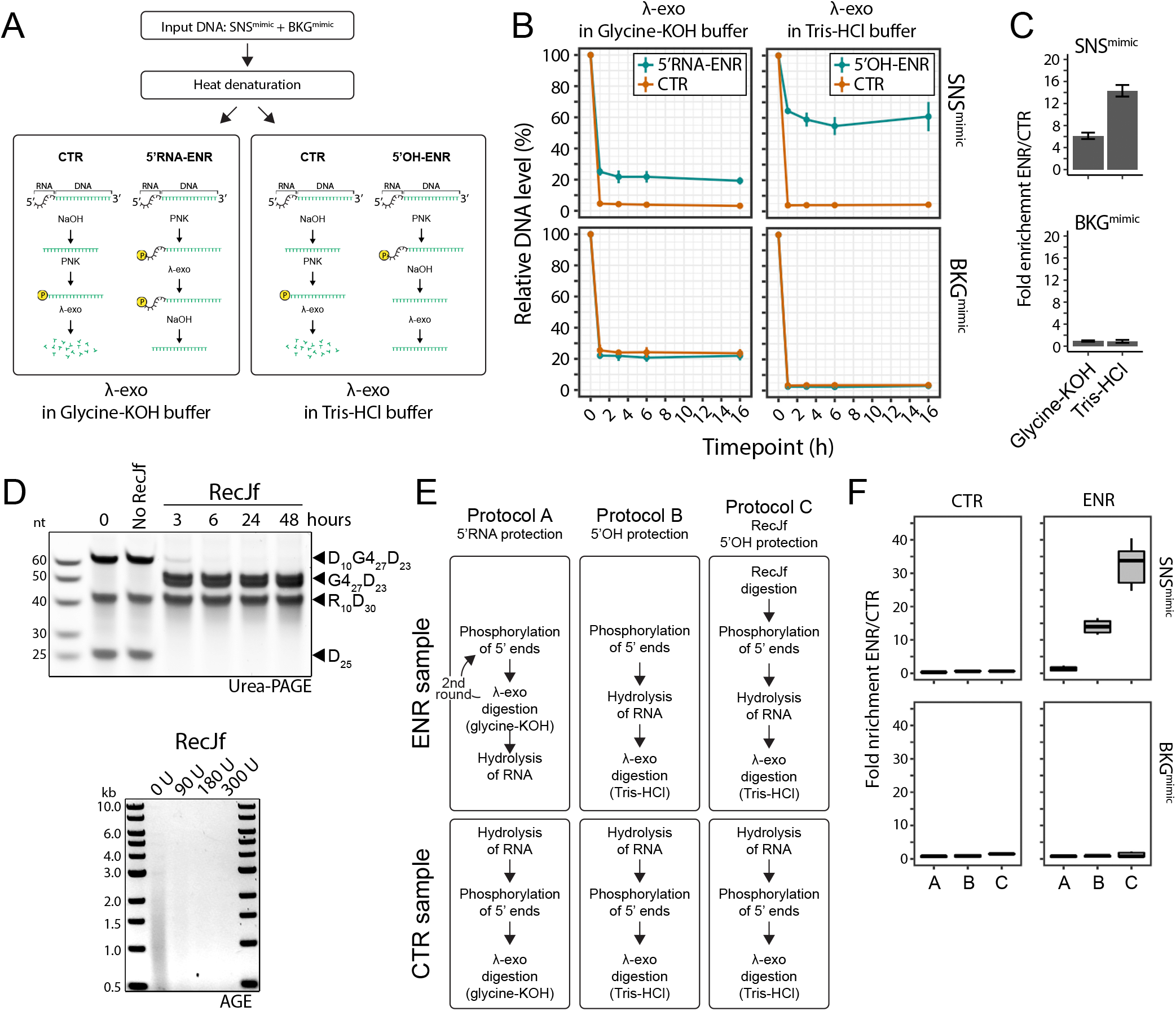
*λ*-exo digestion conditions using 5’ OH, Tris-HCl, and RecJf pre-treatment enhance SNS enrichment in long DNA assays. (A) Schematic of experimental steps used in panel B. 5’RNA-ENR employs *λ*-exo digestion in glycine-KOH buffer with a 5’ RNA-protected ENR sample (left). 5’OH-ENR employs Tris-HCl buffer with a 5’ OH-protected ENR sample (right). The CTR sample, treated with NaOH prior to *λ*-exo, serves as a control expected to degrade both SNS^mimic^ and BKG^mimic^. Note that the CTR sample procedure is the same in both protocols, only the buffer is different. (B) An SNS^mimic^ DNA and a background (BKG^mimic^) DNA were subjected to the conditions in panel A in the same time-course reaction. Each DNA species abundance was quantified by qPCR at indicated time points. Data represent mean *±* 1 standard deviation from three independent experiments. (C) Fold enrichment (ENR/CTR) values calculated from panel B. The location of each bar corresponds to the location of each plot in B, and is computed as the value of the cyan line (ENR) divided by the orange line (CTR) at the 16 h time point. (D) Analysis of RecJf digestion of oligomeric and genomic DNA. Top: urea-polyacrylamide gel electrophoresis (Urea-PAGE) of time course (hours indicated on the plot) of 60 U RecJf digestion of equal molarity of D_10_G4_27_D_23_ and R_10_D_30_ oligomers. Bottom: agarose gel electrophoresis (AGE) of RecJf titration (0–300 U) over 16 hours on sonicated yeast genomic DNA. (E) Schematic of experimental workflow used in panel F. (F) Boxplots showing fold enrichment (ENR/CTR) from experiment described in panel E. Fold enrichment is shown separately for SNS^mimic^ (top) and BKG^mimic^ (bottom). Boxplots represent median, interquartile range, and full data range.

In the 5’RNA-ENR sample, which used glycine–KOH buffer and a 5’ RNA primer to protect against *λ*-exo digestion, the SNS^mimic^ was rapidly depleted to 20–25% of the input within 1 h (Figure 2B, 5’RNA-ENR, top-left quadrant). This level persisted for the remainder of the time course, showing that the 5’ RNA primer conferred only limited protection against *λ*-exo digestion of the long (1.2 kb) SNS^mimic^. Most of the SNS^mimic^ was depleted to about 5% in the glycine-KOH conditions in the no-protection control (CTR) when the RNA primer was hydrolyzed off and the DNA end is phosphorylated, leaving no protection from 5’ RNA or 5’-OH (Figure 2B CTR in the top left quadrant). However, about 20% of BKG^mimic^ remained in the CTR sample, suggesting that *λ*-exo again had trouble digesting through the G4 DNA in the presence of K+ (Figure 2B, CTR and 5’RNA-ENR in the bottom left quadrant). In contrast, 5’-OH provided significantly better protection of SNS^mimic^: about 60% was preserved (Figure 2B, 5’OH-ENR in the top right quadrant). Moreover, in the CTR sample, both the SNS^mimic^ and BKG^mimic^ were reduced to 5% from the original (Figure 2B, CTR in the top and bottom right quadrant, respectively), showing again that *λ*-exo again had no trouble digesting through the G4 DNA in the absence of K+.

The observed fold enrichment of SNS^mimic^ as measured by ENR/CTR ratio at the 16 h time point was 14-fold in 5’OH-ENR Tris-HCl, but was only 6-fold for 5’RNA-ENR glycine-KOH (Figure 2C). As expected, the ENR/CTR ratio shows no enrichment of BKG^mimic^ in either of the procedures. This is because BKG^mimic^ has phosphorylated DNA at its 5’ end regardless of which protocol it is brought through (5’OH-ENR, 5’RNA-ENR, or CTR), and is consequently presented identically to *λ*-exo in the subsequent digestion step.

In summary, our longer DNA mimic experiments were consistent with our short oligomeric DNA experiments, and revealed two problems that the traditional glycine-KOH buffer has when combined with using 5’ RNA as protection from *λ*-exo digestion: (i) digestion through RNA-primed SNS^mimic^ by *λ*-exo leads to a very low fold enrichment, and (ii) the inability of *λ*-exo to digest through G4 structures can lead to erroneously identifying G4 motifs as enriched SNS. Conversely, the experiments show that the new approach of deliberately hydrolyzing the RNA primer to leave single-stranded DNA with a 5’-OH as protection from *λ*-exo digestion in Tris-HCl buffer significantly improved the SNS^mimic^ enrichment while allowing efficient depletion of BKG^mimic^.

### Increased SNS enrichment by using RecJf

The experiments performed above showed that although the *λ*-exo reaction kinetics were fast, the digestion reactions were inefficient in terms of achieving completion. We consistently found that approximately 5% of the DNA remained in the CTR samples after the 16 h *λ*-exo reaction, regardless of digestion buffer. In addition to the improvements offered by our new conditions above, we wanted to test if we can push the improvement further by testing an additional enzymatic step using RecJf as a pre-treatment before either of these steps. RecJf is an exonuclease that specifically digests single-stranded DNA from the 5’ end, but, unlike *λ*-exo, does not require the 5’ end to be phosphorylated.

We first tested whether RecJf digested the oligos R_10_D_30_, D_10_G4_27_D_23_ and D_25_ in a time course experiment similar to Figure 1B. Unlike the limited protection the 5’ RNA in R_10_D_30_ provided against *λ*-exo digestion, the 5’ RNA in R_10_D_30_ fully protected the oligo from digestion by RecJf: there was no reduction in the band intensity over the extended 48-h time course (Figure 2D top panel). However, similar to *λ*-exo, the RecJf was blocked by the G4 motif in the D_10_G4_27_D_23_ oligo, and unlike *λ*-exo, could not work its way through the block over time. In contrast, the all-DNA D_25_ oligo was efficiently digested by RecJf.

Moreover, RecJf significantly reduced the levels of yeast genomic DNA that had been sonicated to the average SNS size range (0.5–2.0 kb), RNase-treated, and denatured by heat (Figure 2D bottom panel). Note that in the iSNS-seq workflow, RecJf digestion serves as an additional bulk gDNA clean-up step upstream of *λ*-exo digestion. Any genomic DNA species that escape RecJf digestion, including those potentially protected by reannealed residual RNA, remain subject to the subsequent PNK, NaOH (that hydrolyzes residual RNA), and *λ*-exo steps that establish specificity for SNS molecules. These results demonstrate that RNA-primed DNA oligos are resistant to RecJf digestion and that RNA provided far more protection against RecJf digestion than against *λ*-exo digestion. Overall, these observations suggested that RecJf could increase the enrichment of SNS as part of an improved SNS-seq workflow.

In the next experiment, we compared three SNS enrichment protocols: (i) Protocol A with 5’RNA to protect the SNS, and two rounds of PNK and *λ*-exo in glycine-KOH, (ii) Protocol B with 5’OH and *λ*-exo reaction in Tris-HCl, and (iii) Protocol C with RecJf pre-treatment followed by Protocol B (Figure 2E). Note that in Protocol A, because traditional SNS-seq typically involves multiple rounds of PNK treatment and *λ*-exo digestion (24), we applied two consecutive PNK–*λ*-exo cycles in this experiment. The input DNA sample for these workflows was composed of a tripartite mixture of SNS^mimic^, BKG^mimic^, and yeast genomic DNA sonicated to an average of 1 kb (also hydrolyzed to remove all RNA). The rationale to include sonicated yeast genomic DNA was to make the sample more representative of SNS-seq experiments and to see if adding genomic DNA changes the reaction kinetics. As in the previous experiments, we prepared protection-based enrichment (ENR) and no-protection control (CTR) samples for each protocol in order to calculate the fold enrichment.

Protocol A with the two rounds of *λ*-exo digestion in glycine-KOH efficiently depleted background DNA but also led to nearly complete loss of all DNA, resulting in only a 2-fold enrichment of SNS^mimic^ (Figure 2F). In contrast, with Protocol B, SNS^mimic^ was enriched 14-fold, which is what we observed in the time course experiment without sonicated yeast DNA (see Figure 2C). Notably, Protocol C, with RecJf pre-treatment further increased the enrichment to 35-fold (Figure 2F). Overall, these results suggest that RecJf pre-treatment in combination with 5’-OH protecting the SNS in Tris-HCl buffer provides significantly better SNS enrichment than the traditional, two-round PNK and *λ*-exo treatment in glycine-KOH. From now on, in the context of genome-wide deep sequencing experiments that also involve an upstream size-selection step, we refer to Protocol C with RecJf pre-treatment as “improved SNS-seq” (iSNS-seq), and refer to Protocol A with the two rounds of *λ*-exo digestion as the “traditional SNS-seq” (tSNS-seq) method.

Benchmarking the performance of iSNS-seq, tSNS-seq and orthogonal origin mapping datasets in finding known origins

After optimizing the enzymatic reaction conditions to maximize the enrichment of SNS, we compared the improved SNS-seq (iSNS-seq) and traditional SNS-seq (tSNS-seq) protocols in budding yeast. The budding yeast replication origins have been well-defined by several other methods (listed in the OriDB database (38)), and thus serve as a good model for evaluating the origin mapping performance. We prepared two replicates of iSNS-seq and tSNS-seq samples using asynchronous logarithmic phase budding yeast cultures (Figure 3A). For each replicate, the genomic DNA was first heat-denatured to make it single-stranded (thereby releasing the nascent DNA from replication bubbles), then fractionated on a 5–30% sucrose gradient. Fractions containing 0.5–2.0 kb DNA were selected for further enrichment. This pre-enriched sample was split into two equal aliquots, one of which was processed using the tSNS-seq workflow and the other using the iSNS-seq workflow. Finally, the single-stranded DNA samples were made double-stranded through random priming and extension, fragmented to an average 300 bp, and prepared for paired-end sequencing by Illumina. By preparing both tSNS-seq and iSNS-seq samples from the same size-selected, pre-enriched material, we ensured that any differences between them are due to the enzymatic enrichment steps and not due to upstream steps (such as culture conditions or the efficiency of the size selection step). This sample pairing procedure was performed on two biological replicates such that there are two replicates for each tSNS-seq and iSNS-seq (four samples total). In order to monitor the success of each method, we added equal molar amounts of SNS^mimic^ and BKG^mimic^ from the experiments above as spike-ins before the size selection step for each replicate. The sequences of these added DNA were different from the yeast genome, allowing easy detection by sequencing.

**Fig. 3:**
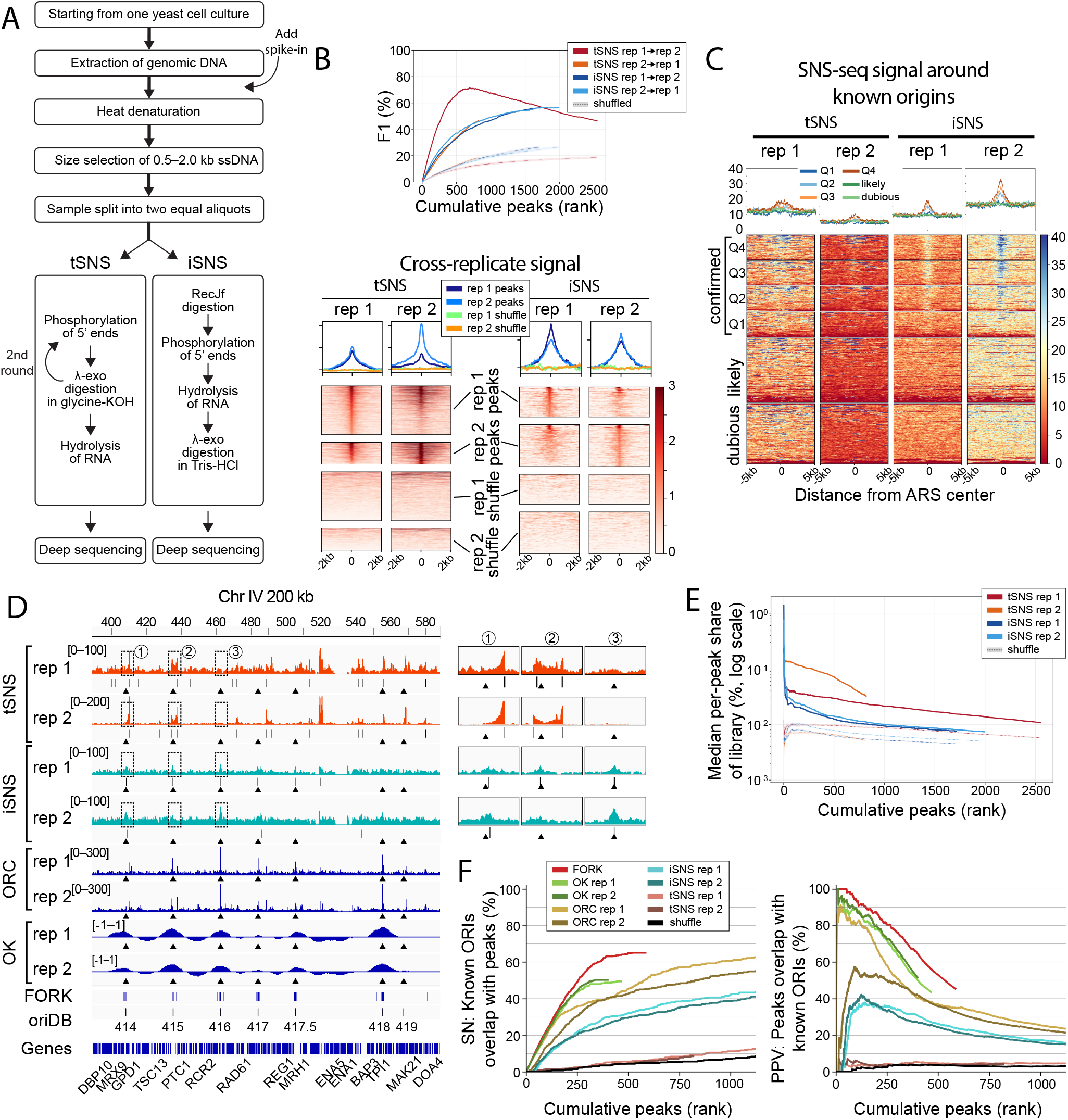
Characterization of improved and traditional SNS-seq datasets in *S. cerevisiae*. (A) Schematic overview of the experimental workflow. The size-selected, pre-enriched ssDNA sample was split into two equal aliquots immediately prior to enzymatic enrichment; one aliquot was processed by the tSNS-seq workflow and the other by the iSNS-seq workflow, such that tSNS-seq and iSNS-seq datasets derive from identical starting material and differ only in the enzymatic steps shown. (B) Cross-replicate F1 score as a function of peak rank. Peaks were ranked by q-value within each replicate, and F1 scores were calculated at each cumulative rank threshold N, using each replicate as the reference and the other as the query (both directions shown: rep1→rep2, rep2→rep1). Curves are shown for tSNS-seq and iSNS-seq separately, extending to each assay’s total called peak count. Dotted lines show the same comparison performed on randomly shuffled peak sets (matched peak widths and number, randomized genomic positions) as a null baseline. (C) Heatmaps of MACS3 pileup tracks generated when calling peaks of iSNS-seq and tSNS-seq within *±*5 kb of all known origins, stratified by classification (‘confirmed’, ‘likely’, and ‘dubious’; oriDB). The confirmed origins are further stratified by origin efficiency into quartiles (Q1=0.003–0.268; Q2=0.271– 0.488; Q3=0.489–0.672; Q4=0.673–0.936, where low and high origin efficiency is between 0 and 1, respectively) as determined in (53). Each row represents one origin. Median profiles per class are shown above the heatmaps. (D) Genomic browser view of a 200 kb region on *S. cerevisiae* chromosome IV (chrIV:388,444–588,444), highlighting seven confirmed origins. Pileup tracks (with range indicated) are shown for both replicates of iSNS-seq and tSNS. Summits of called peaks are indicated below each coverage track. FORK-seq initiation site midpoints, ORC ChIP-seq signal, and OK-seq origin efficiency metric (OEM) signal are shown below for comparison. The triangles below coverage/peak tracks show the location of the origins to help guide the eye. Three peak regions are highlighted and zoomed in on the right side to better show relative localization of signal over origins and peaks. (E) Median per-peak FRiP (Fraction of Reads in Peaks) curves as a function of cumulative peaks ranked by their q value. Curves are shown for both biological replicates of tSNS-seq (red and orange) and iSNS-seq (dark and light blue). Color-matched dashed lines show the same statistic computed on randomly shuffled peak sets (same peak widths and number, randomized genomic positions) as a null baseline for each replicate. The separation between each curve and its shuffled control reflects the degree to which sequencing signal is concentrated within called peaks relative to chance. (F) Sensitivity (SN, top panel) and positive predictive value (PPV, bottom panel) of each origin-mapping dataset with respect to confirmed origins (oriDB) as a function of cumulative peaks ranked by their q value. Length-normalized peaks were rank sorted from highest to lowest confidence (see Methods). Each rank value on the X-axis represents the x most confident peaks. Thus, for each x value, SN and PPV values on the Y-axis were calculated for the x most confident peaks. For example, when x=100, SN and PPV are calculated for the 100 most confident peaks in each dataset. The curve therefore represents the cumulative SN and PPV values across the rank sorted peaks (from best to worst) for each dataset. Moreover, the scores for different datasets are directly comparable at each x-value as the scores are controlled for the number of peaks and peak rank (both represented by the x-value) as well as length (all set to the same length; see Methods). This prevents erroneous conclusions from apparent differences in sensitivity (and PPV) that arise due to arbitrary peak calling decisions and technical factors, such as datasets having different numbers of peaks, different peak widths, or different stringency cutoffs.

Uniquely mapped reads originating from the spike-ins, SNS^mimic^ or BKG^mimic^, in each of the four samples were counted and normalized to the count of all uniquely mapped reads across these sequences and the yeast genome (counts per million, CPM) (Supplementary Figure S7). The BKG^mimic^ spike-in was depleted similarly in tSNS-seq and iSNS-seq samples, while the SNS^mimic^ spike-in was greatly depleted in the tSNS-seq samples compared to the iSNS-seq samples. We note that while iSNS-seq samples had much more SNS^mimic^ retained than tSNS-seq samples, the SNS^mimic^ spike-in DNA did not end up completely depleted in tSNS-seq samples, suggesting that when mixed with genomic background DNA, the RNA primer has a small protecting effect from *λ*-exo digestion as seen in our *in vitro* equivalent experiments (Figure 2B).

Overall, when using a q-value cutoff of 0.05, the tSNS-seq samples had 1,417 peaks and 635 peaks; the iSNS-seq samples had 258 and 362 peaks, in replicate 1 and 2 respectively. As an initial, threshold-dependent assessment of reproducibility, we compared the degree of intersection between called peak sets. Both iSNS-seq and tSNS-seq datasets showed moderately reproducible peak sets by this measure, with iSNS-seq replicates sharing 153 intersecting peaks (Jaccard = 0.33) and tSNS-seq replicates sharing 560 intersecting peaks (Jaccard = 0.38) (Supplementary Figure S8A and B). This apparent moderate reproducibility for tSNS-seq partly reflects the large difference in peak-set size between replicates: 88% of replicate 2 peaks were contained within the replicate 1 peak set, indicating substantial containment despite the modest Jaccard value. Notably, this within-method reproducibility does not imply concordance between the two methods themselves: only 3% (44 peaks) were shared across all four samples combined, indicating that tSNS-seq and iSNS-seq largely call peaks at different genomic locations despite each being internally reproducible (Supplementary Figure S8C).

To assess the reproducibility of iSNS-seq and tSNS-seq more rigorously than a fixed-threshold peak-overlap comparison allows, we computed the F1 score between biological replicates across the full range of peak ranks, using q-value to order peaks within each replicate (Figure 3B, top). Because this approach evaluates concordance at every possible rank cutoff rather than a single arbitrary threshold, it is not subject to the same sensitivity to near-threshold peaks that limits discrete overlap statistics such as Euler diagrams (Supplementary Figure S9). Both assays showed F1 scores well above the corresponding shuffled-peak baseline across the entire ranked peak set, indicating that replicate concordance for both tSNS-seq and iSNS-seq substantially exceeds what would be expected by chance. For tSNS-seq, F1 was higher at low ranks when replicate 1 was treated as the reference than when replicate 2 was, reflecting the disproportionate contribution of a small number of very strong peaks in replicate 2, rather than a general asymmetry in the comparison method. As we show below, these dominant tSNS-seq peaks correspond largely to non-origin sites rather than bona fide replication origins. For iSNS-seq, the F1 scores were similar regardless of reference direction, increasing more gradually across the full ranked peak set and converging with the declining tSNS-seq curve at approximately rank 2000.

As an independent measure of reproducibility that does not rely on peak calling, we assessed whether read density in one replicate was enriched at peaks identified in the other replicate (Figure 3B, bottom). For both tSNS-seq and iSNS-seq, signal from each replicate was clearly enriched at peaks called in the other replicate, well above the corresponding shuffled-region controls, confirming that replicate concordance is present at the level of underlying signal and not solely a property of the peak-calling procedure. Signal shape was broadly similar between the two assays, with tSNS-seq and particularly rep 2, showing a somewhat narrower, more sharply peaked enrichment profile at called peaks. Together, these results confirm that both tSNS-seq and iSNS-seq datasets are reproducible between biological replicates.

The MACS3-derived pileup tracks were visualized as heatmaps and summarized as profile plots depicting the median read-count signal across each origin class (confirmed, likely and dubious; Figure 3C). To test whether signal enrichment reflects true origin activity rather than a fixed set of confirmed loci, the confirmed origin class was further stratified into quartiles of equal size by origin firing efficiency, as determined by (53). Both iSNS-seq replicates showed the strongest signal enrichment around the highest origin firing efficiency quartile (Q4), with median signal decreasing progressively toward lower firing efficiency quartiles. This is consistent with the expected behavior of SNS-seq, which detects signal at each origin in proportion to its firing efficiency across an asynchronous population of cells. The tSNS-seq samples showed a similar but much less pronounced trend across firing efficiency quartiles, with substantially lower median enrichment overall. The likely and dubious origin classes showed no appreciable signal increase in either assay, consistent with these classes representing predominantly low-efficiency or unconfirmed origins.

Visualizing the signal and peak tracks in the genomic viewer showed that tSNS-seq samples had more prominent peaks compared to iSNS-seq samples (Figure 3D). However, those prominent peaks in tSNS-seq did not appear to be over sites of known origins, whereas the peaks in iSNS-seq, albeit less prominent, did appear to be over known origins (see the three zoomed in peak regions in Figure 3D). In fact, for tSNS-seq even short peaks were generally absent above known origins. This suggests (i) that tSNS-seq is enriching sites other than origins, and (ii) that the enrichment of short nascent strands from replication origins was more prominent in iSNS-seq in agreement with the spike-in results and *in vitro* experiments.

The 200 kb region of chromosome IV shown in Figure 3D contains seven confirmed replication origins annotated in OriDB (38), including ARS416 (also known as ARS1), a well-characterized highly efficient origin. Both iSNS-seq replicates showed clear peaks at ARS416, and six of the seven confirmed origins within this region had corresponding iSNS-seq peaks in at least one replicate. In contrast, neither of the tSNS-seq replicates exhibited a peak at ARS416, and the majority of tSNS-seq peaks in this region were located outside annotated origins, a pattern that is even more apparent in the close-up views of the seven origins shown in Supplementary Figure S10.

The apparent difference in peak sizes between tSNS-seq and iSNS-seq prompted us to quantify signal-to-noise, we calculated the median per-peak share of sequencing reads as a function of peak rank for each replicate (Figure 3E). Both iSNS-seq replicates showed closely overlapping curves, with per-peak read share consistently above their respective shuffled baselines across the full ranked peak set. tSNS-seq replicates showed substantially higher per-peak read share than either iSNS-seq replicate or the shuffled baselines, consistent with the higher raw signal-to-noise. This effect was considerably more pronounced in tSNS-seq replicate 2, whose peaks captured a markedly higher share of reads than replicate 1 across the shared rank range (Figure 3E). This is consistent with our F1 analysis above (Figure 3B), where the disproportionate contribution of a small number of very strong replicate 2 peaks elevated early-rank F1 when replicate 2 was used as the reference. However, as shown below, these dominant high-share tSNS-seq peaks correspond to non-origin sites. Thus, the higher apparent signal-to-noise ratio of tSNS-seq, particularly in replicate 2, reflects strong enrichment at non-origin loci rather than more efficient recovery of genuine replication origins.

We next evaluated how well the datasets captured the known origins. About 19% of the iSNS-seq peaks, and only about 5% of the tSNS-seq peaks overlapped with confirmed origins from the OriDB database (38) (Figure S9). However, this uncontrolled comparison does not account for differences in peak number, size, or confidence between methods, all of which can bias the comparison. We therefore determined the sensitivity (SN) and positive predictive value (PPV) of iSNS-seq and tSNS-seq peak sets with respect to known origins under conditions that control for these factors, and compared the results to the SN and PPV scores of other genome-wide origin mapping techniques performed in *S. cerevisiae*: OK-seq (35), ORC ChIP-seq (36), and FORK-seq (20). The putative origins in the tSNS-seq, iSNS-seq, ORC ChIP-seq, OK-seq, and FORK-seq datasets are called “peaks” for simplicity henceforth. For known origins, we used the 410 confirmed origins from OriDB as a gold-standard set. In this analysis, SN is the percentage of known origin locations overlapped by peaks from a given method, and PPV is the percentage of peaks that overlap known confirmed origin locations. To make direct comparisons of SN and PPV between origin-mapping methods, we controlled for the number of peaks, peak interval length, and peak confidence (e.g., score rank or q-value) (see Methods).

With respect to their sensitivity (SN) of detecting known origins, the origin-mapping approaches separated into four categories very early on in the process of assessing the N most confident peaks in each dataset, starting with the single most confident peak and iteratively adding the next most confident peak to the sets (Figure 3F, left panel). The FORK-seq and OK-seq datasets performed best, plateauing after about 350 and 450 peaks for OK-seq and FORK-seq, respectively, after which no new peaks overlapped with known origins. The ORC ChIP-seq and iSNS-seq datasets performed second and third best, respectively. Finally, tSNS-seq had the lowest SN for known origins. Notably, the SN of the two tSNS-seq replicates was only marginally higher than the SN of randomly shuffled peaks. The results were reproducible across replicates for each origin-mapping method. With respect to PPV, the origin-mapping approaches had the same trends, with FORK-seq and OK-seq having the highest PPV, followed by ORC ChIP-seq, then iSNS-seq, and with tSNS-seq having the lowest PPV that was again comparable to randomly shuffled peaks (Figure 3F, right panel). Interestingly, there is a delayed increase in the iSNS-seq, tSNS-seq, and ORC ChIP-seq replicate 2 PPV curves indicating that some of the most confident peaks in these datasets did not overlap with known origins. This suggests that some of the highly ranked peaks may be either origins that are not included in the known origin list, or peaks that are enriched by other unknown phenomena.

To summarize the SN and PPV curves in Figure 3F, we used the SN and PPV statistics from the 390 most confident peaks in each dataset (390 limit set by the number of peaks in OK-seq replicate 2). Compared to the gold standard set of confirmed origins, iSNS-seq replicates 1 and 2 exhibited stronger performance than tSNS-seq, with sensitivity (SN) ranging from 29% to 33%, positive predictive values (PPV) of 30% to 34%, and F-scores of 29% to 34%, all significantly exceeding random expectations (Supplementary Table S2). In contrast, both tSNS-seq replicates 1 and 2 exhibited markedly lower performance, with SN, PPV, and F-score all ranging between 6–8%, approaching values expected by random chance (5%). The SN and PPV of tSNS-seq replicate 1 reached statistical significance, but their values remained substantially lower than those of iSNS-seq replicate 1. The FORK-seq peaks performed overall best with the highest sensitivity (66%), highest positive predictive value (68%), and highest F-score (67%), all statistically significant. The OK-seq SN ranged from 55% to 56%, PPV from 57% to 59%, and F-scores from 56% and 57%, all statistically significant. The ORC ChIP-seq SN ranged from 40% to 41%, PPV ranged from 43% to 54%, and F-score values from 42% to 46%, all statistically significant values. We conclude that iSNS-seq significantly outperforms tSNS-seq for mapping replication origins, achieving sensitivity and precision approaching that of other methods such as ORC ChIP-seq, with FORK-seq and OK-seq the most sensitive and precise techniques overall.

### Traditional SNS-seq signal is associated with highly expressed genes

The tSNS-seq datasets had the lowest concordance with known origins, prompting us to investigate which genomic regions tSNS-seq is actually enriching. Specifically, we tried to understand if the non-origin peaks in tSNS-seq can be explained by some previously undetected phenomenon related to technical aspects of the tSNS-seq procedure.

To explore what biological features might be associated with peaks identified in the tSNS-seq and iSNS-seq datasets (peaks defined at q ≤ 0.05), we performed a Uniform Manifold Approximation and Projection (UMAP) dimensionality reduction on a curated feature matrix. Each peak was characterized by its GC content, q value, distance to the center point of the nearest genomic feature, and the identity of that feature. The list of genomic features and annotations included mRNA transcription start and end sites (TSS and TES, respectively) (n = 6,600), tRNAs (n = 299), small nucleolar RNAs (snoRNAs) (n = 77), non-coding RNAs (ncRNAs) (n = 18), known confirmed origins (n = 410) (38) and G-quadruplex (G4) motifs (n = 668) (54).

Consistent with our analyses above, UMAP projection of the iSNS-seq data revealed a prominent cluster corresponding to known origins, whereas only a very small cluster in the tSNS-seq projection corresponded to known origins (Figure 4). In contrast, the most notable feature of the tSNS-seq data was a distinct tRNA cluster, which was absent in iSNS-seq data. This tRNA cluster was driven by both tSNS-seq replicates, but they were organized by higher q values in the replicate 2 and lower values in replicate 1. This cluster had relatively short distances between the peaks and the tRNAs. There was also a snoRNA-enriched cluster in close proximity to the tRNA cluster, indicating a related feature space.

**Fig. 4:**
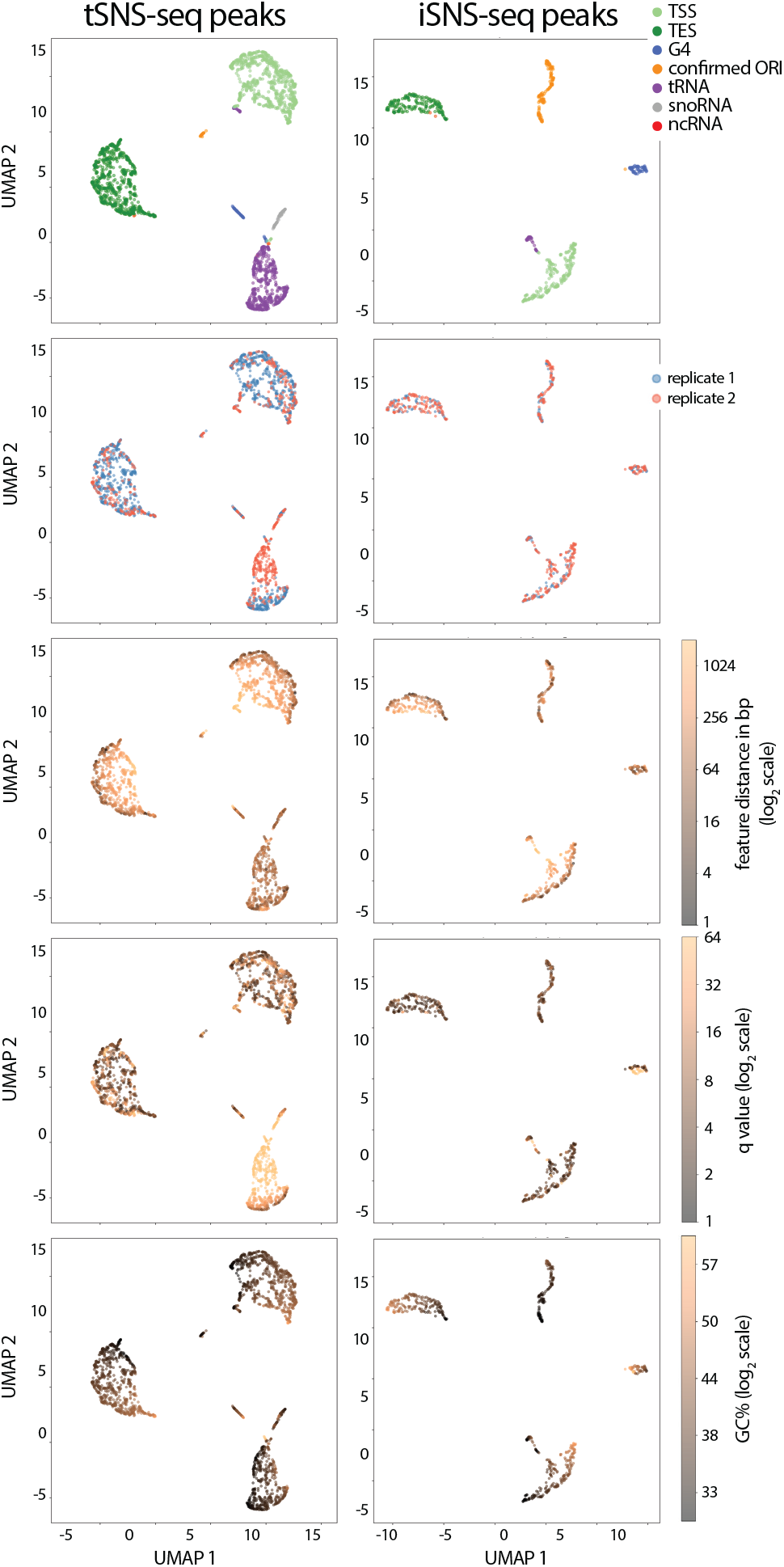
UMAP projection of tSNS-seq and iSNS-seq peaks reveals distinct feature associations. UMAP dimensionality reduction was performed on a feature matrix describing each peak by its GC content, q value, distance to the nearest genomic feature, and identity of that feature. Genomic annotations included mRNA transcription start sites (TSS) and transcription end sites (TES), tRNA genes, snoRNAs, ncRNAs, confirmed origins (ORIs), and G-quadruplex (G4) motifs. Panels show UMAP embeddings from top to down colored by genomic feature identity, dataset replicate, log_2_-transformed distance to nearest feature, log_2_(q value), and log_2_(GC%).

The mRNA TSS and TES clusters were among the largest in both the tSNS-seq and iSNS-seq datasets and these clusters exhibited mostly longer distances to their associated features. Their extended distance distributions combined with being the most numerous features in this analysis suggest that their proximity to peaks may reflect mostly a random association rather than a biologically meaningful enrichment.

GC content did not show a strong global trend across either dataset, reflecting the relatively uniform budding yeast GC distribution. G4 motifs formed small but distinct clusters in both datasets, but only 4–5% of peaks in either tSNS or iSNS datasets overlapped with G4 motifs (Supplementary Figure S11A). Furthermore, we did not observe a marked increase in the tSNS-seq or iSNS-seq signal over G4 motifs for either tSNS-seq or iSNS-seq (Supplementary Figure S11B).

Consistently with tRNAs and snoRNAs driving the separation of the tSNS-seq clusters, we observed that the strongest tSNS-seq peaks were associated with tRNAs and snoRNAs, rather than known origins (Figure 5A). We investigated this relationship further by plotting the tSNS-seq and iSNS-seq signals over non-coding genes tRNAs, snoRNAs, snRNAs and ncRNAs (Figure 5B). Both tSNS-seq replicates showed high signal over all gene classes, except ncRNAs. Interestingly, the signal was consistently skewed upstream from the gene body and abruptly ending at TES. Neither iSNS-seq replicates showed enrichment of signal over these regions.

**Fig. 5:**
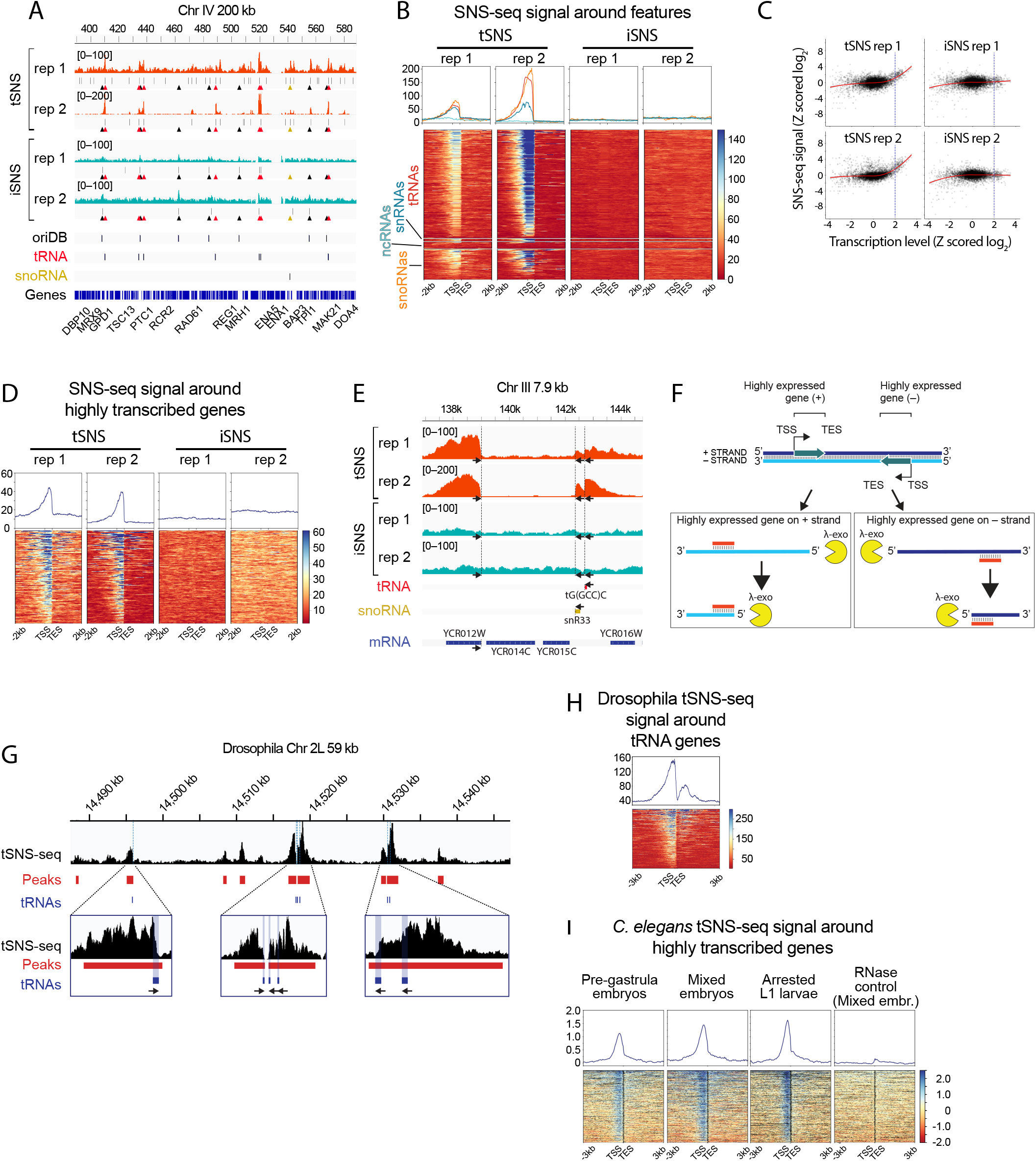
tSNS-seq peaks are enriched at highly transcribed genes and R-loops. (A) Genomic browser view of the same region as in Figure 3, with pileup track on top and peak summits below the tracks. Midpoints of confirmed origins (38), tRNAs and snoRNAs are shown both as black lines and color coded triangles (black, red and yellow, respectively) below pileup and peak tracks. (B) Heatmaps showing pileup of SNS-seq signal for iSNS-seq and tSNS-seq around tRNAs, snoRNAs, ncRNAs and snRNAs, spanning *±*2 kb from transcription start (TSS) and end sites (TES). Median signal profiles are shown above each heatmap. (C) Log_2_-transformed and Z-scored transcription signal (39) versus SNS-seq signal over mRNA gene regions. Loess regression curve is shown in red. The blue vertical dashed line is drawn at Z score = 2, separating the highly expressed genes. (D) Heatmaps showing pileup of SNS-seq signal over highly expressed genes (Z score *>* 2), spanning *±*2 kb from TSS and TES. Median signal profiles are shown above each heatmap. (E) A close-up IGV view of a 7.9 kb region on chromosome III, showing a highly expressed mRNA gene on the + strand on the left side (YCR012W; Phosphoglycerate kinase gene), as well as tRNA and snRNA genes on the – strand on the right side. The arrows point to the direction of transcription and the vertical dashed lines show the TES. (F) A schematic model illustrating how the skewed tSNS-seq signal arises from RNA:DNA hybrids formed between a gene’s RNA transcript and its complementary DNA strand. These hybrids block *λ*-exo digestion at the gene’s 3’ end (the transcription end site, TES) on the complementary DNA strand (or more generally, at the 3’ end of the RNA within the RNA:DNA hybrid). The schematic shows this on both the positive (+) and negative (–) strands with respect to the genome sequence. According to this model, in the genome tracks, a highly expressed gene on the + strand produces leftward-skewing signal, and a rightward-skewing signal when the gene is on the – strand. (G) tSNS-seq signal, called peaks, G4 motifs and tRNA genes in the *Drosophila* chromosome 2L, same region as Figure 3B from Comoglio *et al*. (40). Zoomed-in views highlight the previously unrecognized presence of tRNA genes at this locus, which coincide with the obstruction-shaped peaks. (H) Aggregate profile and heatmap of tSNS-seq signal (generated from Comoglio *et al*. (40)) around all annotated tRNA genes in *Drosophila*, spanning 3 kb from transcription start (TSS) and end sites (TES). (I) Median-normalized log_2_(Cy5/Cy3) *C. elegans* tSNS microarray signal around all tRNA genes, using tSNS samples from Rodríguez-Martínez *et al*. (42). Pre-gastrula embryos and mixed-stage embryos (both presumed actively replicating), L1 larvae (non-replicating control), and RNase-treated mixed embryos (RNase control) are shown.

Genome-wide, about 97% of the tRNA genes and 83% of the snoRNA genes overlapped with tSNS-seq peaks, compared to 6% and 0% overlapping with iSNS-seq peaks, respectively (Supplementary Figure S12A and B).

The surprising association between non-coding genes and tSNS-seq peaks prompted us to investigate whether coding genes exhibited a similar pattern. We plotted a poly(A)-derived transcriptome signal from Liu *et al*. (39) over tSNS-seq and iSNS-seq peak regions and observed that tSNS-seq, but not iSNS-seq, peaks were associated with high transcription levels (Supplementary Figure S13).

Next, we computed the average iSNS-seq and tSNS-seq signal over genes bodies (mRNAs) and then log_2_-transformed and Z-scored the SNS-seq and transcriptome signals in the 6,600 budding yeast genes. The scatterplot showed mostly no correlation between the SNS-seq and transcriptome signals. However, the most highly expressed genes had also the highest tSNS-seq signals, revealed by an upward progressing Loess regression curve (Figure 5C). Spearman’s rank correlation (*ρ*) indicated a weak but statistically significant positive association between transcription level and SNS-seq enrichment across all datasets, with tSNS-seq replicate 2 exhibiting the strongest correlation (*ρ* = 0.31, p *<* 0.001, N = 6472), followed by tSNS-seq replicate 1 (*ρ* = 0.22, p *<* 0.001, N = 6479). The iSNS-seq replicates 1 and 2 showed significantly weaker, albeit statistically significant associations (*ρ* = 0.08, p *<* 0.001, N = 6474 and *ρ* =0.05, p *<* 0.001, N = 6482, respectively).

When the tSNS-seq and iSNS-seq signals were plotted over only the most highly expressed genes (Z-scored log_2_ ≥ 2), the tSNS-seq signal showed similar skewing upstream of the gene bodies and abruptly ending at TES, than did the tSNS-seq signal over tRNAs (Figure 5D). The iSNS-seq samples did not show any enrichment over highly transcribed genes.

In order to get an idea of how much of the highly expressed genes explain the tSNS-seq peaks, we assessed the sensitivity (SN) and positive predictive value (PPV) of iSNS-seq and tSNS-seq in identifying the over two Z-scored mRNA genes, tRNAs and snoRNAs. In total, this set comprised of 565 highly expressed genes. The tSNS-seq replicates 1 and 2 reached very high sensitivity values of 76% and 81%, respectively (Supplementary Table S3). In other words, 76–80% of highly expressed genes were overlapped by tSNS-seq peaks. Similarly, the tSNS-seq replicates 1 and 2 achieved high PPV values, 29% and 67%, respectively. In other words, 29–67% of the tSNS-peaks overlapped highly expressing genes. These high SN and PPV values drastically shrink to about 15% when the peaks were randomly shuffled around the genome, demonstrating the effects are much higher than random. In contrast, the iSNS-seq replicates had sensitivity values around 10% and positive predictive values of about 20%. When randomly shuffling the tSNS-seq replicate 1 peaks (the highest number of peaks) across the genome, the SN and PPV values were 15% and 6%, respectively, showing this was close to what is expected at random. Collectively these results indicate that, likely due to the RNA hydrolysis step, our improved SNS-seq protocol drastically reduces this newly-discovered systematic RNA-linked bias of the tSNS-seq protocol.

The skewing and abrupt collapse of tSNS-seq signal at TES was observed in the vast majority of peaks spanning the entire budding yeast genome (Supplementary Figure S14, S15 and S16). The skewing is evident in the origin-centered close-ups in Supplementary Figure S10, corresponding to the chromosome IV region shown in Figures 3D and 5E. An illustrative close-up example of skewed tSNS-seq peaks is shown in Figure 5E) (see also close-ups in Supplementary Figure S17). A 7.9 kb region in chromosome III shows three highly expressing genes from different classes and on different strands, one on the plus strand (YCR012W mRNA) and two on the minus strand [tG(GCC)C tRNA and snR33 snoRNA]. Both tSNS-seq replicates show the skewing and abrupt ending of signal at the transcription termination site in all of these genes. The gene on the plus strand is transcribed from left to right, using the opposite strand as a template. The resulting transcript therefore has the same sequence as the strand to which the gene is assigned (in this case, the plus strand) and will hybridize with the opposite strand (see schematic in Figure 5F).

To investigate the universality of this bias mode in the traditional SNS-seq protocol, we used data previously generated for *Drosophila melanogaster* (40) and *Caenorhabditis elegans* (42), and focused on tRNA genes as a marker for the obstruction shape. Indeed, the obstruction-shaped peaks appeared to be associated with tRNA genes with the same exact placement and directionality learned from the yeast tSNS-seq experiments (Figure 5G, Supplementary Figure S18). The obstruction shape was even more apparent in the pileup of *Drosophila* tSNS-seq signal over tRNA genes when oriented by the strand (Figure 5H). Similarly, tSNS array samples from *C. elegans* stages including replicating pre-Gastrula embryos and mixed embryos, as well as non-replicating arrested L1 larvae, all had the obstruction shaped peaks associated with tRNA genes (Figure 5I, Supplementary Figure S19). However, it was completely removed in the RNase control sample, in which the extracted genomic DNA was treated with RNase A before *λ*-exo treatment. Although one interpretation is that the RNase destroyed the RNA primers of nascent strands, given our observations in yeast, it is more likely that RNase is destroying free tRNA that will otherwise be hybridized with denatured genomic DNA in some form of RNA:DNA hybrid that interferes with *λ*-exo digestion.

### Traditional SNS-seq signal correlates with R-loops in yeast and mouse

If RNA:DNA hybrids generate the tSNS-seq signal, tSNS-seq peaks might coincide with independently mapped R-loops. We therefore compared tSNS-seq and iSNS-seq profiles with published S1-DRIP-seq (44) and S9.6 ChIP-seq (43) datasets, including RNase H-deficient (RNase HΔ) samples from the same studies in which R-loops accumulate because of impaired R-loop resolution. tSNS-seq signal was clearly elevated at S1-DRIP-seq and S9.6 ChIP peak summits, and this enrichment was most pronounced in the RNase HΔ datasets (Figure 6A), consistent with tSNS-seq preferentially capturing sites of stabilized RNA:DNA hybrids. In contrast, iSNS-seq signal was flat across all R-loop datasets, showing no comparable enrichment. Both tSNS-seq and iSNS-seq signals were flat over the corresponding RNase-treated control peak sets and over shuffled intervals, indicating that the tSNS-seq enrichment depends specifically on the presence of RNA:DNA hybrids rather than some other feature of the underlying genomic loci.

**Fig. 6:**
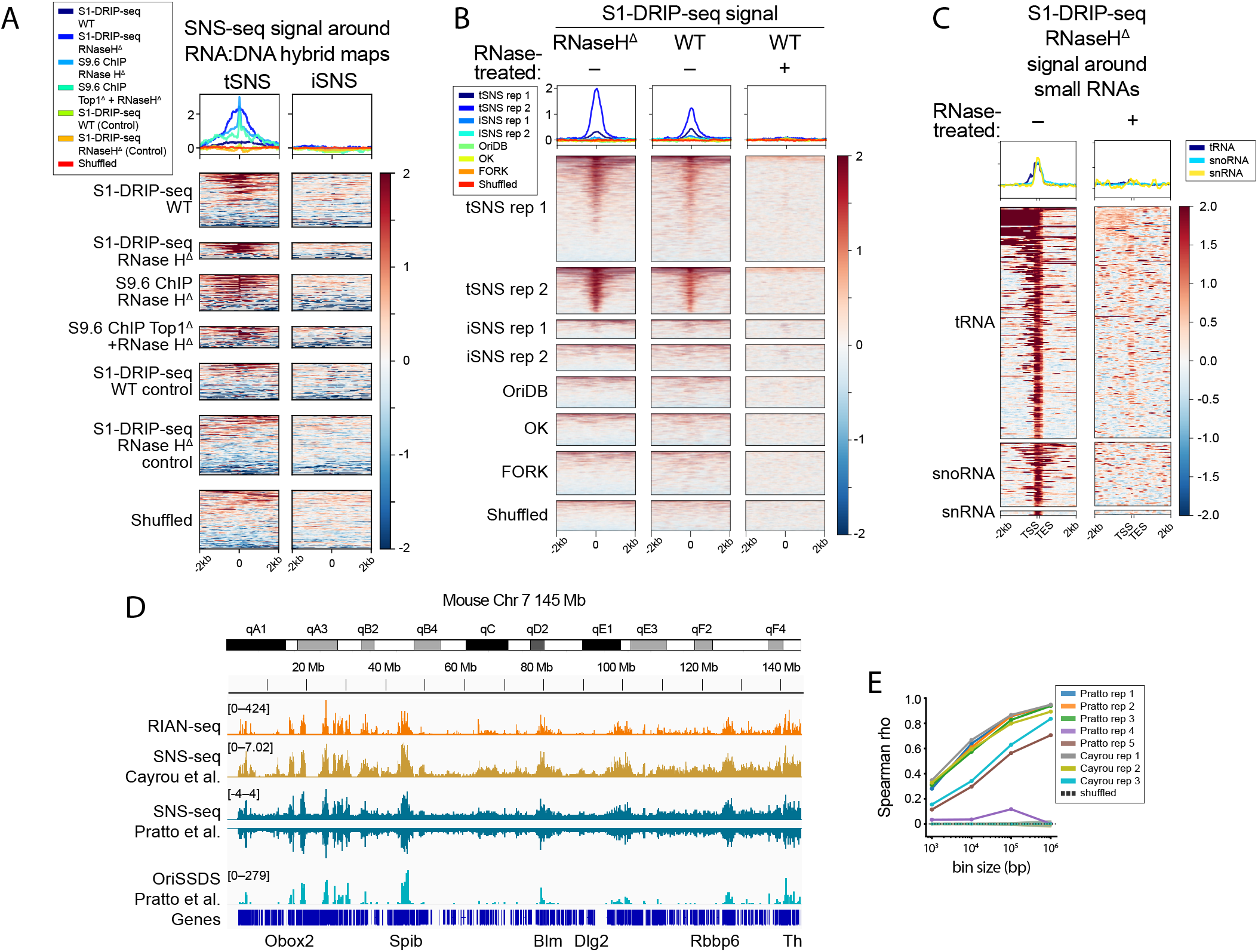
tSNS-seq signal correlates with R-loops in yeast and mouse. (A) Heatmaps showing log_2_ tSNS-seq and iSNS-seq signal centered on summits of peaks called from yeast RNA:DNA hybrid (R-loop) datasets S1-DRIP-seq (44) and S9.6 ChIP-seq (43), with corresponding median signal profile lines plotted above each heatmap. Each heatmap represents the log_2_ median-normalized average signal across replicates. (B) Heatmaps and aggregate profile of median-centered log_2_(IP/input) S1-DRIP signal in the RNase H^Δ^ and wild-type (WT) strain (with and without RNase H treatment, indicated by plus and minus sign) centered on peak summits called by tSNS-seq (rep 1, rep 2), iSNS-seq (rep 1, rep 2), confirmed origins (OriDB), OK-seq, and FORK-seq, as well as randomly shuffled control positions. (C) Heatmaps and aggregate profiles of log_2_ fold enrichment of S1-DRIP-seq signal in the RNaseH^Δ^ strain around tRNA, snoRNA, and snRNA genes (2 kb from TSS/TES), without (–) or with (+) in vitro RNase H treatment prior to DRIP. (D) An IGV view of mouse chromosome 7 (mm10) showing replicate-averaged signal tracks for RIAN-seq (45) (mean of two wild-type replicates), OriSSDS (46) strand-specific fragment coverage, shown separately for the plus and minus strands (mean of five replicates each), OriSSDS peak calls (mean of five replicates), and tSNS-seq (47) (mean of three replicates). (E) Spearman’s rank correlation (*ρ*) between total RIAN-seq signal (mean of the two wild-type replicates) and eight individual tSNS-seq/OriSSDS replicates in mouse, computed genome-wide at four bin sizes (1 kb, 10 kb, 100 kb, 1 Mb). For each replicate (colored line), solid lines with circular markers show the observed correlation; dashed lines with cross markers and shaded bands show the corresponding permutation-null mean *±* SD (100 independent permutations per replicate per bin size).

We next asked whether the converse relationship held: whether sites of genuine origin activity or orthogonal origin-mapping signal show any reciprocal enrichment of R-loop signal. Centering S1-DRIP-seq signal from wild-type cells (with and without RNase treatment) and the RNase H^Δ^ strain on peak summits from tSNS-seq, iSNS-seq, OK-seq, FORK-seq, and confirmed origins (OriDB) revealed robust enrichment exclusively at tSNS-seq peaks (Figure 6B and Supplementary Figure S20). This enrichment was strongest in the RNase H^Δ^ strain, intermediate in untreated wild-type cells, and was largely abolished by RNase treatment, consistent with an RNA:DNA hybrid-dependent signal. In contrast, iSNS-seq, OK-seq, FORK-seq, OriDB, and shuffled control peak sets showed no detectable enrichment in any condition. This shows that the DRIP-seq signal captured by S1-DRIP-seq is not associated with bona fide origins, but is specifically co-located with the sites preferentially enriched by tSNS-seq. Finally, we examined S1-DRIP-seq signal in the RNase H^Δ^ strain around tRNA, snoRNA, and snRNA genes, the same non-coding gene classes found above to drive tSNS-seq peak enrichment. Robust DRIP-seq signal was observed flanking the TSS and TES of these genes, and this signal was abolished when *in vitro* RNase treatment was applied prior to DRIP (Figure 6C and Supplementary Figure S21). This confirms that R-loops form at practically all small RNA genes in yeast and are readily detected regardless of assay, and it further supports our model that tSNS-seq peaks arise from *λ*-exo obstruction by RNA:DNA hybrids at these loci, rather than reflecting genuine replication initiation events.

Recently, RIAN-seq (R-loop identification assisted by nucleases and sequencing) was developed as an antibody-free, high-resolution approach for genome-wide R-loop mapping (45). Interestingly, RIAN-seq relies on a nuclease-based digestion strategy, where restriction-enzyme digested genomic DNA is treated with a cocktail of nuclease P1, T5 exonuclease, and *λ*-exo to selectively degrade single-stranded RNA, single-stranded DNA, and double-stranded DNA, while RNA:DNA hybrids are preserved and subsequently captured for sequencing. Importantly, RIAN-seq was developed entirely independently and for a different purpose than SNS-seq-based origin mapping, providing an orthogonal dataset against which to evaluate published SNS-seq datasets.

We compared mouse RIAN-seq signal (45) to two independently generated mouse SNS-seq datasets: strand-specific SNS-seq (Ori-SSDS) from Pratto *et al*. (46) and traditional SNS-seq from Cayrou *et al*. (47). Visual inspection of these datasets in a genome browser revealed a striking concordance between RIAN-seq signal and both mouse SNS-seq datasets (chr 7 shown in Figure 6D). This qualitative colocalization was supported by a substantial genome-wide correlation between RIAN-seq and SNS-seq signal across replicates and genomic scales (Spearman’s *ρ* up to 0.7–0.95 at 1 kb–1 Mb bin resolution (Figure 6E). One out of five Ori-SSDS replicates (rep4) showed negligible correlation with RIAN-seq signal at all bin sizes tested, indistinguishable from the shuffled-position null, and was therefore considered as a likely low-quality or low-complexity replicate. For the remaining replicates, correlation was consistently well above a randomized-position null generated by shuffling genomic bins (mean *ρ* = 0 across 100 permutations at each resolution), confirming that the observed concordance reflects genuine position-specific correspondence between SNS-seq and RIAN-seq signal rather than an artifact of genomic binning.

Taken together, mouse SNS-seq signal colocalizes substantially with independently mapped R-loops, consistent with RNA:DNA hybrid formation contributing to SNS-seq enrichment beyond the yeast system. This parallels the tSNS-seq bias mode we characterized in budding yeast, *Drosophila*, and *C. elegans*. Because this transcript-dependent bias generalizes across species and across studies from multiple independent laboratories, we conclude that the peaks generated by traditional SNS-seq represent, at best, a mixture of weakly enriched origins and strongly enriched artifacts, and at worst, are nearly all artifactual. This bias complicates the interpretation of RNase-treated and non-replicating controls, whose transcriptional landscapes differ from that of the experimental sample.

## Discussion

In this study, through *in vitro* experiments, we first investigated and improved the reaction conditions of *λ*-exo within the context of enriching short nascent strands (SNS) for SNS-seq. These experiments utilized both short oligomeric DNA and longer PCR-generated DNA that matched the size of short nascent strands. Both experimental designs included tests on RNA-DNA chimeras to simulate RNA-primed nascent strands as well as tests on G-quadruplex (G4) motifs, a known impediment to *λ*-exo digestion in traditional SNS-seq conditions. Our goal was to refine the SNS-seq protocol to minimize systematic false-positive enrichments and preferentially enrich true DNA replication origins. The *in vitro* experiments revealed that the optimized conditions enhanced the recovery of RNA-primed DNA and markedly diminished the enrichment of G4 DNA. To benchmark our new conditions, we compared our improved SNS-seq (iSNS-seq) protocol to the traditional SNS-seq (tSNS-seq) protocol using budding yeast cells where the genomic locations of DNA replication origins are known. To our surprise, our data revealed a novel major systematic bias of tSNS-seq at sites of highly expressed genes, coinciding with known R-loops. The iSNS-seq protocol increased the detection of known origins compared to tSNS-seq and, importantly, did not share the major bias mode of tSNS-seq.

The *λ*-exo digestion of short oligomeric DNA and longer PCR-generated DNA stops at G4 motifs in the buffer commonly used in traditional SNS-seq protocol. This is consistent with the previous findings that *λ*-exo is paused by a GC-rich sequence (25) and blocked by G4 motifs (28). It is likely that G4 structure formation occurs in the reaction solution during the *λ*-exo reaction, and may be dependent on conditions that allow G4 formation. To support this notion, we demonstrated that the G4 bias is largely buffer-dependent, consistent with G4 formation being less favorable in Tris-HCl buffer (51) and allowing *λ*-exo to digest through G4 DNA.

Given the clear G4-mediated *λ*-exo obstruction we observed in traditional conditions *in vitro*, as well as previously demonstrated genome-wide blockage at G4s in human cells and preference of *λ*-exo for AT-rich over GC-rich DNA (28), we were somewhat surprised not to observe a higher proportion of G4-overlapping fragments in tSNS-seq compared to iSNS-seq. However, this is not entirely unexpected for several reasons. First, the budding yeast genome is markedly more uniform than the human genome: local GC content (100 bp bins) spans a 10th–90th percentile range of 30–46% in yeast versus 26–55% in human, with roughly half the standard deviation and MAD around the mean (Supplementary Figure S22). Budding yeast also has substantially fewer G4 motifs than humans, both in absolute number and as a proportion of the genome (55), limiting the opportunity for a genome-wide G4 signal to emerge regardless of any per-motif blocking effect. Finally, we cannot rule out that budding yeast G4 motifs simply form less stable quadruplexes, as has been shown for yeast telomeric G4s specifically (56).

Importantly, our *in vitro* analyses revealed that although a 5’ RNA primer slows *λ*-exo digestion under traditional SNS-seq conditions, it does not completely protect single-stranded DNA from degradation, consistent with previous observations at high enzyme-to-DNA ratios (57). These findings suggest that many RNA-primed nascent strands would be degraded during the enrichment procedure. Moreover, G4-containing DNA proved more resistant to *λ*-exo digestion than RNA-primed DNA, implying that repeated or prolonged digestion preferentially enriches G4-containing contaminants while simultaneously destroying bona fide nascent strands. Thus, increasing *λ*-exo concentration or performing multiple rounds of digestion, a common strategy for reducing background DNA (24), is unlikely to improve the specificity of SNS enrichment.

These observations motivated the development of improved SNS-seq, in which RNA-primed termini are converted into *λ*-exo-resistant 5’-OH ends while contaminating DNA remains susceptible to digestion. An additional RecJf digestion step further enhances the selective removal of background DNA.

To directly assess the impact of these modifications, we benchmarked iSNS-seq against the traditional tSNS-seq protocol using the same size-selected DNA preparation from asynchronously growing budding yeast, split immediately before the enzymatic enrichment steps. This design minimized upstream technical variation, allowing differences between the protocols to be attributed primarily to the enrichment procedure itself, including the retention of RNA in tSNS-seq and its removal in iSNS-seq. Under these conditions, iSNS-seq substantially improved the enrichment of bona fide replication origins. Although it did not outperform orthogonal origin-mapping approaches, it represents, to our knowledge, the first SNS-based protocol to achieve genome-wide enrichment of independently validated replication origins, whereas the poor performance of tSNS-seq highlights the impact of systematic biases on traditional SNS-seq.

Agreement between results from different ORI mapping methods has been considered proof of their validation. For example, INI-seq, which does not use *λ*-exo, has been reported to overlap with tSNS-seq at a subset of efficient human origins (16), and this overlap has been invoked as cross-method validation. However, the interpretation of such overlap is complicated by several factors that have been noted in independent reviews. Different human replication origin maps show overlap that is in some cases not substantially greater than that obtained from random sets of loci (9), and quantitative analyses reveal striking asymmetries that inflate apparent concordance: 88% of INI-seq 2 origins (total 23,905) overlap with only 12% of tSNS-seq origins (total 175,536) in the same cells (16; 10), reflecting the high baseline probability of overlap when peak sets differ substantially in number. These considerations do not exclude the possibility that the most efficient, multiply-confirmed origin sites represent bona fide initiation events, but they argue against treating cross-method overlap as sufficient evidence that tSNS-seq peaks are free of the systematic biases described here.

One of the most striking and consequential findings of our study is that the traditional SNS-seq protocol preferentially enriches non-origin DNA. Among all methods evaluated, tSNS-seq showed the lowest concordance with established replication origins, performing only slightly better than random. Instead, tSNS-seq peaks were strongly associated with genes encoding highly abundant RNA species, including tRNAs and snoRNAs.

Because tSNS-seq and iSNS-seq were prepared from the same input material and differ in whether RNA is retained (tSNS-seq) or hydrolyzed (iSNS-seq) prior to *λ*-exo digestion, iSNS-seq provides an internal RNA-depleted control. The selective enrichment observed only in tSNS-seq therefore indicates that retention of RNA during the *λ*-exo step generates these non-origin signals.

A plausible explanation is that RNA molecules retained during tSNS-seq form RNA:DNA hybrids that impede *λ*-exo digestion. Consistent with this model, tSNS-seq peaks at highly expressed RNA genes display a characteristic asymmetric profile, with an abrupt loss of signal at the transcription end site and enrichment extending toward the transcription start site and upstream of the gene. This distinctive obstruction-shaped signature was also evident in previously published tSNS-seq datasets from *Drosophila* and *C. elegans*, suggesting that it is an inherent feature of the traditional protocol rather than a species-specific phenomenon. We propose that the transcribed RNA hybridizes to the complementary DNA strand, forming an RNA:DNA hybrid as illustrated in Figure 5F. During digestion, *λ*-exo progresses along single-stranded DNA in the 5’ to 3’ direction until it encounters this hybrid, where digestion is blocked. DNA molecules therefore accumulate with sharply defined 5’ ends at the position corresponding to the transcription end site, producing a strand-dependent asymmetric peak: left-skewed for genes on the plus strand and right-skewed for genes on the minus strand.

Further support for this model comes from the close correspondence between tSNS-seq artifacts and independently mapped R-loops. R-loops are RNA:DNA hybrid structures that preferentially form at highly transcribed loci, including tRNA and snoRNA genes (44). Consistent with this distribution, tSNS-seq, but not iSNS-seq, colocalized with R-loops mapped independently by both S1-DRIP-seq and S9.6 ChIP-seq. A recent study of *Trypanosoma brucei* replication origins found that 90% of strand-specific tSNS-seq peaks overlapped known R-loops (58), and we likewise observed a strong correspondence between two independent mouse tSNS-seq datasets (47; 46) and R-loops mapped by RIAN-seq (45). The specificity of this association is underscored by the absence of reciprocal R-loop enrichment at iSNS-seq, OK-seq, FORK-seq, or experimentally confirmed origin sites, as well as by the loss of R-loop signal at non-coding RNA genes following RNase treatment. Together, these observations from independent laboratories, multiple R-loop mapping technologies, and evolutionarily divergent species strongly support RNA:DNA hybrid-mediated obstruction of *λ*-exo as a general feature of traditional SNS-seq rather than a yeast-specific or laboratory-specific artifact.

Importantly, we do not propose that *in vivo* R-loops survive the denaturation steps of the iSNS-seq protocol. Rather, we propose that RNA:DNA hybrids reform *in vitro* after denaturation at genomic loci that are prone to R-loop formation. The likelihood of hybrid reformation would be expected to increase with local RNA abundance. These reformed hybrids would then selectively block *λ*-exo digestion, producing the characteristic asymmetric enrichments observed in tSNS-seq.

A significant contributor of controversy in the origin mapping field, particularly around tSNS-seq results, has arisen from tSNS-seq controls where the RNA is removed either with RNase, or by hydrolysis, leading to abolishment of the SNS-seq signal enrichment (24; 46). The common interpretation is that RNase destroyed the RNA-primer on nascent strands that normally protect them from digestion, thereby leading to their enrichment. However, in light of our observations, a new and more likely interpretation is that RNase or hydrolysis destroys the RNA before it can form RNA:DNA hybrid structures. Indeed, this is exactly what we see in the RNase control for tRNA genes in *C. elegans* (42) (Figure 5I and Supplementary Figure S19). Our own experimental design provides a direct parallel to these RNase controls: tSNS-seq and iSNS-seq were prepared from the same size-selected genomic DNA, differing only in that residual cellular RNA was hydrolyzed prior to enzymatic digestions in iSNS-seq but retained through this step in tSNS-seq. iSNS-seq can therefore be considered an internal, within-system RNase control, and the loss of the tRNA/snoRNA-associated peaks in iSNS-seq is consistent with the interpretation that RNA hydrolysis eliminates RNA:DNA hybrid-mediated obstruction rather than depleting a protective RNA primer.

Finally, the RNA:DNA hybrid formation could also explain why the traditional SNS-seq approach maps origins within active genes that are not mapped there by other techniques and why the read depth is sometimes correlated with exons (*Myc* gene) (28). It may also explain the discrepancy observed between tSNS-seq and OK-seq results in *Caenorhabditis elegans*. OK-seq identified initiation zones that remain largely stable across developmental stages and are located near early-activated genes (59). In contrast, tSNS-seq showed dynamic changes in enrichment patterns, that shifted across the genome in parallel with transcriptional activity during *C. elegans* development (42). Combined with the previously characterized G-quadruplex and GC-rich biases of *λ*-exo (25; 26; 27; 28), transcript-mediated obstruction represents an additional and potentially more pervasive bias mode that operates at any locus producing complementary RNA. Strong, clean peaks in SNS-seq datasets are not, in themselves, evidence of bona fide origin enrichment; they may instead reflect the selective retention of off-target species whose starting abundance exceeds that of genuine nascent strands.

Recent efforts have sought to reconcile the apparently contradictory pictures of metazoan replication initiation provided by tSNS-seq and zonal methods, proposing that discrete tSNS-seq signals correspond to efficient initiation sites clustered within broader zones detected by OK-seq and related techniques (10). Our findings complicate this hypothesis without invalidating it: if a substantial fraction of discrete tSNS-seq signals reflect RNA:DNA hybrid artifacts rather than bona fide initiation events, then attributing biological significance to discrete tSNS-seq peaks within initiation zones requires first distinguishing artifactual peaks from authentic intermediates. The improved protocol described here provides one route toward this distinction by eliminating the RNA:DNA hybrid contribution before *λ*-exo digestion.

Overall, we conclude that the interference of RNAs complementary to genomic sites with *λ*-exo digestion represents a plausible mechanism for an additional violation of the assumption of uniform depletion of parental DNA, and may represent a pervasive systematic factor in traditional SNS-seq datasets. The significance of such an artifact is underscored by a rarely discussed constraint: the intrinsic signal-to-noise ratio of the starting material. Cadoret *et al*. estimated that around 10 ng of 1.5–2 kb short nascent strands could theoretically be recovered from 10^8^ human cells, assuming 100% origin efficiency (11). Under these idealized conditions, this corresponds to 8 *×* 10^9^ SNS molecules, whereas the same cell population contains 3.7 *×* 10^14^ genomic DNA fragments of comparable size if parental DNA is randomly broken during extraction. Thus, even under assumptions deliberately favorable to SNS-seq, bona fide SNS molecules would represent only 1 in 46,000 fragments in the input material, and in asynchronous populations with stochastic origin firing, the true fraction is likely to be much lower. A parallel calculation in budding yeast yields 1 SNS per 380 fragments under maximal origin activity, indicating that yeast should be a substantially easier substrate for SNS-seq than the human genome.

These ratios refer to the unfractionated starting material, and the size-selection step that precedes *λ*-exo digestion is intended to improve them by depleting background DNA outside the 0.5– 2 kb single-stranded range. However, the magnitude of this pre-enrichment has, to our knowledge, never been rigorously quantified, and it is widely acknowledged that the size-selected fraction still contains a substantial excess of parental DNA over nascent strands. Indeed, the very rationale for the *λ*-exo step is the persistence of contaminating background after size selection. The size-selection step therefore improves the starting ratio by an unknown factor, leaving *λ*-exo to perform the bulk of the remaining enrichment against a background that is still in considerable excess.

This is the regime in which off-target biases of *λ*-exo become consequential. The modest enrichments we observe with iSNS-seq in budding yeast are consistent with the intrinsic constraints on what iSNS-seq can achieve when the enrichment step itself is unbiased. By contrast, the several-fold enrichments reported in some human tSNS-seq studies are difficult to reconcile with a starting ratio two orders of magnitude less favorable than the yeast case, even allowing for substantial pre-enrichment by size selection. Such enrichments are more readily explained by contributions from non-SNS sources whose abundance is not bounded by the theoretical SNS yield — including the RNA:DNA hybrid-mediated enrichment described here, which has no inherent abundance ceiling tied to replication initiation and can plausibly produce strong, reproducible, and locus-specific but erroneous signal at highly transcribed regions.

A corollary of this analysis is that the signal-to-noise ratio achievable by iSNS-seq in its current form is unlikely to be sufficient for de novo origin mapping in organisms where the theoretical SNS fraction is already substantially less favorable than in yeast. This expectation is directly supported by our efficiency-stratified analysis (Figure 3C), which shows signal at confirmed origins declining with decreasing annotated efficiency, indicating that peak-calling reliability for any given origin is tied to its firing frequency in the population, and will be correspondingly reduced in systems with less efficient or less synchronized origins. This is not a limitation specific to iSNS-seq. No origin mapping technique currently offers a demonstrated, reliable route to genome-wide origin identification in an asynchronous, otherwise untreated population of cells at this scale. Nevertheless, further development of iSNS-seq remains highly desirable because, unlike methods such as OK-seq or FORK-seq that infer initiation from perturbed cells, SNS-based approaches have the unique potential to directly capture nascent replication intermediates in unperturbed cells. Realizing that potential will require further improvements in the enrichment of authentic short nascent strands. Despite the limitations, the present work represents the first demonstration that SNS-seq can enrich for bona fide replication origins in a system where those origins are independently and confidently mapped. Furthermore, our results make clear the necessity of rigorous, origin-independent controls, and of validating any new origin-mapping method in a system where ground truth is available.

## Data availability

The raw sequencing data for this study have been deposited in the European Nucleotide Archive (ENA) at EMBL-EBI under accession number PRJEB95115.

Processed datasets, including normalized coverage tracks (bigWig) and peak calls (narrowPeak) are available through Zenodo at https://10.5281/zenodo.17781404.

The detailed iSNS-seq protocol is available at protocols.io (https://dx.doi.org/10.17504/protocols.io.kxygx4xo4l8j/v1).

## Acknowledgments

We thank Joachim Li for the kind gift of *S. cerevisiae* W303 strain.

## Author Contributions Statement

**Miiko Sokka**: Conceptualization, Formal Analysis, Investigation, Methodology, Visualization, Writing - Original Draft Preparation, Writing - Review & Editing; **John M Urban**: Conceptualization, Formal Analysis, Methodology, Visualization, Writing - Original Draft Preparation, Writing - Review & Editing; **Nicola Neretti**: Resources, Writing - Review & Editing; **Susan A Gerbi**: Conceptualization, Funding Acquisition, Resources, Supervision, Writing - Review & Editing

## References

1. Costa, A. and Diffley, J. F. X. (2022) The Initiation of Eukaryotic DNA Replication. Annu. Rev. Biochem., 91, 107–131.

2. Hu, Y. and Stillman, B. (2023) Origins of DNA Replication in Eukaryotes. Mol. Cell, 83(3), 352–372.

3. DePamphilis, M. L. (1993) Eukaryotic DNA replication: Anatomy of an origin. Annu. Rev. Biochem., 62(1), 29–63.

4. Petryk, N., Kahli, M., D’Aubenton-Carafa, Y., Jaszczyszyn, Y., Shen, Y., Silvain, M., Thermes, C., Chen, C. L., and Hyrien, O. (2016) Replication Landscape of the Human Genome. Nat. Comm., 7.

5. Wang, W., Klein, K. N., Proesmans, K., Yang, H., Marchal, C., Zhu, X., Borrman, T., Hastie, A., Weng, Z., Bechhoefer, J., Chen, C. L., Gilbert, D. M., and Rhind, N. (2021) Genome-Wide Mapping of Human DNA Replication by Optical Replication Mapping Supports a Stochastic Model of Eukaryotic Replication. Mol. Cell, 81(14), 2975–2988.e6.

6. Foss, E. J., Lichauco, C., Gatbonton-Schwager, T., Gonske, S. J., Lofts, B., Lao, U., and Bedalov, A. (2024) Identification of 1600 Replication Origins in S. Cerevisiae. eLife, 12, RP88087.

7. Urban, J. M., Foulk, M. S., Casella, C., and Gerbi, S. A. (2015) The Hunt for Origins of DNA Replication in Multicellular Eukaryotes. F1000Prime Rep., 7(30).

8. Hyrien, O. (2015) Peaks Cloaked in the Mist: The Landscape of Mammalian Replication Origins. J. Cell Biol., 208(2), 147–160.

9. Hulke, M. L., Massey, D. J., and Koren, A. (2020) Genomic Methods for Measuring DNA Replication Dynamics. Chromosome Res., 28(1), 49–67.

10. Hyrien, O., Guilbaud, G., and Krude, T. (2025) The Double Life of Mammalian DNA Replication Origins. Genes Dev., 39(5–6), 304–324.

11. Cadoret, J.-C., Meisch, F., Hassan-Zadeh, V., Luyten, I., Guillet, C., Duret, L., Quesneville, H., and Prioleau, M.-N. (2008) Genome-Wide Studies Highlight Indirect Links between Human Replication Origins and Gene Regulation. Proc. Natl. Acad. Sci. U. S. A., 105(41), 15837–15842.

12. Cayrou, C., Coulombe, P., Vigneron, A., Stanojcic, S., Ganier, O., Peiffer, I., Rivals, E., Puy, A., Laurent-Chabalier, S., Desprat, R., and Méchali, M. (2011) Genome-Scale Analysis of Metazoan Replication Origins Reveals Their Organization in Specific but Flexible Sites Defined by Conserved Features. Genome Res., 21(9), 1438–1449.

13. Besnard, E., Babled, A., Lapasset, L., Milhavet, O., Parrinello, H., Dantec, C., Marin, J. M., and Lemaitre, J. M. (2012) Unraveling Cell Type-Specific and Reprogrammable Human Replication Origin Signatures Associated with G-quadruplex Consensus Motifs. Nat. Struct. Mol. Biol., 19(8), 837–844.

14. Picard, F., Cadoret, J. C., Audit, B., Arneodo, A., Alberti, A., Battail, C., Duret, L., and Prioleau, M. N. (2014) The Spatiotemporal Program of DNA Replication Is Associated with Specific Combinations of Chromatin Marks in Human Cells. PLoS Gen., 10(5).

15. Langley, A. R., Gräf, S., Smith, J. C., and Krude, T. (2016) Genome-Wide Identification and Characterisation of Human DNA Replication Origins by Initiation Site Sequencing (Ini-Seq). Nucleic Acids Res., 44(21), 10230–10247.

16. Guilbaud, G., Murat, P., Wilkes, H. S., Lerner, L. K., Sale, J. E., and Krude, T. (2022) Determination of Human DNA Replication Origin Position and Efficiency Reveals Principles of Initiation Zone Organisation. Nucleic Acids Res., 50(13), 7436–7450.

17. Mesner, L. D., Valsakumar, V., Cieslik, M., Pickin, R., Hamlin, J. L., and Bekiranov, S. (2013) Bubble-Seq Analysis of the Human Genome Reveals Distinct Chromatin-Mediated Mechanisms for Regulating Early- and Late-Firing Origins. Genome Res., 23(11), 1774–1788.

18. Smith, D. J. and Whitehouse, I. (2012) Intrinsic Coupling of Lagging-Strand Synthesis to Chromatin Assembly. Nature, 483(7390), 434–438.

19. Müller, C. A., Boemo, M. A., Spingardi, P., Kessler, B. M., Kriaucionis, S., Simpson, J. T., and Nieduszynski, C. A. (2019) Capturing the Dynamics of Genome Replication on Individual Ultra-Long Nanopore Sequence Reads. Nat. Methods, 16(5), 429–436.

20. Hennion, M., Arbona, J.-M., Lacroix, L., Cruaud, C., Theulot, B., Tallec, B. L., Proux, F., Wu, X., Novikova, E., Engelen, S., Lemainque, A., Audit, B., and Hyrien, O. (2020) FORK-seq: Replication Landscape of the Saccharomyces Cerevisiae Genome by Nanopore Sequencing. Genome Biol., 21(1), 125.

21. Cayrou, C., Grégoire, D., Coulombe, P., Danis, E., and Méchali, M. (2012) Genome-Scale Identification of Active DNA Replication Origins. Methods, 57(2), 158–164.

22. Bielinsky, A.-K. and Gerbi, S. A. (1998) Discrete Start Sites for DNA Synthesis in the Yeast ARS1 Origin. Science, 279(5347), 95–98.

23. Lombraña, R., Álvarez, A., Fernández-Justel, J. M., Almeida, R., Poza-Carrión, C., Gomes, F., Calzada, A., Requena, J. M., and Gómez, M. (2016) Transcriptionally Driven DNA Replication Program of the Human Parasite Leishmania Major. Cell Rep., 16(6), 1774–1786.

24. Akerman, I., Kasaai, B., Bazarova, A., Sang, P. B., Peiffer, I., Artufel, M., Derelle, R., Smith, G., Rodriguez-Martinez, M., Romano, M., Kinet, S., Tino, P., Theillet, C., Taylor, N., Ballester, B., and Méchali, M. (2020) A Predictable Conserved DNA Base Composition Signature Defines Human Core DNA Replication Origins. Nat. Comm., 11(1).

25. Perkins, T. T., Dalal, R. V., Mitsis, P. G., and Block, S. M. (2003) Sequence-Dependent Pausing of Single Lambda Exonuclease Molecules. Science, 301(5641), 1914–1918.

26. van Oijen, A. M., Blainey, P. C., Crampton, D. J., Richardson, C. C., Ellenberger, T., and Xie, X. S. (2003) Single-Molecule Kinetics of λ Exonuclease Reveal Base Dependence and Dynamic Disorder. Science, 301(5637), 1235–1238.

27. Conroy, R. S., Koretsky, A. P., and Moreland, J. (2010) Lambda Exonuclease Digestion of CGG Trinucleotide Repeats. Eur. Biophys. J., 39(2), 337–343.

28. Foulk, M. S., Urban, J. M., Casella, C., and Gerbi, S. A. (2015) Characterizing and Controlling Intrinsic Biases of Lambda Exonuclease in Nascent Strand Sequencing Reveals Phasing between Nucleosomes and G-quadruplex Motifs around a Subset of Human Replication Origins. Genome Res., 125(5), 725–735.

29. Bell, S. P. and Labib, K. (2016) Chromosome Duplication in Saccharomyces Cerevisiae. Genetics, 203(3), 1027–1067.

30. Langmead, B. and Salzberg, S. L. (2012) Fast Gapped-Read Alignment with Bowtie 2. Nat. Methods, 9(4), 357–359.

31. Langmead, B., Wilks, C., Antonescu, V., and Charles, R. (2019) Scaling Read Aligners to Hundreds of Threads on General-Purpose Processors. Bioinform., 35(3), 421–432.

32. Li, H., Handsaker, B., Wysoker, A., Fennell, T., Ruan, J., Homer, N., Marth, G., Abecasis, G., Durbin, R., and 1000 Genome Project Data Processing Subgroup (2009) The Sequence Alignment/Map Format and SAMtools. Bioinform., 25(16), 2078–2079.

33. Zhang, Y., Liu, T., Meyer, C. A., Eeckhoute, J., Johnson, D. S., Bernstein, B. E., Nusbaum, C., Myers, R. M., Brown, M., Li, W., and Liu, X. S. (2008) Model-Based Analysis of ChIP-Seq (MACS). Genome Biol., 9(9), R137.

34. Robinson, J. T., Thorvaldsdóttir, H., Winckler, W., Guttman, M., Lander, E. S., Getz, G., and Mesirov, J. P. (2011) Integrative Genomics Viewer. Nat. Biotechnol., 29(1), 24–26.

35. McGuffee, S. R., Smith, D. J., and Whitehouse, I. (2013) Quantitative, Genome-Wide Analysis of Eukaryotic Replication Initiation and Termination. Mol. Cell, 50(1), 123–135.

36. Belsky, J. A., MacAlpine, H. K., Lubelsky, Y., Hartemink, J., and MacAlpine, D. M. (2015) Genome-Wide Chromatin Footprinting Reveals Changes in Replication Origin Architecture Induced by Pre-RC Assembly. Genes Dev., 29(2), 212–224.

37. Quinlan, A. R. and Hall, I. M. (2010) BEDTools: A Flexible Suite of Utilities for Comparing Genomic Features. Bioinform., 26(6), 841–842.

38. Siow, C. C., Nieduszynska, S. R., Müller, C. A., and Nieduszynski, C. A. (2012) OriDB, the DNA Replication Origin Database Updated and Extended. Nucleic Acids Res., 40(D1), D682–D686.

39. Liu, S., Liu, S., He, B., Li, L., Li, L., Wang, J., Cai, T., Chen, S., and Jiang, H. (2021) OXPHOS Deficiency Activates Global Adaptation Pathways to Maintain Mitochondrial Membrane Potential. EMBO rep., 22, e51606.

40. Comoglio, F., Schlumpf, T., Schmid, V., Rohs, R., Beisel, C., and Paro, R. (2015) High-Resolution Profiling of Drosophila Replication Start Sites Reveals a DNA Shape and Chromatin Signature of Metazoan Origins. Cell Rep., 11(5), 821–834.

41. Shumate, A. and Salzberg, S. L. (2021) Liftoff: Accurate Mapping of Gene Annotations. Bioinformatics, 37(12), 1639–1643.

42. Rodríguez-Martínez, M., Pinzón, N., Ghommidh, C., Beyne, E., Seitz, H., Cayrou, C., and Méchali, M. (2017) The Gastrula Transition Reorganizes Replication-Origin Selection in Caenorhabditis Elegans. Nature Structural and Molecular Biology, 24(3), 290–299.

43. Hage, A. E., Webb, S., Kerr, A., and Tollervey, D. (2014) Genome-Wide Distribution of RNA-DNA Hybrids Identifies RNase H Targets in tRNA Genes, Retrotransposons and Mitochondria. PLOS Genet., 10(10), e1004716.

44. Wahba, L., Costantino, L., Tan, F. J., Zimmer, A., and Koshland, D. (2016) S1-DRIP-seq Identifies High Expression and polyA Tracts as Major Contributors to R-loop Formation. Genes Dev., 30(11), 1327–1338.

45. Li, Y., Sheng, Y., Di, C., and Yao, H. (2025) Base-Pair Resolution Reveals Clustered R-loops and DNA Damage-Susceptible R-loops. Mol. Cell, 85(8), 1686–1702.e5.

46. Pratto, F., Brick, K., Cheng, G., Lam, K. W. G., Cloutier, J. M., Dahiya, D., Wellard, S. R., Jordan, P. W., and Camerini-Otero, R. D. (2021) Meiotic Recombination Mirrors Patterns of Germline Replication in Mice and Humans. Cell, 184(16), 4251–4267.e20.

47. Cayrou, C., Ballester, B., Peiffer, I., Fenouil, R., Coulombe, P., Andrau, J. C., Helden, J. V., and Méchali, M. (2015) The Chromatin Environment Shapes DNA Replication Origin Organization and Defines Origin Classes. Genome Res., 25(12), 1873–1885.

48. Singh, H., Brooke, R. G., Pausch, M. H., Williams, G. T., Trainor, C., and Dumas, L. B. (1986) Yeast DNA Primase and DNA Polymerase Activities. An Analysis of RNA Priming and Its Coupling to DNA Synthesis. J. Biol. Chem., 261(18), 8564–8569.

49. Perera, R. L., Torella, R., Klinge, S., Kilkenny, M. L., Maman, J. D., and Pellegrini, L. (2013) Mechanism for Priming DNA Synthesis by Yeast DNA Polymerase α. eLife, 2, e00482.

50. Little, J. W. (1967) An Exonuclease Induced by Bacteriophage λ. J. Biol. Chem., 242(4), 679–686.

51. Shim, J. W., Tan, Q., and Gu, L. Q. (2009) Single-Molecule Detection of Folding and Unfolding of the G-quadruplex Aptamer in a Nanopore Nanocavity. Nucleic Acids Res., 37(3), 972–982.

52. Radding, C. M. (1966) Regulation of λ Exonuclease: I. Properties of λ Exonuclease Purified from Lysogens of λT11 and Wild Type. J. Mol. Biol., 18(2), 235–250.

53. Hawkins, M., Retkute, R., Müller, C. A., Saner, N., Tanaka, T. U., de Moura, A. P. S., and Nieduszynski, C. A. (2013) High-Resolution Replication Profiles Define the Stochastic Nature of Genome Replication Initiation and Termination. Cell Rep., 5(4), 1132–1141.

54. Capra, J. A., Paeschke, K., Singh, M., and Zakian, V. A. (2010) G-Quadruplex DNA Sequences Are Evolutionarily Conserved and Associated with Distinct Genomic Features in Saccharomyces Cerevisiae. PLoS Comput. Biol., 6(7), e1000861.

55. Wu, F., Niu, K., Cui, Y., Li, C., Lyu, M., Ren, Y., Chen, Y., Deng, H., Huang, L., Zheng, S., Liu, L., Wang, J., Song, Q., Xiang, H., and Feng, Q. (2021) Genome-Wide Analysis of DNA G-quadruplex Motifs across 37 Species Provides Insights into G4 Evolution. Commun. Biol., 4(1), 98.

56. Tran, P. L. T., Mergny, J.-L., and Alberti, P. (2011) Stability of Telomeric G-quadruplexes. Nucleic Acids Res., 39(8), 3282–3294.

57. Yang, W. and Li, X. (2013) Next-Generation Sequencing of Okazaki Fragments Extracted from Saccharomyces Cerevisiae. FEBS Lett., 587(15), 2441–2447.

58. Stanojcic, S., Barckmann, B., Monsieurs, P., Crobu, L., George, S., and Sterkers, Y. (2026) Stranded Short Nascent Strand Sequencing Reveals the Topology of DNA Replication Origins in Trypanosoma Brucei. eLife, 14, RP108143.

59. Pourkarimi, E., Bellush, J. M., and Whitehouse, I. (2016) Spatiotemporal Coupling and Decoupling of Gene Transcription with DNA Replication Origins during Embryogenesis in C. Elegans. eLife, 5, e21728.

