## Supplemental file. for "Improved short nascent strand sequencing (iSNS-seq) enhances DNA replication origin detection and reduces non-origin biases"

### Supplementary Tables

**Table S1.** Sequences of $BKG_{mimic}$ and $SNS_{mimic}$.

| Spikein | sequence from 5' to 3' |
|---|---|
| BKGmimic | CGTCCAGACCCTCGCATTATAAAGGGCCGGTGGGCGGAGATTAGCGAGAGAGGATCTTTTTTCTTTTCCCCCACGCCCTCTGCTTTGGGAACCCGGGAGGGGCGCTTA<br>TGGGGAGGGTGGGGAGGGTGGGGAAGGTGGGGAGGAGACTCAGCCGGGCAGCCGAGCACTCTAGCTCTAGGATGTAAACCTCGAGAGGTCACGCTGACCTACTTGGAG<br>GTCACGCTGACCTGACTTTAGGTCACGCTGACCTAACAAGAGGTCACGCTGACCTTTAGCCAGGTCACGCTGACCTCAGTACAGGTCACGCTGACCTCTCGAGCGCGT<br>ACCAGGCTGCAGGGCGCCTCGCTAAGGCTGGGGAAAGGGCCGCGCTTTGATCAAGAGTCCCAGGGAGAGTGGAGGAAAGAAGGGTATTAATGGGCGCGCGTTCAGAGC<br>GTGGGATGTTAGTGTAGATAGGGAGGAATGATAGAGGCATAAGGAGGAAAACGATGCCTAGAATGATTAAAATAACCCAGCAACGCATTGCCACGTATACTTGGAGAG<br>CGCGTTATGAATAAACTCCCATTGCATTTGTTGGGGGGAGTCATGTATTATGCATTATGTATGCACAGCTATCTGGATTGGATACCTTCCACCCAGACTGAGTCCCCC<br>AATTTGCTGCCAAAGCAGCAGATACCGCCCCTCCTGCCCCGCCTGCTCAGGCTTCCGTGGGGCCCCGTGCGGGAGGCGTCTGTTTAGCCCTGAGATGTGTCTGCCTGT<br>TCCAGAGCTGGGCTAGGGCGAGAGGGAGGTTGCCTGCTCTCTGCCAGTCTGTACCCCACCGTCCCACCCCCACGCCTCCCACACGGAGTTCCCAATTTCTCAGCCAGG<br>TTTCAGAAGAGACAAATCCCCTTTGCGCCCTGTGGCGCCGGTTTGCAACAGTCTCGGGCTGCCGGGTTTGGGAGAAATCAAAGGTGCTAGACGGGAGAATATGGGAGG<br>GGCAGGGGGTACCCGAACCGCGGGACCGGACTTCCTAAAAGGGGCAAGTGGAGAGCTTGTGGACCGAGCCGGGGGAGTCAGCAGAGACCCTTGTGAAAAAAACCGTTA<br>ACTCTTTCCTCCCCGGACAAACCGGACGTTTAATTCCTTTCCAGGTCCTCTTTCCCCTTTTATTATTGGAAATGCGGTCATGCACAA |
| SNSmimic | CCGAGTTTGCTAGGCACTGATACATAACTCTTTTCCAATAATTGGGGAAGTCATTCAAATCTATAATAGGTTTCAGATTTGCTTCAATAAATTCTGACTGTAGCTGCT<br>GAAACGTTGCGGTTGAACTATATTTCCTTATAACTTTTACGAAAGAGTTTCTTTGAGTAATCACTTCACTCAAGTGCTTCCCTGCCTCCAAACGATACCTGTTAGCAA<br>TATTTAATAGCTTGAAATGATGAAGAGCTCTGTGTTTGTCTTCCTGCCTCCAGTTCGCCGGGCATTCAACATAAAAACTGATAGCACCCGGAGTTCCGGAAACGAAAT<br>TTGCATATACCCATTGCTCACGAAAAAAAATGTCCTTGTCGATATAGGGATGAATCGCTTGGTGTACCTCATCTACTGCGAAAACTTGACCTTTCTCTCCCATATTGC<br>AGTCGCGGCACGATGGAACTAAATTAATAGGCATCACCGAAAATTCAGGATAATGTGCAATAGGAAGAAAATGATCTATATTTTTTGTCTGTCCTATATCACCACAAA<br>ATGGACATTTTTCACCTGATGAAACAAGCATGTCATCGTAATATGTTCTAGCGGGTTTGTTTTTATCTCGGAGATTATTTTCATAAAGCTTTTCTAATTTAACCTTTG<br>TCAGGTTACCAACTACTAAGGTTGTAGGCTCAAGAGGGTGTGTCCTGTCGTAGGTAAATAACTGACCTGTCGAGCTTAATATTCTATATTGTTGTTCTTTCTGCAAAA<br>AAGTGGGGAAGTGAGTAATGAAATTATTTCTAACATTTATCTGCATCATACCTTCCGAGCATTTATTAAGCATTTCGCTATAAGTTCTCGCTGGAAGAGGTAGTTTTT<br>TCATTGTACTTTACCTTCATCTCTGTTCATTATCATCGCTTTTAAAACGGTTCGACCTTCTAATCCTATCTGACCATTATAATTTTTTAGAATGGTTTCATAAGAAAG<br>CTCTGAATCAACGGACTGCGATAATAAGTGGTGGTATCCAGAATTTGTCACTTCAAGTAAAAACACCTCACGAGTTAAAACACCTAAGTTCTCACCGAATGTCTCAAT<br>ATCCGGACGGATAATATTTATTGCTTCTCTTGACCGTAGGACTTTCCACATGCAGGATTTTGGAACCTCTTGCAGTACTACTGGGGAATGAGTTGCAATTATTGCTAC<br>ACCATTGCGTGCATCG |

**Table S2.** All peak summits were extended by $\pm 500$ bp, and sensitivity (SN), positive predictive value (PPV), and F-score were computed relative to confirmed origins from oriDB (38). Statistical significance is indicated as follows: $p < 0.05$ (*), $p < 0.01$ (**), $p < 0.001$ (***)

| | SN | | | PPV | | | F-score | |
|---|---|---|---|---|---|---|---|---|
| Sample | Observed | Expected | P-value | Observed | Expected | P-value | Observed | Expected |
| iSNS rep1 | 0.33 | 0.05 | *** | 0.34 | 0.05 | *** | 0.34 | 0.05 |
| iSNS rep2 | 0.29 | 0.05 | *** | 0.30 | 0.05 | *** | 0.29 | 0.05 |
| tSNS rep2 | 0.06 | 0.05 | – | 0.06 | 0.05 | – | 0.06 | 0.05 |
| tSNS rep1 | 0.08 | 0.05 | * | 0.08 | 0.05 | * | 0.08 | 0.05 |
| ORC rep1 | 0.41 | 0.05 | *** | 0.43 | 0.05 | *** | 0.42 | 0.05 |
| ORC rep2 | 0.40 | 0.04 | *** | 0.54 | 0.05 | *** | 0.46 | 0.05 |
| OK rep1 | 0.55 | 0.05 | *** | 0.57 | 0.05 | *** | 0.56 | 0.05 |
| OK rep2 | 0.56 | 0.05 | *** | 0.59 | 0.05 | *** | 0.57 | 0.05 |
| FORK | 0.66 | 0.05 | *** | 0.68 | 0.05 | *** | 0.67 | 0.05 |
| shuffle | 0.05 | 0.05 | – | 0.05 | 0.05 | – | 0.05 | 0.05 |

**Table S3.** All iSNS-seq and tSNS-seq peak summits were extended by ±500bp, and sensitivity (SN), positive predictive value (PPV), and F-score were computed relative to highly expressed genes, tRNAs and snoRNAs. Statistical significance is indicated as follows: $p < 0.05$ (*), $p < 0.01$ (**), $p < 0.001$ (***).

| | SN | | | PPV | | | F-score | |
|---|---|---|---|---|---|---|---|---|
| Sample | Observed | Expected | P-value | Observed | Expected | P-value | Observed | Expected |
| tSNS rep1 | 0.76 | 0.12 | *** | 0.29 | 0.05 | *** | 0.42 | 0.07 |
| tSNS rep2 | 0.81 | 0.06 | *** | 0.67 | 0.05 | *** | 0.73 | 0.05 |
| iSNS rep1 | 0.05 | 0.02 | *** | 0.11 | 0.05 | *** | 0.07 | 0.03 |
| iSNS rep2 | 0.04 | 0.03 | – | 0.06 | 0.05 | – | 0.05 | 0.04 |
| shuffle | 0.15 | 0.12 | – | 0.06 | 0.05 | – | 0.09 | 0.07 |

### Supplementary Figures

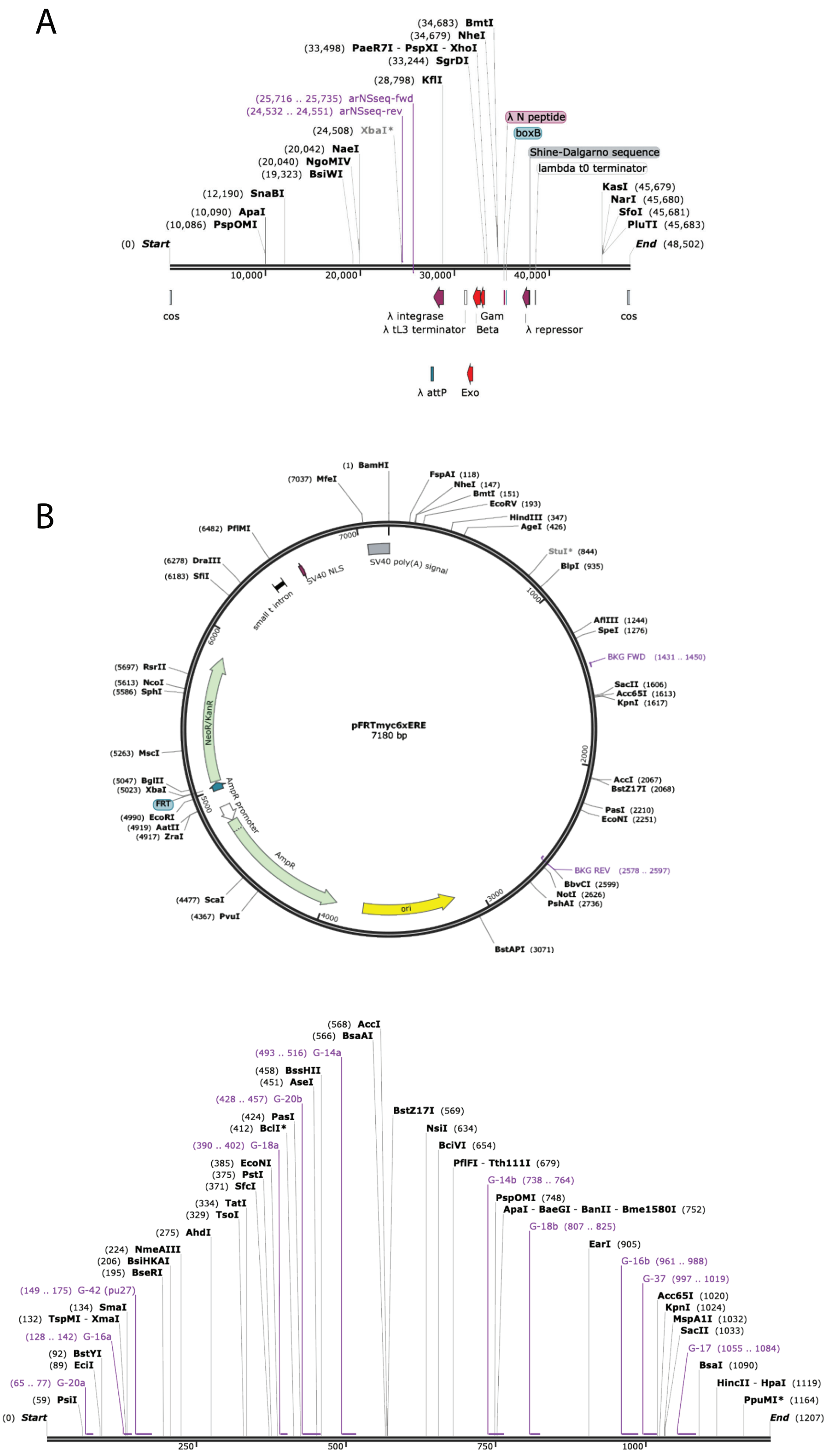


Fig. S1: DNA template maps used to generate SNS$^{mimic}$ and BKG$^{mimic}$. (A) $\lambda$ DNA map showing the forward (arNSseq-fwd) and reverse (arNSseq-rev) primer locations for generating SNS$^{mimic}$. (B) pFRTmyc6xERE plasmid map (on top) from (28) and the forward (BKG FWD) and reverse (BKG REV) primer locations for generating BKG$^{mimic}$. The panel on bottom shows the full BKG$^{mimic}$ with putative G4 motifs, including the much studied pu27 G4. All sequence maps and the annotations were generated using SnapGene Viewer.

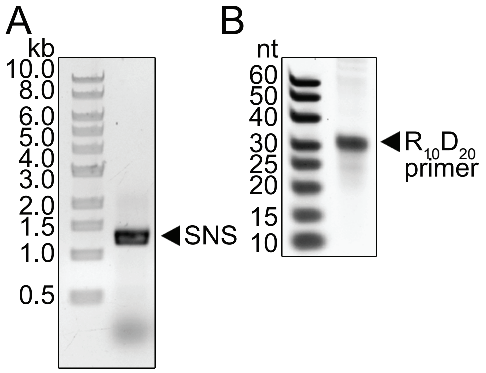


Fig. S2: The RNA primer is preserved during production of the artificial SNS. (A) The PCR conditions efficiently amplify the 1.2 kb SNS (AGE). (B) A subsample from A was run in PAGE to examine if the primers survived the PCR. The lack of 20 nt band shows that the 30 nt long primer that contains 10 nt 5' RNA survived the PCR without degradation.

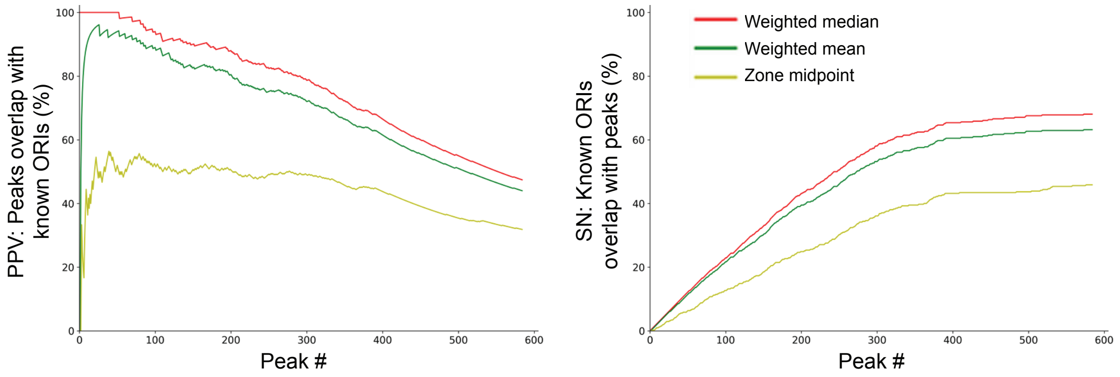


Fig. S3: Comparison of approaches to call "peak summits" (aka preferred initiation sites within broader initiation zones) in FORK-seq dataset (20). The initiation site midpoints (ISM) were aggregated into zones and the 'peak summits' were called either using weighted mean or median of all ISMs within the zones, or by taking the midpoint of the zones (equidistant between the minimum and maximum ISMs). All 'peak summits' were extended 500 bp up- and downstream. Sensitivity (SN) and positive predictive value (PPV) plots as a function of peak confidence (see Methods) overlapping with confirmed origins.

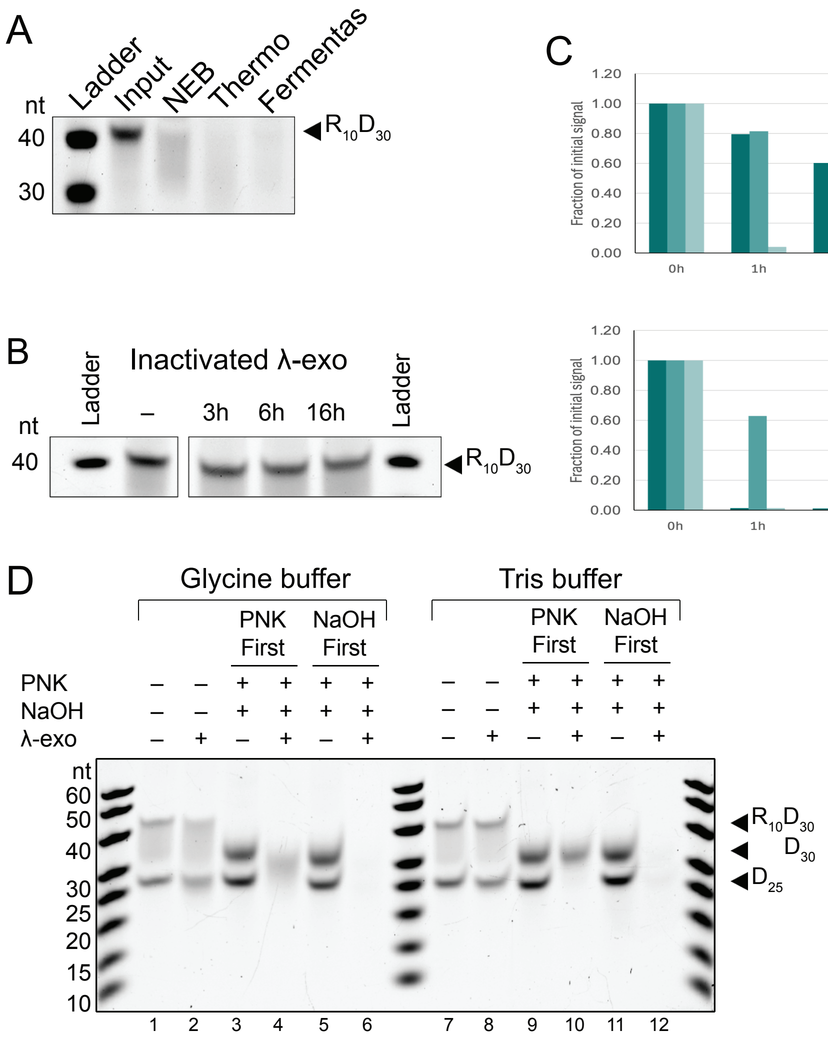


Fig. S4: Controls for λ-exo. (A) $R_{10}D_{30}$ is digested by all tested λ-exo enzymes from different suppliers. $R_{10}D_{30}$ was subjected to a 16 h digestion by λ-exo from either New England Biolabs, ThermoFisher or Fermentas (old custom preparation, which is now discontinued). (B) RNA digestion is not due to contaminating RNase activity in the reaction. Phosphorylated $R_{10}D_{30}$ was incubated with heat-inactivated λ-exo for the indicated times. (C) Semi-quantification of λ-exo digestion kinetics from Figure 1C. Band intensities for the RNA-DNA chimera ($R_{10}D_{30}$), G4 oligo ($D_{10}G4_{27}D_{23}$), and all-DNA control ($D_{25}$) were quantified by densitometry and normalized to the $D_{20}$ loading control at each timepoint, then expressed as a fraction of the initial (t=0) signal. Bar plots show normalized band intensity at each timepoint of the λ-exo digestion time course in Glycine-KOH (left panel) and Tris-HCl buffers (right panel). (D) Two oligos, $R_{10}D_{30}$ and $D_{25}$ were subjected in the same reaction to phosphorylation (PNK) and hydrolysis (NaOH) in indicated order in either Glycine-KOH or Tris-HCl buffer and then subjected to λ-exo digestion for 16 h.

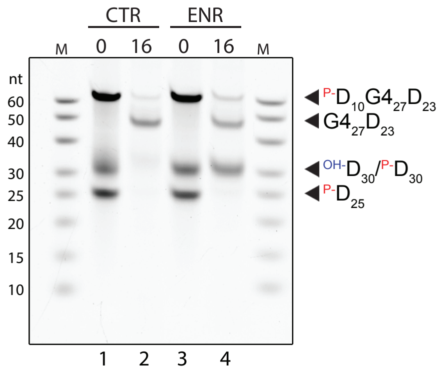


Fig. S5:

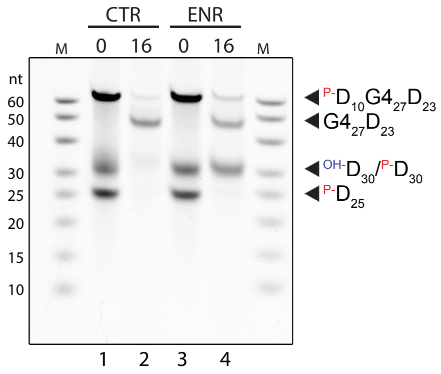


Fig. S6: The same experiment as in Figure 1E performed in glycine-KOH buffer. Briefly, 30 pmol of each oligo was subjected to 0 h or 16 h of $\lambda$-exo digestion, as indicated. See Figure 1D for the experimental setup. The picture is from the same gel as in Figure 1E, and the right marker (M) lane is the same as the left marker lane.

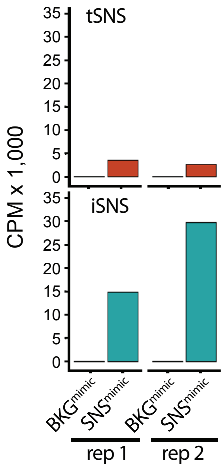


Fig. S7: Normalized read counts (CPM) of SNS$^{\mathrm{mimic}}$ and BKG$^{\mathrm{mimic}}$ spike-ins in the two replicates of iSNS-seq and tSNS-seq.

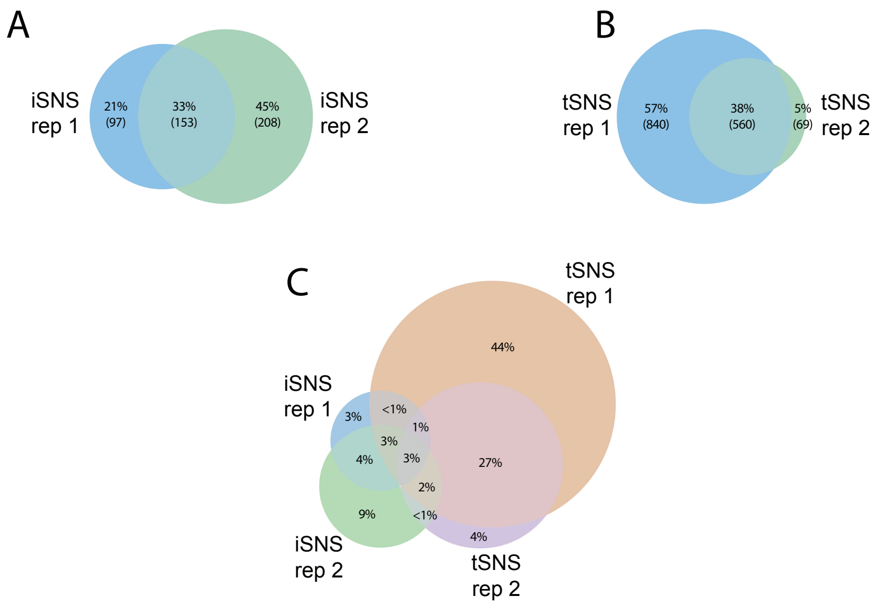


Fig. S8: Euler diagrams of intersecting peaks between (A) iSNS-seq replicates, (B) tSNS-seq replicates, and (C) all four datasets.

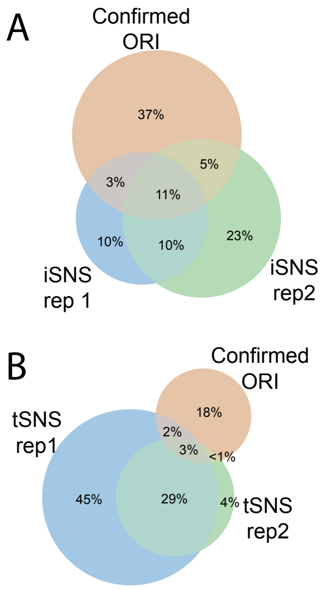


Fig. S9: Euler diagrams showing overlap of called peaks across iSNS-seq (top) or tSNS-seq (bottom) replicates and their intersection with confirmed origins of replication (ORIs) from oriDB. Peaks were considered overlapping if within 1 bp. Percentages represent the fraction of the total combined peaks across all three sets being compared (i.e., the union of iSNS-seq rep 1, iSNS-seq rep 2, and confirmed ORIs for panel A; tSNS-seq rep 1, tSNS-seq rep 2, and confirmed ORIs for panel B) that fall into each region of the diagram.

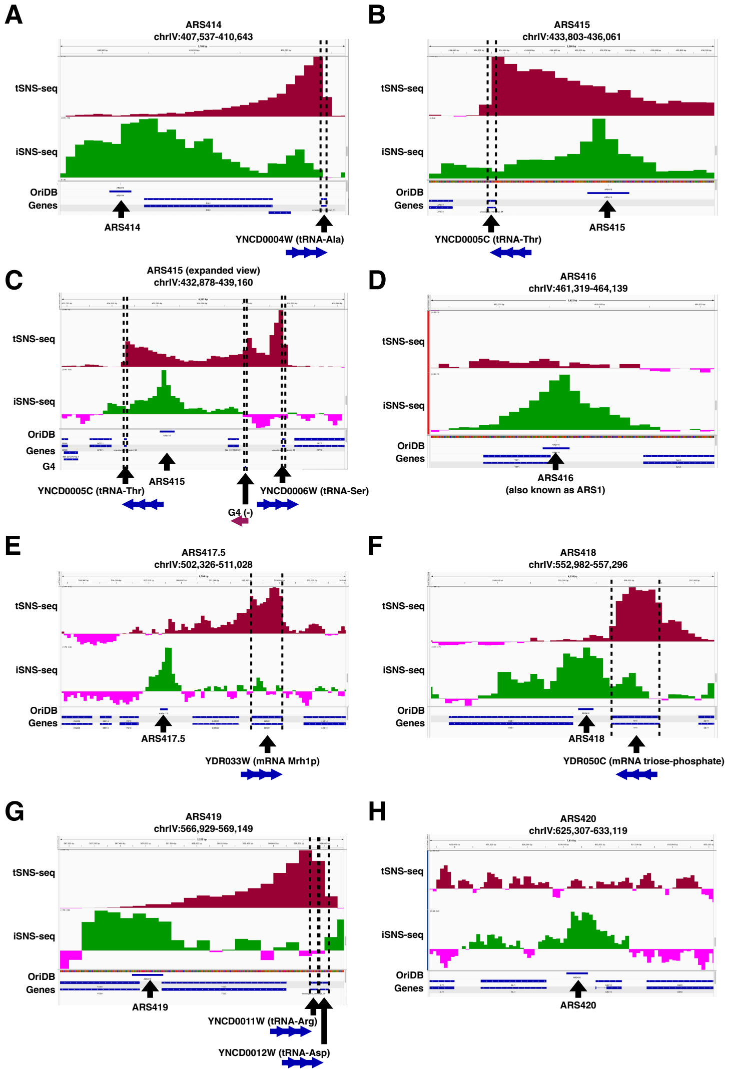


Fig. S10: iSNS-seq signal is symmetrically enriched around known replication origins whereas tSNS-seq signal is asymmetric, not enriched over origins, and is correlated with nearby genes. This figure shows close up views in the 2-9 kb range of the known replication origins (autonomously replicating sequences, ARS) according to OriDB at the locus highlighted in Figure 3D (and Figure 5A). In all IGV traces, from top to bottom, tSNS-seq signal, iSNS-seq signal, locations of OriDB origins, and the locations of genes are shown. When a G4 motif was also at the locus, it is shown below the genes track. The iSNS-seq and tSNS-seq are presented as GC-content-informed robust Z-scores (as described below) with scores below the median in pink, and scores above the median in maroon for tSNS-seq (top track) and green for iSNS-seq (second from top). (A) ARS414, (B) ARS415, (C) ARS415 with broader view, (D) ARS416 (also known as ARS1), (E) ARS417.5, (F) ARS418, (G) ARS419, (H) ARS420. Note that asymmetric tSNS-seq enrichments at these loci are mostly at nearby tRNA genes with the exceptions of ARS416 and ARS420 for which there is no nearby tSNS-seq enrichments, and ARS17.5 and ARS418 where the nearby tSNS-seq enrichments are associated with mRNA genes: Mrh1 (YDR033W) and triose-phosphate isomerase (YDR050C), respectively. **Detailed methods**: Concordant fragment coverage was computed from BAM files and sorted with BEDtools (bedtools genomecov -bga -pc -g *genomefile* -ibam *bamfile* — sortBed -i -) to produce bedGraph files, which were converted to bigWig with bedGraphToBigWig. BEDtools was used to break the genome up into 100 bp bins (makewindows -w 100 -s 100). BEDtools (nucBed) was used to get the GC content in each bin. For each sample, the average fragment coverage in each bin was computed using the above bigWigs with bigWigAverageOverBed to produce a 100-bp step coverage bedGraph, and the coverage and GC content bedGraphs were joined into a 5-column table file (chr, start, end, GC content, coverage). A custom GC-content-informed median-normalization pipeline, named "gecko", was run on the table file (`https://github.com/JohnUrban/geckocorrection`). Briefly, for a given 5-column table file, the 100 bp bins were separated into groups based on their integer GC content level from 0 to 100. The median coverage and the median absolute deviation (MAD) from that median for each GC content level was learned. Then raw coverage levels were converted to GC-informed MAD units by subtracting the median coverage and dividing by the associated MAD given the GC content of each bin: (x_gc - median_gc)/MAD_gc. Thus, the signal in each bin is presented in units of median absolute deviations from the median coverage for the GC content in that bin. MAD units are analogous to robust Z-scores, but are not multiplied by a constant to approximate standard deviations. For each bin, the average MAD unit score across replicates was used. All code referring to this figure can be found at (`https://github.com/JohnUrban/iSNS-seq`).

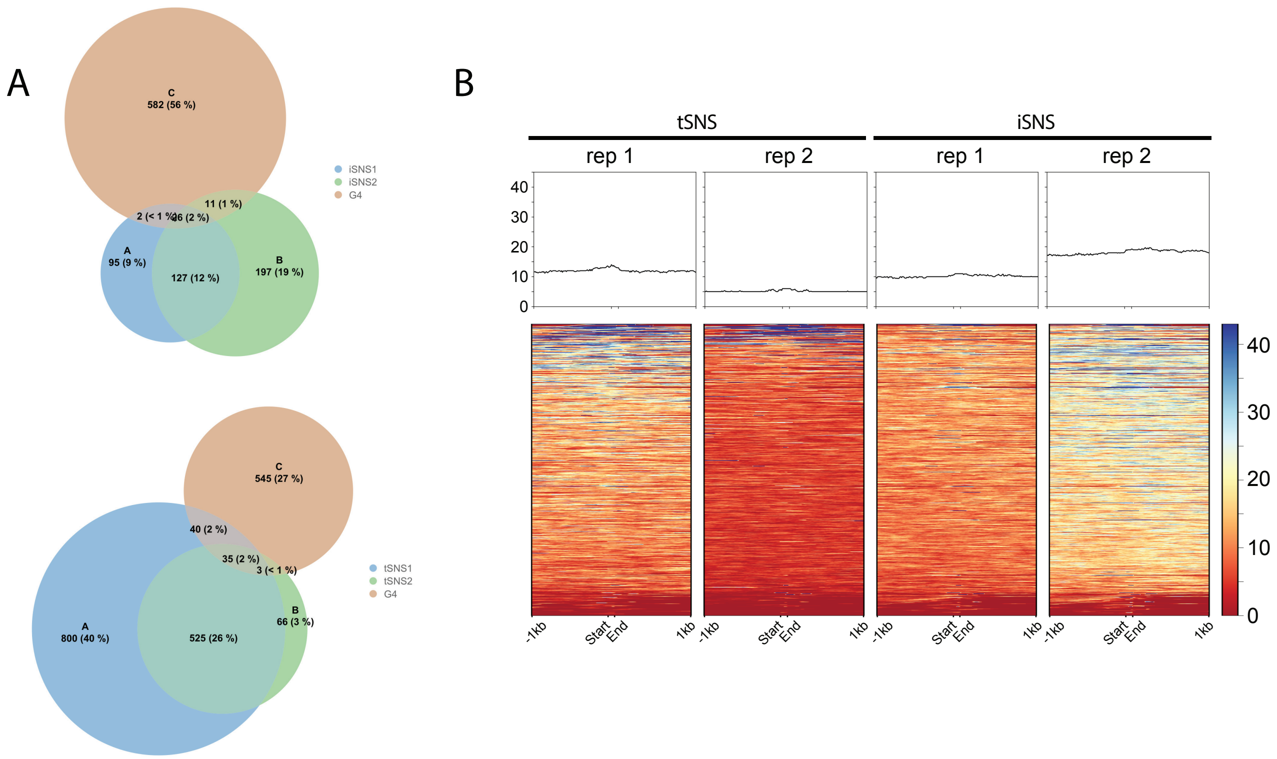


Fig. S11: Euler diagrams of intersecting G4 motifs (54) and (A) iSNS-seq peaks (top) and tSNS-seq peaks (bottom). (B) Heatmaps showing pileup of iSNS-seq and tSNS-seq signal around G4 motifs, spanning ±1 kb from start and end of the motifs. Median signal profiles are shown above each heatmap.

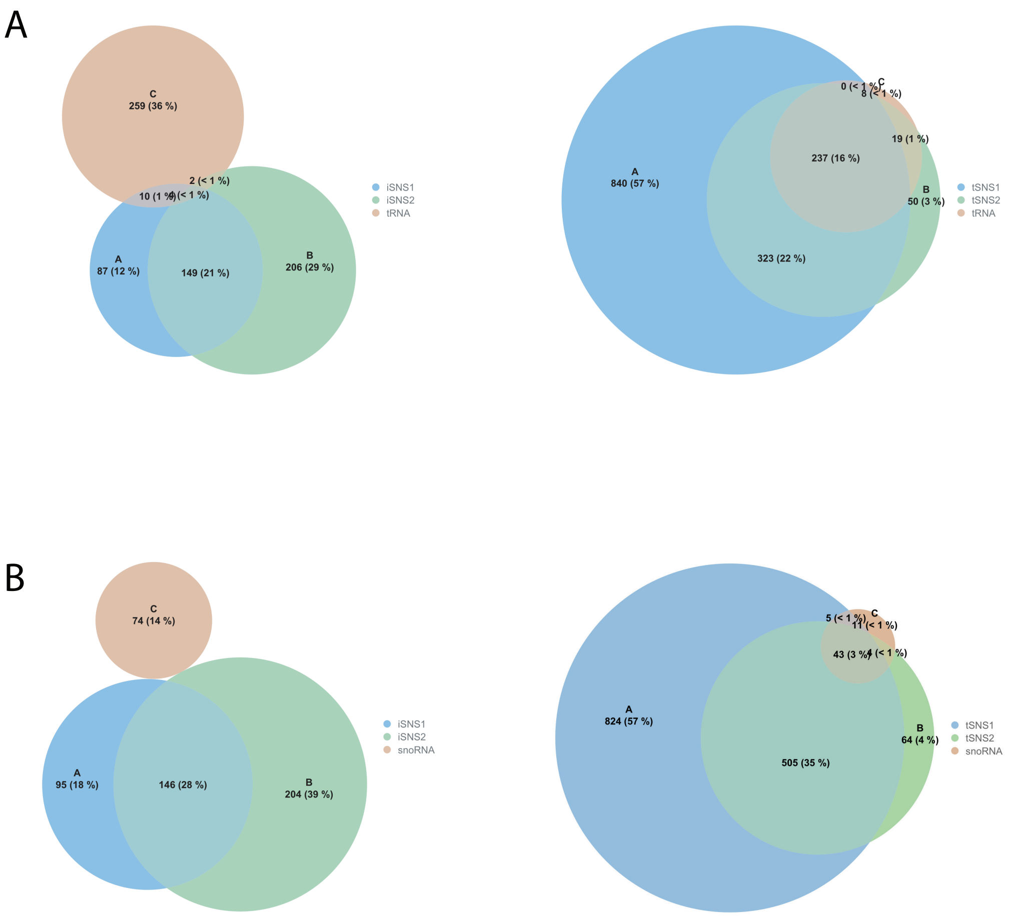


Fig. S12: Euler diagrams of intersecting peaks between (A) tRNAs or (B) snoRNAs and iSNS-seq (left) or tSNS-seq (right).

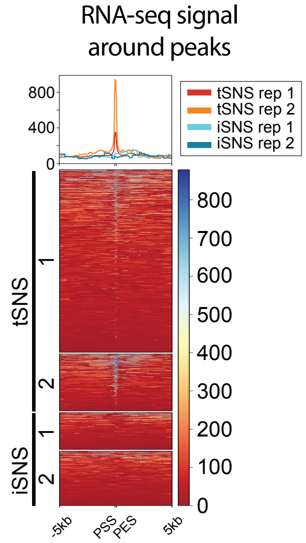


Fig. S13: Poly(A)-derived RNA-seq signal (39) around iSNS- and tSNS-seq peak regions (PSS, peak start site; PES, peak end site), shown as heatmaps ±5 kb from peak boundaries, with median signal above.

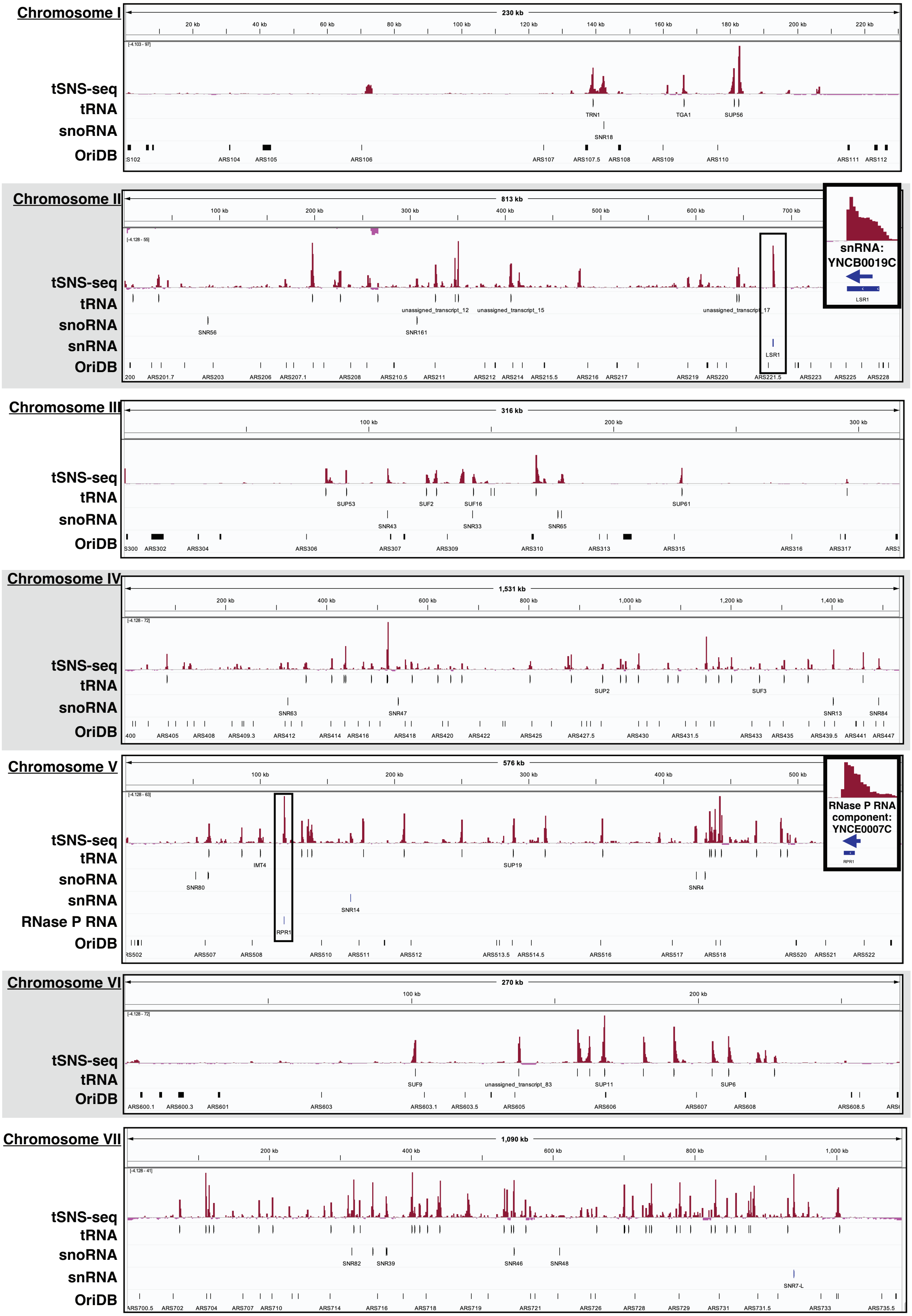
Chromosome I
230 kb
tSNS-seq
tRNA
snoRNA
OriDB
TRN1
TGA1
SUP56
SNR18
Chromosome II
813 kb
tSNS-seq
tRNA
snoRNA
snRNA
OriDB
unassigned_transcript_12
unassigned_transcript_15
unassigned_transcript_17
SNR56
SNR161
LSR1
snRNA: YNCB0019C
Chromosome III
316 kb
tSNS-seq
tRNA
snoRNA
OriDB
SUP53
SUF2
SUF16
SUP61
SNR43
SNR33
SNR65
Chromosome IV
1,531 kb
tSNS-seq
tRNA
snoRNA
OriDB
SUP2
SUF3
SNR63
SNR47
SNR13
SNR84
Chromosome V
576 kb
tSNS-seq
tRNA
snoRNA
snRNA
RNase P RNA
OriDB
IMT4
SUP19
SNR80
SNR4
SNR14
RPR1
RNase P RNA component: YNCE0007C
Chromosome VI
270 kb
tSNS-seq
tRNA
OriDB
SUF9
unassigned_transcript_83
SUP11
SUP6
Chromosome VII
1,090 kb
tSNS-seq
tRNA
snoRNA
snRNA
OriDB
SNR82
SNR39
SNR46
SNR48
SNR7-L

Fig. S14: Visual correlation of prominent tSNS-seq peaks with tRNA genes (and other types of non-coding RNA genes) across the entire lengths of chromosomes I, II, III, IV, V, VI, and VII. Although most prominent peaks correspond with tRNA genes, followed by snoRNA genes, other non-coding RNA genes (as well as mRNA genes) also demonstrate the same effect, with the gene position and orientation following the same trend as tRNA genes with respect to the tSNS-seq asymmetric enrichment profile. Chromosome II highlights an snRNA gene example, and chromosome V highlights the gene for the RNA component of RNase P. The tSNS-seq signal in each bin is presented as GC-content-informed MAD units, which are units of median absolute deviations from the median coverage for the GC content in that bin (as described in the Detailed methods section of the legend for Supplementary Figure S8). Scores below the median are pink, and scores above the median are maroon.

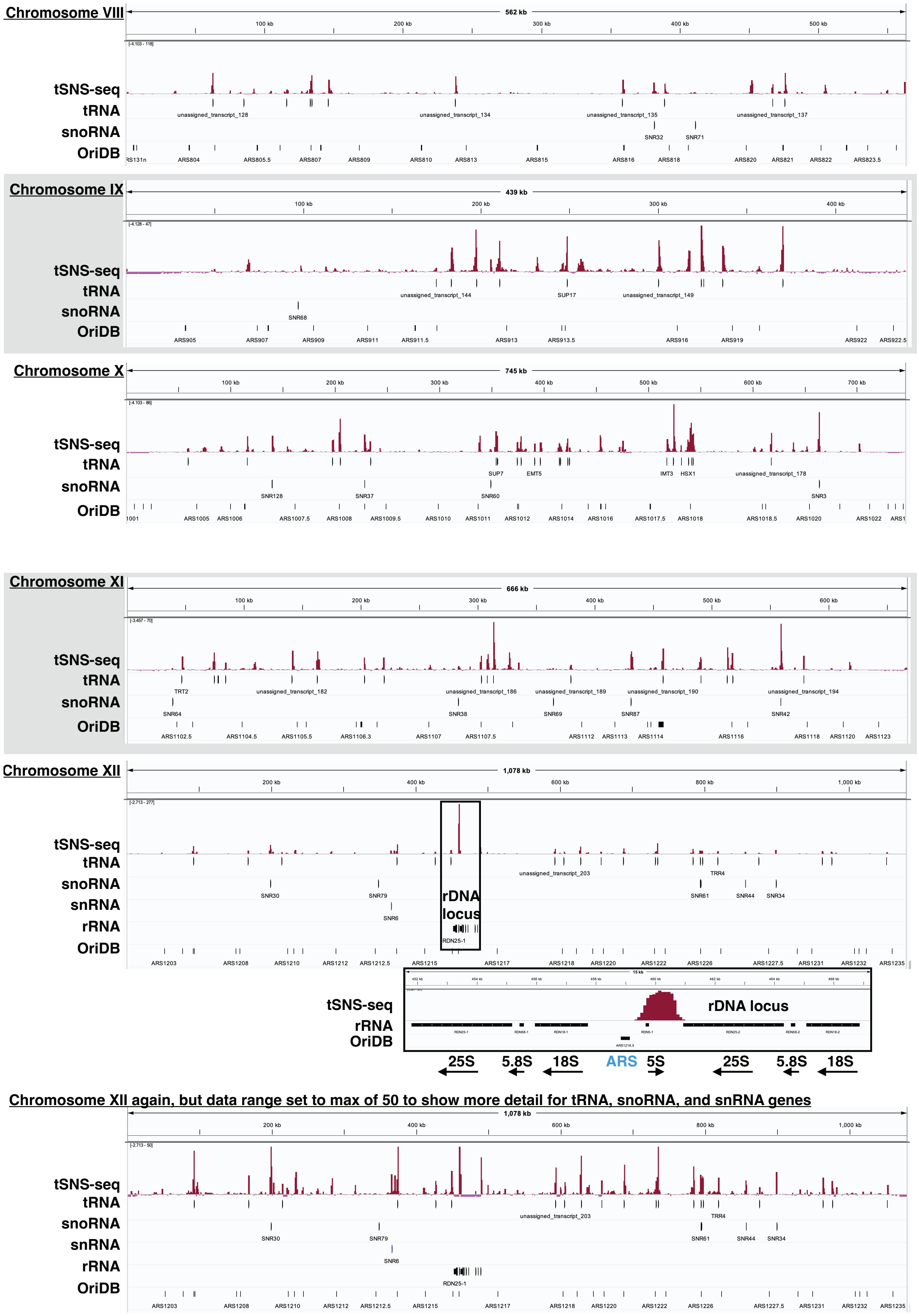

Chromosome VIII
562 kb
100 kb
200 kb
300 kb
400 kb
500 kb
tSNS-seq
tRNA
snoRNA
OriDB
unassigned_transcript_128
unassigned_transcript_134
unassigned_transcript_135
unassigned_transcript_137
SNR32
SNR71
RS131n ARS804 ARS805.5 ARS807 ARS809 ARS810 ARS813 ARS815 ARS816 ARS818 ARS820 ARS821 ARS822 ARS823.5
Chromosome IX
439 kb
100 kb
200 kb
300 kb
400 kb
tSNS-seq
tRNA
snoRNA
OriDB
unassigned_transcript_144
SUP17
unassigned_transcript_149
SNR68
ARS905 ARS907 ARS909 ARS911 ARS911.5 ARS913 ARS913.5 ARS916 ARS919 ARS922 ARS922.5
Chromosome X
745 kb
100 kb
200 kb
300 kb
400 kb
500 kb
600 kb
700 kb
tSNS-seq
tRNA
snoRNA
OriDB
SUP7
EMT5
IMT3
HSX1
unassigned_transcript_178
SNR128
SNR37
SNR60
SNR3
1001 ARS1005 ARS1006 ARS1007.5 ARS1008 ARS1009.5 ARS1010 ARS1011 ARS1012 ARS1014 ARS1016 ARS1017.5 ARS1018 ARS1018.5 ARS1020 ARS1022 ARS1
Chromosome XI
666 kb
100 kb
200 kb
300 kb
400 kb
500 kb
600 kb
tSNS-seq
tRNA
snoRNA
OriDB
TRT2
unassigned_transcript_182
unassigned_transcript_186
unassigned_transcript_189
unassigned_transcript_190
unassigned_transcript_194
SNR64
SNR38
SNR69
SNR87
SNR42
ARS1102.5 ARS1104.5 ARS1105.5 ARS1106.3 ARS1107 ARS1107.5 ARS1112 ARS1113 ARS1114 ARS1116 ARS1118 ARS1120 ARS1123
Chromosome XII
1,078 kb
200 kb
400 kb
600 kb
800 kb
1,000 kb
tSNS-seq
tRNA
snoRNA
snRNA
rRNA
OriDB
unassigned_transcript_203
TRR4
SNR30
SNR79
SNR6
SNR61
SNR44
SNR34
rDNA locus
RDN25-1
ARS1203 ARS1208 ARS1210 ARS1212 ARS1212.5 ARS1215 ARS1217 ARS1218 ARS1220 ARS1222 ARS1226 ARS1227.5 ARS1231 ARS1232 ARS1235
15 kb
tSNS-seq
rRNA
OriDB
rDNA locus
25S
5.8S
18S
ARS
5S
25S
5.8S
18S
Chromosome XII again, but data range set to max of 50 to show more detail for tRNA, snoRNA, and snRNA genes
1,078 kb
200 kb
400 kb
600 kb
800 kb
1,000 kb
tSNS-seq
tRNA
snoRNA
snRNA
rRNA
OriDB
unassigned_transcript_203
TRR4
SNR30
SNR79
SNR6
SNR61
SNR44
SNR34
RDN25-1
ARS1203 ARS1208 ARS1210 ARS1212 ARS1212.5 ARS1215 ARS1217 ARS1218 ARS1220 ARS1222 ARS1226 ARS1227.5 ARS1231 ARS1232 ARS1235

Fig. S15: Visual correlation of prominent tSNS-seq peaks with tRNA genes (and other types of non-coding RNA genes) across the entire lengths of chromosomes VIII, IX, X, XI, and XII. The most extreme tSNS-seq enrichment on chromosome XII is at the rDNA locus above the 5S gene at its left end, and is between the 5S and 25S genes. However, the position and orientation of the 5S gene does not follow the same pattern seen for tRNA genes. Moreover, the enrichment is not asymmetric. Although it is symmetric, it is not over the known replication origin at the rDNA locus (ARS1216.5). Note that the iSNS-seq signal (not shown) also shows the exact same enrichment here. In contrast, an orthogonal origin-mapping technique (FORK-seq) shows an enrichment of single-molecule initiation events exactly over the origin at this locus. It is unclear at the moment what causes this enrichment common to both tSNS-seq and iSNS-seq. Chromosome XII is shown a second time (bottom-most image) with the data range capped to better show the correlation of tSNS-seq signal with tRNA (and other non-coding) genes on this chromosome. The tSNS-seq signal in each bin is presented as GC-content-informed MAD units, which are units of median absolute deviations from the median coverage for the GC content in that bin (as described in the Detailed methods section of the legend for Supplementary Figure S8). Scores below the median are pink, and scores above the median are maroon.

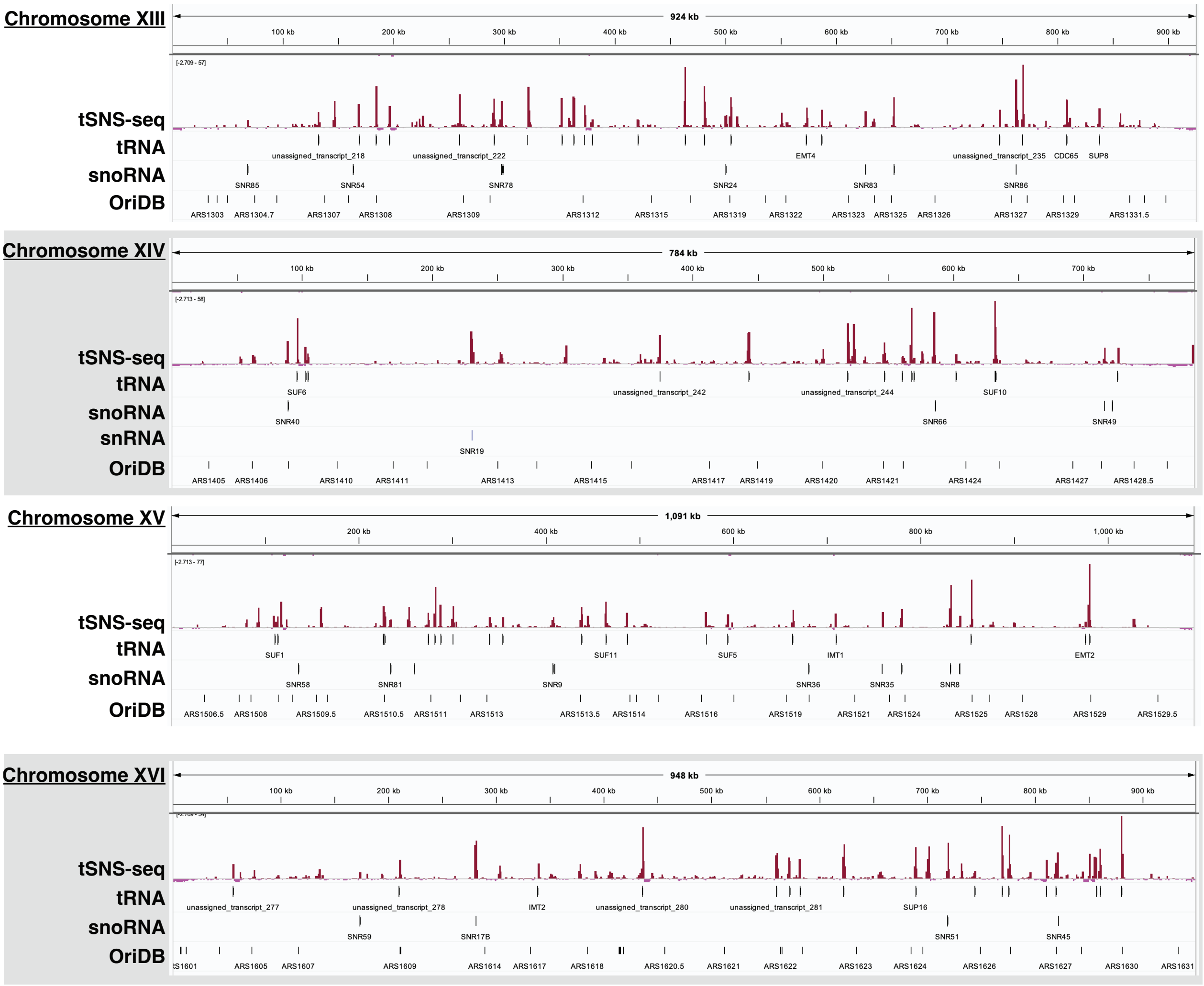


Fig. S16: Visual correlation of prominent tSNS-seq peaks with tRNA genes (and other types of non-coding RNA genes) across the entire lengths of chromosomes XIII, XIV, XV, and XVI. The tSNS-seq signal in each bin is presented as GC-content-informed MAD units, which are units of median absolute deviations from the median coverage for the GC content in that bin (as described in the Detailed methods section of the legend for Supplementary Figure S8). Scores below the median are pink, and scores above the median are maroon.

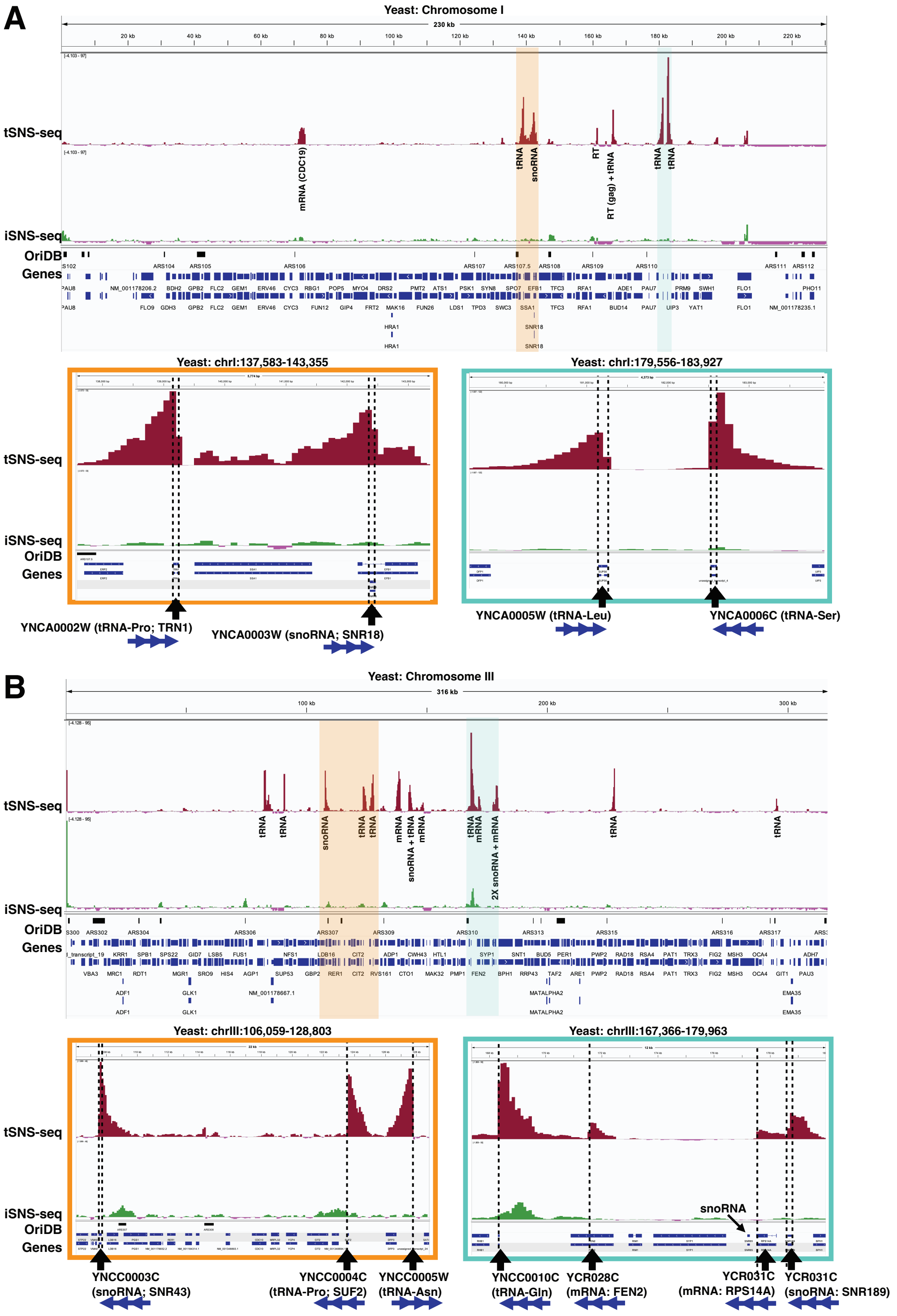


Fig. S17: Examples of the most prominent tSNS-seq peaks on chromosomes I and III. The most prominent tSNS-seq peaks are correlated mostly with tRNA and snoRNA genes, but also mRNA genes. (A) A 230 kb locus with the most prominent tSNS-seq peaks on chromosome I. (B) A 316 kb locus with the most prominent tSNS-seq peaks on chromosome III. In both, the regions of the locus highlighted in light orange or light blue are shown zoomed in the sub-locus images below that are bordered with light orange and light blue, respectively. Chromosome II and the chromosome III locus also highlights this effect, albeit not as extreme, at two mRNA genes: FEN2 (YCR028C) and RPS14A (YCR031C). The tSNS-seq signal in each bin is presented as GC-content-informed MAD units, which are units of median absolute deviations from the median coverage for the GC content in that bin (as described in the Detailed methods section of the legend for Supplementary Figure S8). Scores below the median are pink, and scores above the median are maroon.

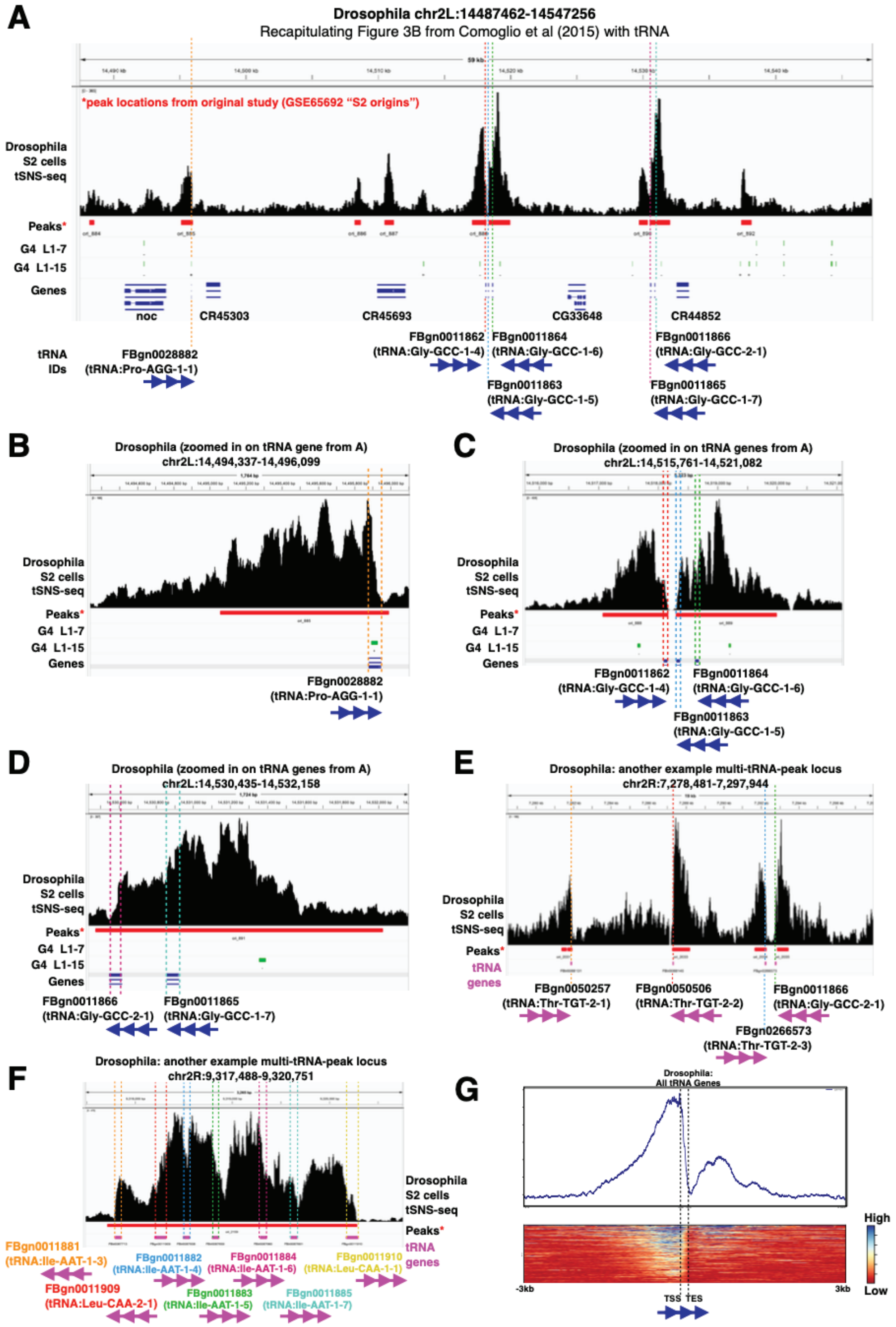


Fig. S18: (A) Recapitulating 'figure 3B' from Comoglio *et al.* (40), but now highlighting the previously-unrecognized presence of tRNA genes at this locus that are responsible for the obstruction-shaped peaks. (B) Zoom in on left-most tRNA gene from A. (C) Zoom in on central three tRNA genes from A. (D) Zoom in on right-most tRNA genes from A. (E) Another example locus with multiple tRNA genes causing obstruction shaped peaks. (F) A third example locus of multiple tRNA genes causing obstruction shaped peaks. (G) An aggregate profile and heatmap of the tSNS-seq signal around all tRNA genes.

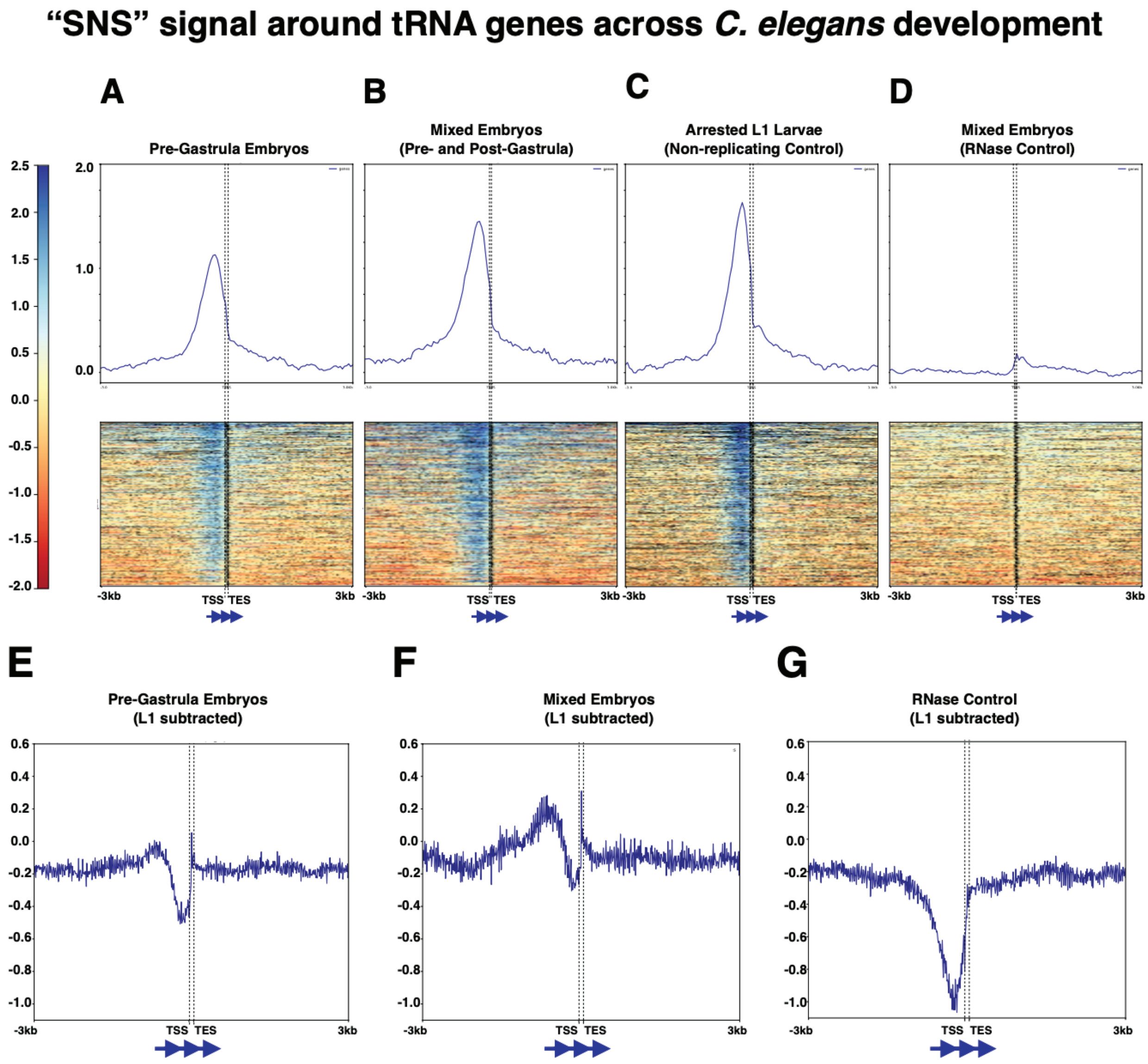


Fig. S19: The tRNA bias shows up in the published *C. elegans* tSNS-seq data (42). Median-normalized $\log_2$(Cy5/Cy3) tSNS microarray signal around all tRNA genes using tSNS samples from pre-gastrula embryos, mixed embryos, arrested L1 larvae that serves as a non-replicating control, and the RNase control of mixed stage embryos, where the samples were treated with RNase prior to the tSNS procedure (top row). Pre-computed $\log_2$(Cy5/Cy3) ratios that were subject to normalization, L1 larval control background subtraction, and smoothing by the original authors (bottom row). Since the tRNA gene bias is present in the L1 non-replicating control, subtracting out the L1 signal from the other samples somewhat remediates the tRNA-gene bias. However, the non-replicating control will ultimately have a different transcriptional landscape that can't account for this bias everywhere it occurs in the target sample. Moreover, it does not appear to fully account for the signal even at the tRNA genes, perhaps from differential tRNA abundances.

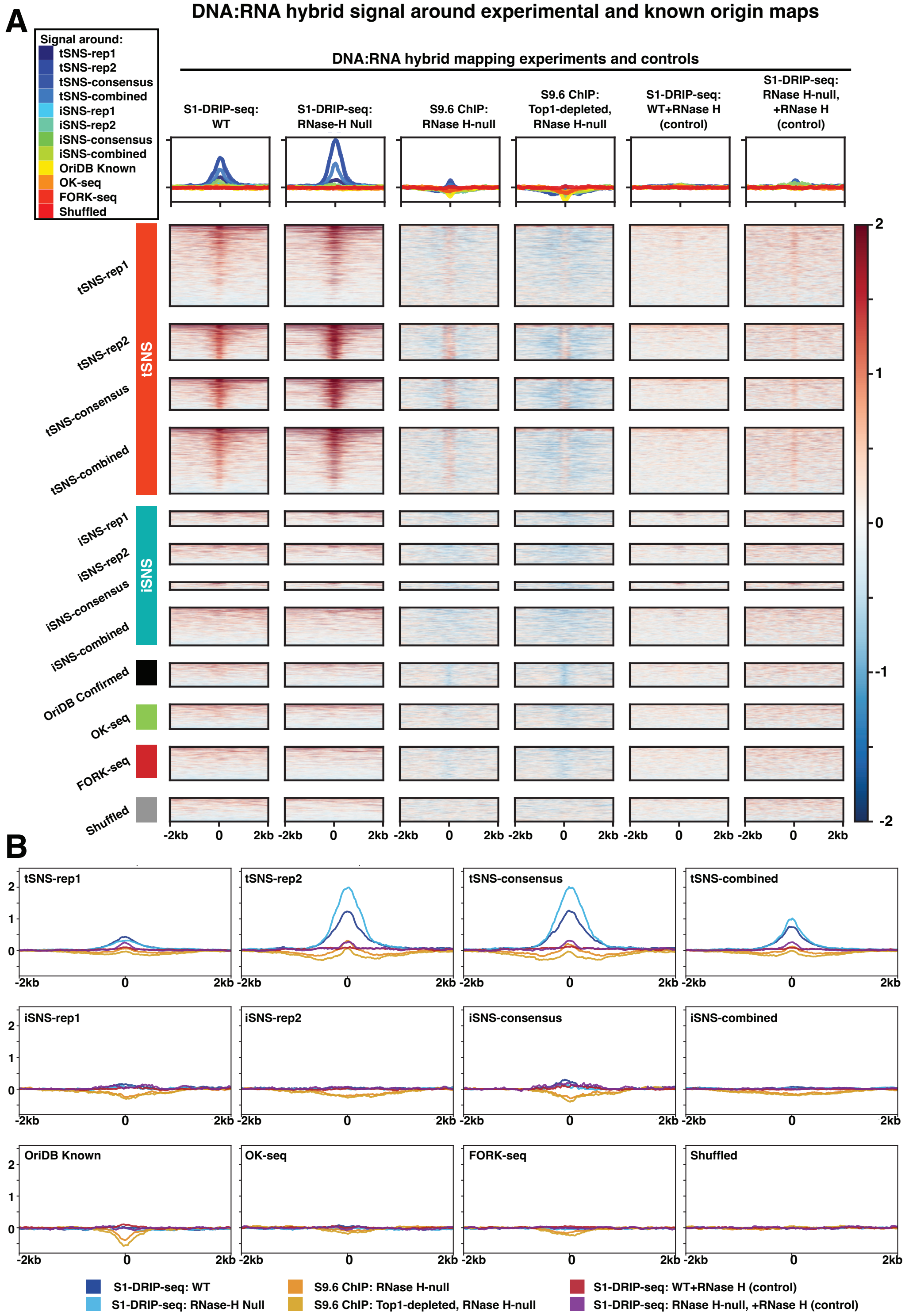


Fig. S20: $Log_2$ fold enrichment of RNA-DNA hybrid signal around experimental and known origin maps Both (A) and (B) show $log_2$ R-loop signal around tSNS-seq and iSNS-seq peak summits as well as confirmed known origins from OriDB, origin maps from OK-seq and FORK-seq, and randomly shuffled intervals. R-loop signal is enriched at tSNS summits, but flat or sometimes depleted at iSNS summits, known origins, OK-seq and FORK-seq origin maps, and randomly shuffled intervals. In contrast, R-loop control samples are relatively flat across all. Four different sets of peak summits were used for tSNS and iSNS to interrogate this effect: peak summits from replicates 1 and 2 as well as from peaks found in both replicates (consensus), and from peaks called when combining reads from both replicates (combined). (A) Shows heatmaps with summary profile lines across all regions above them. There is one set of heatmaps for each R-loop experimental and control dataset. The summary profile lines show the trend for that dataset over each set of "origin" locations. (B) Shows summary profile lines grouped by "origin" set rather than by which R-loop or control dataset the signal track is from. Each plot shows the signal from all R-loop and control datasets mapped around the given set of "origins".

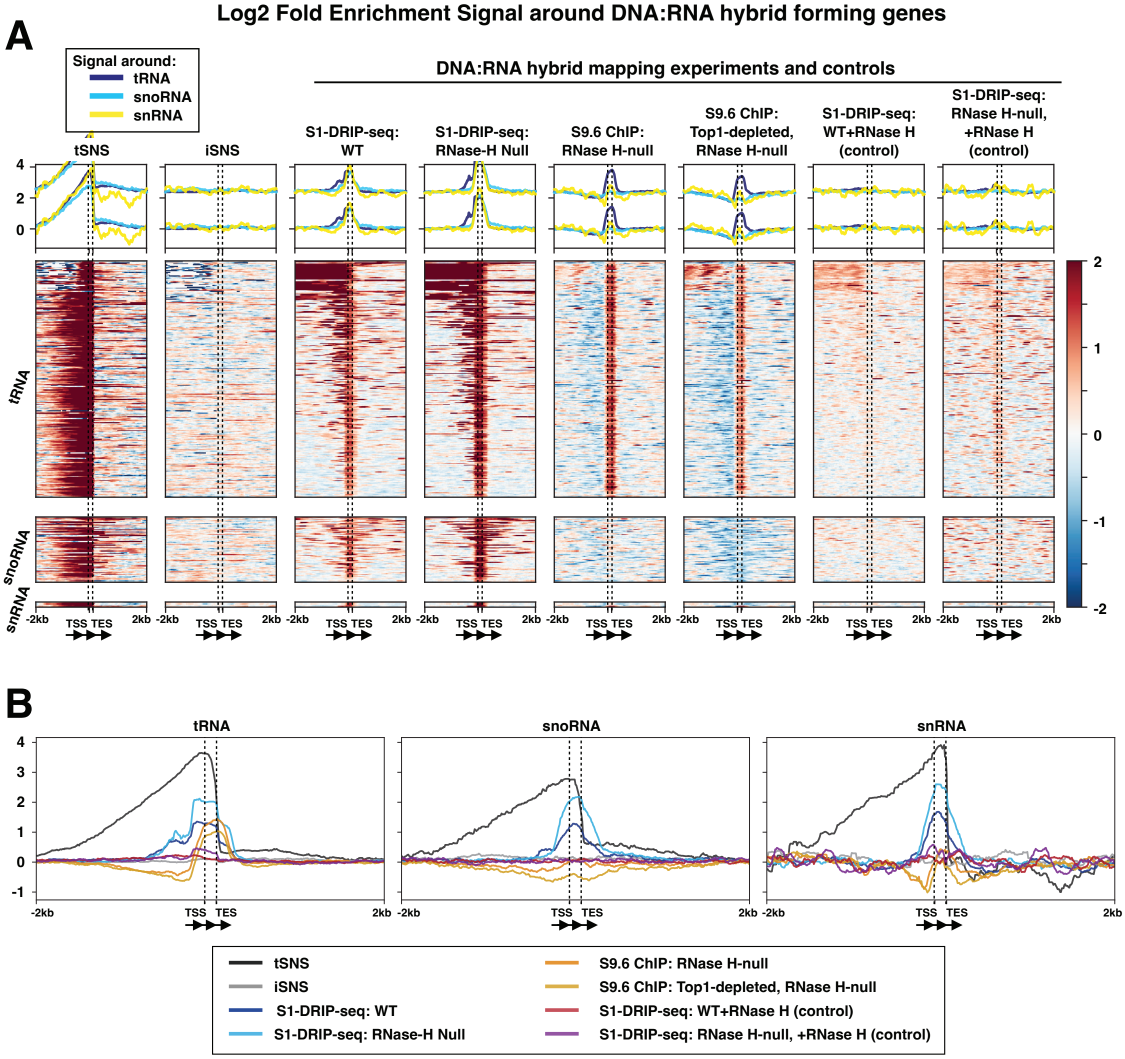


Fig. S21: $Log_2$ fold enrichment signal around RNA-DNA hybrid forming genes. Both (A) and (B) show $log_2$ signal from tSNS-seq, iSNS-seq, and various RNA-DNA hybrid (R-loop) mapping experiments and controls around tRNA, snoRNA, and snRNA genes in yeast. tSNS and the R-loop datasets are both enriched at these noncoding RNA loci whereas iSNS and the RNase-H controls are flat. (A) Shows heatmaps with summary profile lines across all regions above them. There is one set of heatmaps for each dataset. The summary profile lines show the trend for that dataset over each type of gene. (B) Shows summary profile lines grouped by gene type rather than dataset. Each plot shows the signal from all datasets mapped around the given type of gene.

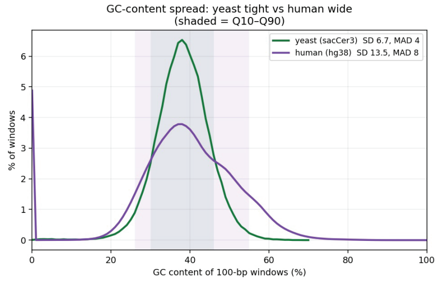


Fig. S22: Histogram showing the distribution of GC content calculated in 100 bp non-overlapping bins across the *Saccharomyces cerevisiae* (green) and *Homo sapiens* (purple) genomes, expressed as the percentage of all 100 bp bins falling into each GC% category. Shaded regions denote the 10th–90th percentile range for each species (yeast: 30–46%; human: 26–55%).
